# Retinitis pigmentosa associated mutations in mouse Prpf8 cause misexpression of circRNAs and degeneration of cerebellar granule neurons

**DOI:** 10.1101/2022.11.01.514674

**Authors:** Michaela Krausová, Michaela Kreplová, Poulami Banik, Jan Kubovčiak, Martin Modrák, Dagmar Zudová, Jiří Lindovský, Agnieszka Kubik-Zahorodna, Marcela Pálková, Michal Kolář, Jan Prochazka, Radislav Sedlacek, David Staněk

## Abstract

A subset of patients suffering from a familial retinitis pigmentosa (RP) carry mutations in several spliceosomal components including PRPF8 protein. Here, we established two novel alleles of murine *Prpf8* that genocopy or mimic aberrant PRPF8 found in RP patients - the substitution p.Tyr2334Asn and an extended protein variant p.Glu2331ValfsX15. Homozygous mice expressing either of the aberrant Prpf8 variants developed within first 2 months progressive atrophy of the cerebellum due to extensive granule neuron loss. Comparison of transcriptome from pre-degenerative and degenerative tissues revealed a subset of circRNAs that were deregulated in all tissues and both Prpf8-RP mouse strains. To identify potential risk factors that sensitize cerebellum for Prpf8 mutations we monitored expression of several splicing proteins during first eight weeks. We observed downregulation of all selected splicing proteins in wild-type cerebellum, which coincided with neurodegeneration onset. The decrease in splicing protein expression was further pronounced in mouse strains expressing mutated Prpf8. Collectively, we propose a model where physiological reduction of spliceosomal components during postnatal tissue maturation sensitizes cells to expression of aberrant Prpf8 and the subsequent deregulation of circRNAs triggers neuron death.

## Introduction

PRPF8 protein is the central scaffolding component of the spliceosome, which organizes its catalytic RNA core and directly regulates the activity of the key spliceosomal RNA helicase SNRNP200 (also known as Brr2) to control correct timing of spliceosome activation (Mozaffari-Jovin et al., 2014). Congenital mutations in genes encoding core spliceosome constituents including *PRPF8* (McKie et al., 2001), *SNRNP200* (Zhao et al., 2009), *PRPF3* (Chakarova et al., 2002), *PRPF4* (Chen et al., 2014), *PRPF6* (Tanackovic et al., 2011a), and *PRPF31* (Vithana et al., 2001) have been identified causative in a subset of familial blindness disorders known as non-syndromic retinitis pigmentosa (RP) (for review, see (Krausova and Stanek, 2018)). These dystrophies of retinal tissues feature dysfunction of retinal pigment epithelium (RPE) and gradual loss of both photoreceptor types. The pre-mRNA splicing factor genes are systemically expressed, the narrow phenotypic restriction observed with the human RP disease however indicates enhanced sensitivity of certain tissues towards expression of aberrant splicing factors.

Recent years have provided the first molecular insights into the pathophysiology of splicing factor forms of RP. The most studied is *PRPF31*, which is compromised by a spectrum of inactivating mutations (Audo et al., 2010; Huranova et al., 2009; Rio Frio et al., 2008; Sullivan et al., 2006; Vithana et al., 2003). RP mutations in *PRPF31* evoke mis-splicing of numerous retina specific genes including rhodopsin and ciliary and cellular adhesion genes, which disrupt the structural organization of RPE and photoreceptors (Buskin et al., 2018; Mordes et al., 2007; Ray et al., 2010; Yin et al., 2011; Yuan et al., 2005). The etiology of *PRPF8*-linked RP, comprises nearly two dozen of heterozygous genetic variants (De Erkenez et al., 2002; McKie et al., 2001; Ruzickova and Stanek, 2017) with predominance of missense mutations clustering to very C-terminus of PRPF8 that is responsible for the SNRNP200 modulation (Mozaffari-Jovin et al., 2013). Functional screens demonstrated that the majority of aberrant PRPF8 proteins exhibited altered ability to interact with spliceosome cofactors in human cell culture and impacted development including eye development in fly (Malinova et al., 2017; Stankovic et al., 2020). The only exception was the PRPF8 p.Tyr2334Asn substitution. This variant displayed in *in vitro* assays selective disruption of splicing efficiency and affected strongly pupation in Drosophila, but in contrast to the other pathological PRPF8 variants the PRPF8-Y2334N protein was properly incorporated into small nuclear ribonucleoprotein particles (snRNPs), key components of the spliceosome (Malinova et al., 2017; Stankovic et al., 2020). In addition, several RP-linked mutations in *PRPF8* target codons at the very 3’ end of the PRPF8 coding sequence causing shortening or aberrant prolongation of the PRPF8 C-terminus (De Erkenez et al., 2002; Martinez-Gimeno et al., 2003; McKie et al., 2001; Tiwari et al., 2016).

Several of the RP-causing PRPF8 missense mutations have been recapitulated in model organisms. Experimental animals homozygous for the aberrant substitutions Prpf8 p.His2309Pro (Graziotto et al., 2011) or p.Arg2310Gly (Kukhtar et al., 2020) were viable, ruling out a full Prpf8 loss-of-function effect since full genetic deficiency in *Prpf8* was embryonic lethal (Graziotto et al., 2011). However, although challenged to homozygosity the *Prpf8^H2309P/H2309P^* mice manifested mild, very late-onset retinal degeneration (Farkas et al., 2014; Graziotto et al., 2011). We are thus still lacking a robust vertebrate model that would clarify the disease-provoking mechanisms associated with distal C-tail-disrupted Prpf8 variants, and that would, moreover, elucidate principles underlying sensitivity of specific cell types to *PRPF8* RP-mutations.

Here, we genocopied the PRPF8 p.Tyr2334Asn missense mutation in a mouse because the PRPF8 p.Tyr2334Asn mutation confers severe clinical phenotype with early macular involvement and the molecular mechanism is unclear (Towns et al., 2010). In addition, we also generated an extended Prpf8 protein variant (*Prpf8^Δ17^*) that resembles human mutagenic frameshifts originating from PRPF8 C-terminal residues and a *Prpf8* knock-out allele that serves direct comparison with a presumptive *Prpf8* loss-of-function mode.

## Results

### Generation of the *Prpf8^Y2334N^* and *Prpf8^Δ17^* mice

To introduce the Prpf8 p.Tyr2334Asn mutation to the mouse C57BL/6N background, we supplemented CRISPR/Cas9 gene editing tools with DNA-based homology directed repair (HDR) templates (Fig. S1A and Table S1). In parallel to the correctly established *Prpf8^Y2334N^* allele in two founder animals, in a third individual the non-homologous end joining repair pathway gave rise to a *Prpf8* deletion allele, where a removal of terminal 17 base pairs (bp; nucleotides chr11:75,509,271-11:75,509,287; hereafter termed the *Prpf8^Δ17^* allele) within the Prpf8 open reading frame abolished five C-terminal amino acids including the stop codon. This allele thus encodes a Prpf8 protein variant with altered residues aa2331-2335 that is moreover extended by aberrant 9 amino acids at the C-terminus (Prpf8 p.Glu2331ValfsX15; called Prpf8 Δ17 in the following text; Fig. 1A). The *Prpf8^Δ17^* allele suitably resembles human PRPF8 RP variants where mutagenic frameshifts originate from the stop codon itself, or from mutations within or shortly upstream of the penultimate residue Tyr2334 and we therefore included this variant into further analysis (Martinez-Gimeno et al., 2003; Tiwari et al., 2016) (Fig. 1B). A detailed sequencing analysis of the targeted *Prpf8* regions in all three founder animals can be found in Fig. S1B.

**Figure 1.**
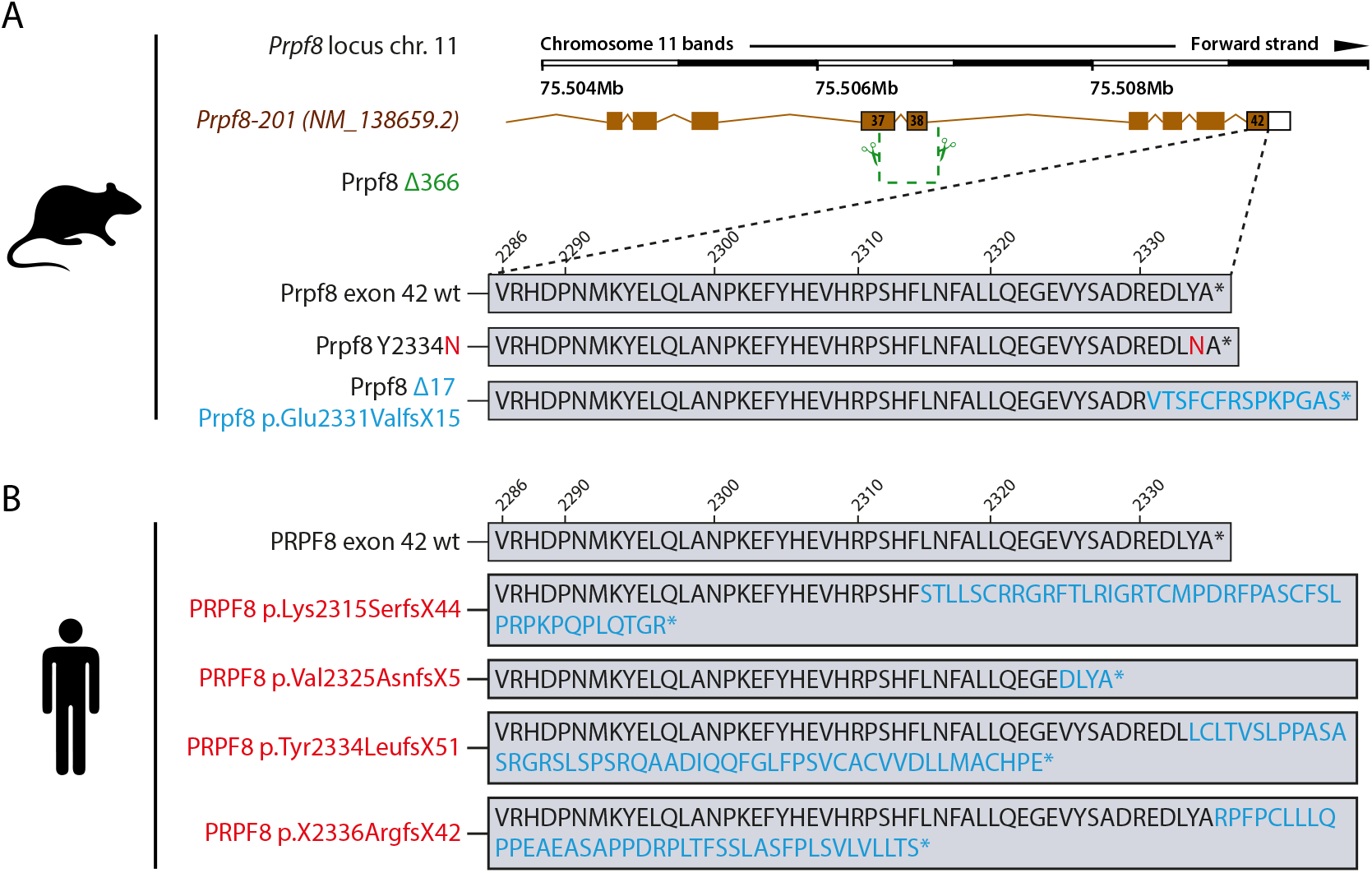
The newly established murine *Prpf8* alleles serve as a model for human RP-linked mutations. (*A*) Illustration of the three newly established *Prpf8* alleles: 1. The RP-causative substitution Prpf8 p.Tyr2334Asn (in red letters); 2. The *Prpf8^Δ17^* allele with an aberrantly prolonged Prpf8 ORF (in blue). 3. The *Prpf8^Δ366^* allele, where a large deletion of 366 bp eliminated the 3’ portion of *Prpf8* exon 37 until 5’ of intron 38-39 (in green). (*B*) The *Prpf8^Δ17^* allele serves as a genetic model for human RP-linked mutagenic frameshifts that map to the PRPF8 distal region (in red). See Table S1 and Fig. S1 for primer design and sequencing of founders and Fig. S2 for analysis of mutated offspring.

In previous *in vitro* experiments in human cells the PRPF8 Y2334N substitution did not alter the binding of PRPF8 with its key interactors (Malinova et al., 2017), the effect of the Prpf8 Δ17 mutation is however unclear. To inspect the binding profile of the aberrant Prpf8 variants, we performed immunoprecipitation of Prpf8 from cerebellar tissues and monitored co-precipitation of dedicated Prpf8 binding partners Prpf6 and Snrnp200 (Fig. S1C). Both mutated Prpf8 variants pulled down similar amounts of Prpf6 and Snrnp200 when compared to the wt protein, which indicated that neither of the mutations perturbed the Prpf8 interactions with these two U5 snRNP-specific factors.

All novel *Prpf8* strains were generated on default C57BL6/N genetic background given the higher efficiency of editing efforts we achieved with C57BL6/N-derived zygotes. However, the C57BL/6N lines carry the Crumbs homolog 1 (*Crb1) rd8* mutation that is known to progressively compromise photoreceptor integrity (Mattapallil et al., 2012). To avoid any effect of the *rd8* mutation, we bred the founders to C57BL/6J genetic background that does not carry the mutation at the *Crb1* locus. For phenotypic analyses, progeny was successively backcrossed to C57BL/6J for >7 generations when the contribution of the recurrent parent genome reaches >99% (Visscher, 1999). This approach also eliminated potential off target effects of CRISPR/Cas9 editing

### Homozygous *prpf8^Y2334N/Y2334N^* and *Prpf8^Δ17/Δ17^* mice develop cerebral neurodegeneration

Both newly established *Prpf8^Y2334N^* and *Prpf8^Δ17^* lines were crossbred to homozygosity. Pups were viable and born in Mendelian ratios, but we recorded a mild shortage of *Prpf8^Δ17/Δ17^* offspring and a surplus of *Prpf8^Y2334N/Y2334N^* progeny (Fig. S2A-B) at the expense of heterozygous animals. Homozygous animals of both strains were moreover slightly smaller compared to their heterozygous and wt littermates of matched gender. In *Prpf8^Δ17/Δ17^* mice, the difference in body weight was already noticeable at 4 weeks (lower by 13% and 12% in males and females, respectively), and persisted to later ages (15% reduction in males and 12% in females at 8 weeks) (Fig. S2C). The *Prpf8^Y2334N/Y2334N^* animals showed a smaller weight disproportion (6% decline in males and 8% in females at 12 weeks of age) (Fig. S2C).

When reaching approximately 15 weeks of age, homozygous *Prpf8^Y2334N/Y2334N^* and *Prpf8^Δ17/Δ17^* animals exhibited tremor indicating a potential neurodegeneration. To investigate the onset and progression of the suspected neurodegenerative changes, we histologically examined cerebellar structures at 4, 6, 17/15, and 22 weeks of age (Fig. S3 for *Prpf8^Y2334N/Y2334N^* animals and Fig. S4 for *Prpf8^Δ17/Δ17^* mice). The postnatal maturation of the cerebellar tissue was normally completed, however at week 6 we observed reduction in cellularity in the granule cell layer in the posterior lobe (Figs. S3A-B and S4A-B). In aging mice, progressive thinning in the stratum granulosum roughly followed the anatomical subdivision of the cerebellar tissue and overall, hemispheric segments of lobules seemed more strongly afflicted compared to vermian regions (Fig. S3C). The granular layer content was severely decreased by 17 and 15 weeks of age in *Prpf8^Y2334N/Y2334N^* and *Prpf8^Δ17/Δ17^* animals, respectively (Figs. S3D and S4C), which coincided with the onset of tremor and locomotion disturbances. By 22 weeks of age, the granule cell layer almost diminished in the whole cerebellum (Figs. S3E and S4D), and mice had to be euthanized. Heterozygous peers of *prpf8^Y2334N/wt^* and *Prpf8^Δ17/wt^* genotypes did not develop any gross pathological changes by the ultimate timepoint examined (Figs. S3E and S4D).

In the latest, 22 weeks-cohorts of both Prpf8 lines, we histologically inspected specimen from spinal cord, peripheral nerves, skeletal muscle and eye, but did not observe any pathological changes (Fig. S5). In the aged *Prpf8^Δ17/Δ17^* animals we also examined the extracerebellar populations of granule neurons that reside in the olfactory bulb and dentate gyrus, but no apparent perturbances were observed (Fig. S6A-C). To survey the condition of retinal structures, both eyes of the *Prpf8^Δ17^* 22 weeks-cohorts were screened by paralleled OCT and ERG examinations. The ophthalmic inspection of fundi did not reveal any abnormalities in retinal layering, nor in the size and placement of the head of the optic nerve, and the structure and distribution of superficial blood vessels as well did not vary from wt controls. The retinal layers were well developed, however in homozygous *Prpf8^Δ17/Δ17^* animals we recorded a slight, but significant reduction in retinal thickness (Fig. S6D-E). In the electroretinographic examinations, all mice showed scotopic and photopic responses with normal waveform shape preserved. Despite a large variability among animals, in homozygous mutants the responses were slightly smaller in amplitude and delayed in time when compared to *Prpf8^wt/wt^* and *Prpf8^Δ17/wt^* counterparts. This effect seemed to be more pronounced in the wave b rather than wave a, suggesting that the photoreceptor function was affected to a lesser extent than post-photoreceptor retinal processing (Fig. S7). The subsequent histopathological inspection of ocular tissues nonetheless did not reveal any apparent retinopathy (Fig. S6F). Taken together, the cerebellar cortex is likely the primary site perturbed by the presence of both aberrant Prpf8 variants.

### Prpf8 mutations induce changes in cerebral RNA landscape

To investigate how the mutations in splicing factor Prpf8 affect the transcriptional landscape of the cerebellum, total RNA was isolated from cerebella of 12-weeks-old mice of the *Prpf8^Y2334N^* strain and from 8-weeks-old *Prpf8^Δ17^* animals, and subjected to next-generation RNA sequencing (RNA-Seq). These timepoints were selected to precede the manifestation of tremor and locomotion disturbances, and represented two consecutive stages of the granular layer degeneration. In more detail, at the earlier timepoint of 8 weeks the *Prpf8^Δ17/Δ17^* animals displayed apoptosis occurring in the granular layer in apical portions of the posterior lobe, while the vermian regions as well as the anterior cerebellum segment were less strongly afflicted (Fig. 2A and C). The more advanced stage of the *Prpf8^Y2334N/Y2334N^* mice was, in comparison, characterized by noticeable reduction of granule neuron density in the posterior apices that suggested preceding clearance of the deceased neurons and residual debris by phagocytic microglia (see below). Neuronal death has by then proceeded to lobule bases and was present in the anterior compartment (Fig. 2B-D).

**Figure 2.**
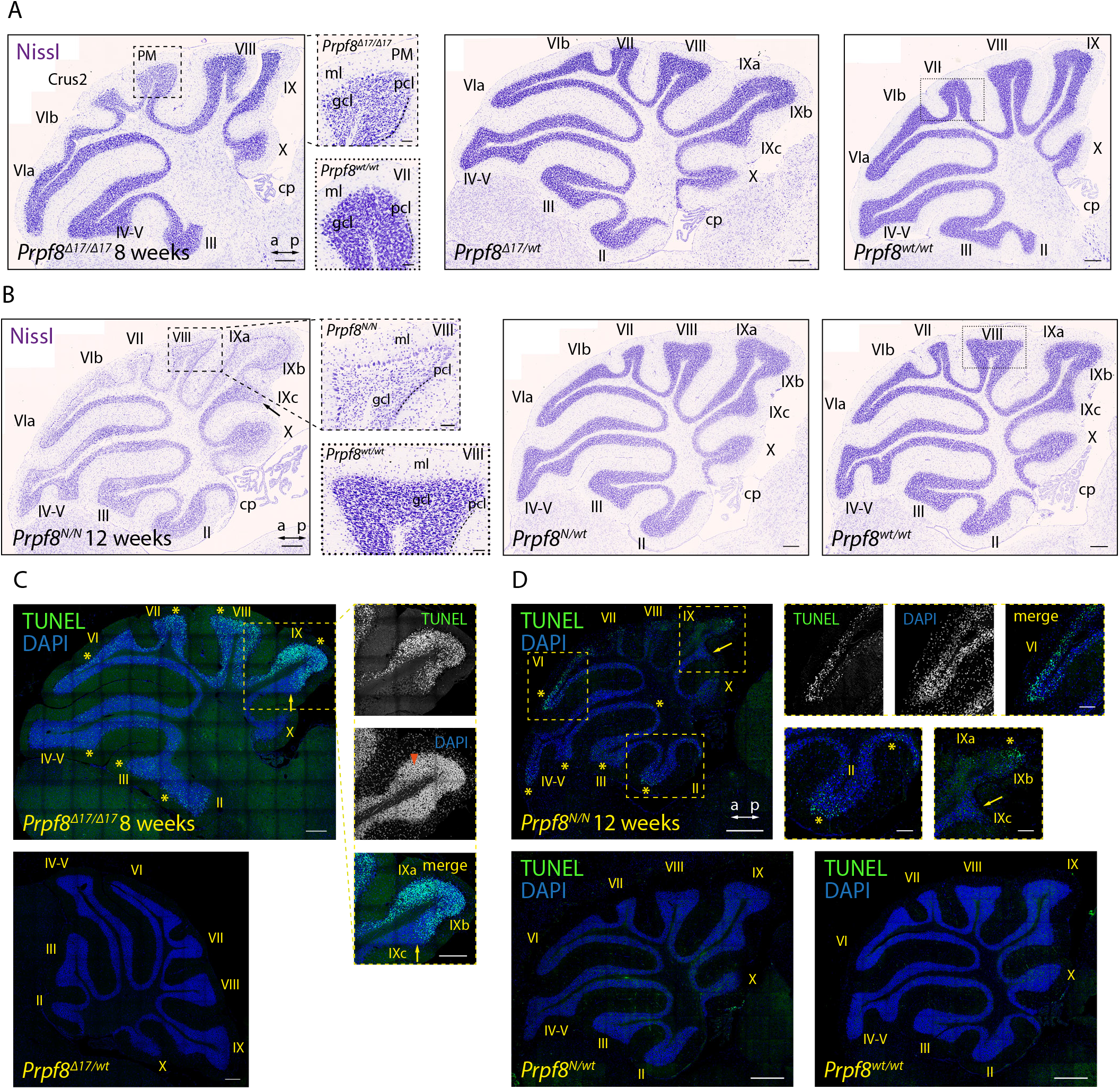
Aberrant Prpf8 variants provoke degeneration of the cerebellar granule cell layer. Histopathological analysis and TUNEL assay on cerebellar specimen collected from 8 weeks old *Prpf8^Δ17^* animals and from 12 weeks old *Prpf8^Y2334N^* mice. (*A-B*) NissI staining on sagittal cerebellar sections acquired from *Prpf8^Δ17^* (*A*) and *Prpf8^Y2334N^* (*B*) individuals. (*A*) In 8 weeks old *Prpf8^Δ17/Δ17^* mice we detected shrunken granule neurons in apical portions of the posterior lobe (the detail is from paramedian lobule (PM)). (*B*) In 12-weeks-old homozygous *Prpf8^Y2334N/Y2334N^* animals we observed substantial reduction in granule cell density in apices of the posterior lobe (detail is from lobule VIII (dashed rectangle)). The loss of the granule neurons followed the anatomical subdivision of the cerebellum (note the sharp boundary between posterior lobule IXb and lobule IVc that belongs to the nodular lobe (black arrow)). a - anterior, cp - choroid plexus, Crus2 - crus 2 of the ansiform lobule, gcl - granule cell layer, ml - molecular layer, p - posterior, pcl - Purkinje cell layer. Scale bar set to 200 μm, in details 50 μm. For additional time points and tissues see Figs. S3-S7. (*C-D*) Visualization of apoptotic cells by TUNEL assay. (*C*) In 8-weeks-old *Prpf8^Δ17/Δ17^* cerebella, neuronal death was present in apices of the posterior lobe, with numerous shrunken granule cells detected (orange arrowhead). Note the strict apoptotic boundary between lobule IXb and IXc (yellow arrow). Scale bar uniformly set to 200 μm. (*D*) In 12-weeks-old *Prpf8^Y2334N/Y2334N^* animals apoptosis was prevalent in the apex of lobule VI and was also present in the granular layer in both apical and base regions of the anterior lobe (yellow asterisks). The apoptosis followed anatomical division of the cerebellum, for now sparing granule neurons in lobule IXc that belongs to the nodular lobe (yellow arrow). No apoptotic signal was recorded in heterozygous *Prpf8^Y2334N/wt^* animals (bottom row). Scale bar = 200 μm, in insets 50 μm. The numbers of animals in individual cohorts can be found in the Methods section.

The RNA-seq analysis revealed significant changes in gene expression and hundreds of genes were up- and down-regulated in both Prpf8 mutant lines but not in the heterozygote animals (Fig. 3A and Table S2). To explore the likely fate of individual cerebellar cell types, a set of classificatory marker genes was extracted from a cerebellar single-cell RNA-Seq profiling study (Saunders et al., 2018) as a proxy that delimited homeostatic cerebellar populations (Fig. 3B). In concordance with the prior histopathological findings, we observed in homozygous animals a decline in expression of well-established granule cells markers including *Gabra6, Rbfox3/NeuN* and *Reln*. We also detected preferential decline of genes physiologically enriched in the granular layer of posterior cerebellum such as paired box 6 (*Pax6*), and T-cell leukemia, homeobox 3 (*Tlx3*) (Divya et al., 2016) (Fig. 3C), which was consistent with observed onset of degeneration in the posterior region (Fig. 2).

**Figure 3.**
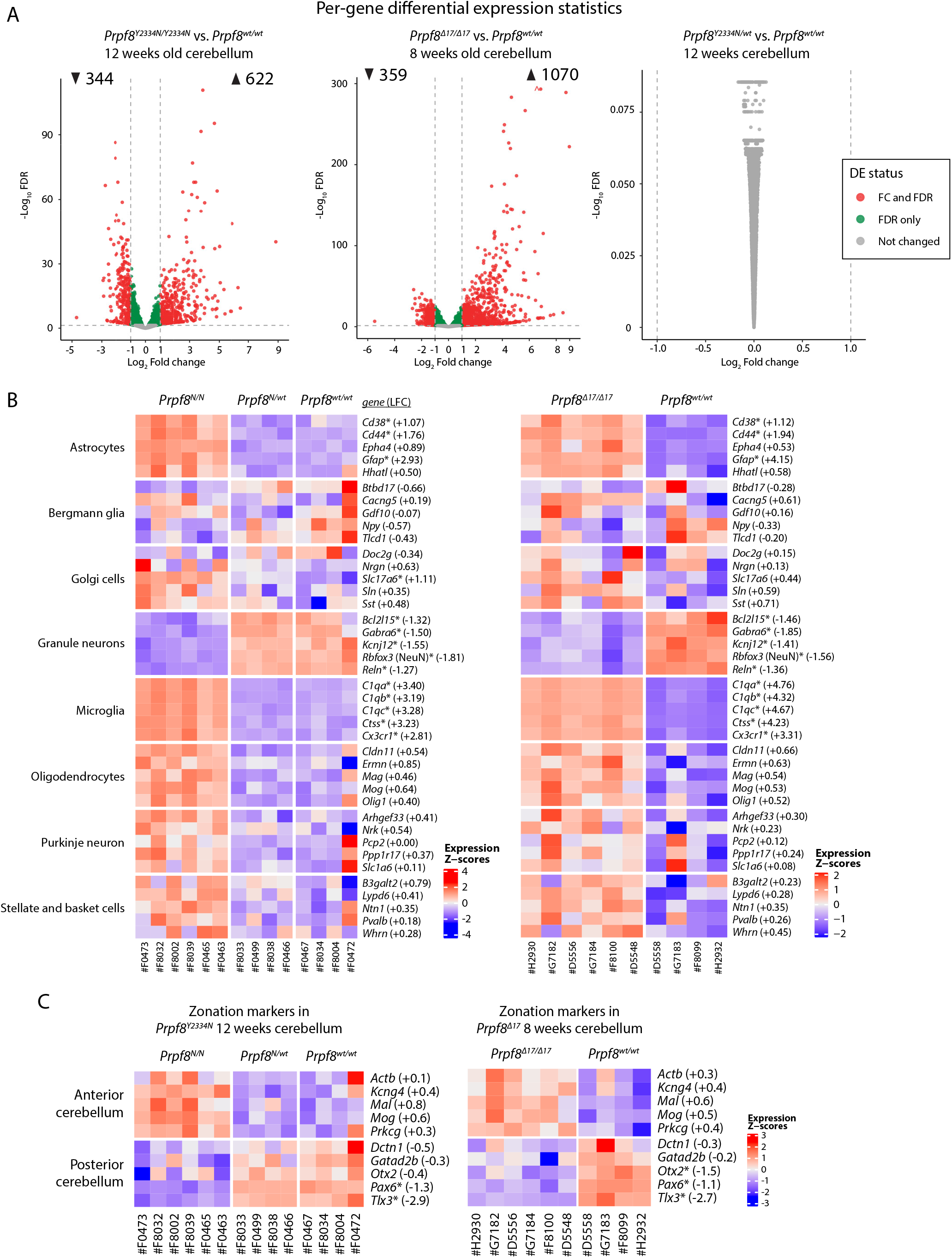
Differential gene expression in Prpf8-mutant cerebella. RNA was isolated from cerebellar samples collected of 12 weeks old animals of the *Prpf8^Y2334N^* strain, or 8-weeks-old *Prpf8^Δ17/Δ17^* animals and their *Prpf8^wt/wt^* littermates, and analyzed by RNA-Seq. (*A*) Volcano plot depicting -log10 False Discovery Rate (FDR) versus log_2_ Fold change difference in gene expression between Prpf8 homozygous mutant mice and *Prpf8^wt/wt^* controls. Red circles denote significantly up- and downregulated genes (FDR<0.05, foldchange cutoff is set at 1; see also Table S2). n=6 for *Prpf8^Δ17/Δ17^* and *Prpf8^Y2334N/Y2334N^* biological replicates and n=4 for *Prpf8^wt/wt^* litter-matched controls (both strains). (*B-C*) Heatmap representation of gene expression levels of selected marker genes representing major cerebellar homeostatic cell types (*B*) and anterior/posterior parts of cerebellum (*C*). Values were standardized to display the same range of expression values for each gene. * - differentially expressed genes defined by |Log_2_ Fold change|>1 and FDR<0.05; the estimated Log_2_ Fold change value (LFC) is provided for individual genes and represents the difference between homozygous mutant and *Prpf8^wt/wt^* cohorts. Expression of selected marker genes was validated by RT-qPCR and can be found in Fig. S8C.

Concerning the synapse organization, we recorded a drop in prominent scaffolding proteins of the active zone cytomatrix including *Bsn* and *Erc1* as well as decrease in multiple small GTPase effectors that play essential role in neurotransmitter release (*Rims1, Rims2*, and *Rims3*) (Fig. S8A,C). Homozygous mutant cerebella displayed also reduction genes involved in the Wnt signaling cascade (*Wnt7a, Dvl1 and Fzd7*), but not *Fzd5*), which indicated perturbance of the Wnt7a signaling axis present at the dendritic, post-synaptic side of glomerular rosettes (Ahmad-Annuar et al., 2006). Altogether, the spectrum of the downregulated genes suggested loss of components specific to axonal as well as dendritic synaptic zones of granule cells.

The category of upregulated genes was enriched with multitude of microglial markers, that indicated presence of microglia at various stages of activation (Figs. 3B and S8B,C). We also observed a slight rise in transcripts characteristic for oligodendrocytes, such as *Mag* and *Mog*, which can originate in the physiologic enrichment of myelination factors in the anterior cerebellum (Divya et al., 2016), and may reflect the disproportion of representation of specialized cerebellar cell types in the degenerating tissue. Similarly, the small increase in Purkinje neuron-specific genes might be linked to their natural overrepresentation in the anterior lobe and can reflect the regional variation of Purkinje cell populations within cerebellum.

To confirm the expression profiling we detected selected proteins using immunocytochemistry in cerebellar sections of 8- and 12-week-old animals (Fig. S9). In accordance with RNA-seq results we observed downregulation of granule cell marker Rbfox3 (NeuN) and synaptic marker PSD95 (encoded by *Dlg4*) in wider pre-apoptotic areas, that might hallmark distressed neurons and precede the upcoming cell death (Fig. S9A-B). Consistent with RNA-seq data, the granule cell layer was also strongly positive for astrocytes (Fig. S9C) and CD45^+^ microglia (Fig. S9D), We did not observe any major alterations affecting Purkinje neurons and Bergmann glia (Fig. S9E-F). The blend of histopathological and transcriptomic analyses revealed substantial changes occurring in the cerebellar tissue of homozygous Prpf8-mutant animal, namely the degeneration of granule neurons in the posterior lobe was accompanied by extensive induction of reactive microglia and astrogliosis.

### Deregulation circular RNAs expression in the cerebellum of homozygous *prpfg^Y2334N/Y2334N^* and *Prpf8^Δ17/Δ17^* mice

We further analyzed RNA-Seq data for changes in alternative splicing (Table S3), but given the distorted representation of cerebellar cell subtypes it was not feasible to attribute the recorded differences to Prpf8 mutations only. The analysis of alternative splicing nonetheless revealed altered usage of exons, which were previously shown to be included in circular RNAs (Rybak-Wolf et al., 2015). A detailed analysis focused on the circRNA expression revealed that the cerebella of mutant mice exhibited a considerable deregulation of the circular transcriptome (Fig. 4 and Table S4). In detail, we observed 161 (*Prpf8^Δ17/Δ17^*) and 26 (*Prpf8^Y2334N/Y2334N^*) circRNAs more than twice elevated and on the other hand we recorded more than a twofold decline in 63 (*Prpf8^Δ17/Δ17^*) and 34 (*Prpf8^Y2334N/Y2334N^*) circRNAs (Fig. 4A). Interestingly, in both strains the majority of significantly deregulated circRNAs were altered by a larger factor compared to the shift in host gene abundance, and there were also classes of circRNAs that followed an inverse scenario compared to change in their parental mRNA (Fig. 4B). Several abundant up- and down-regulated circRNA were selected (Table S5) and changes in expression confirmed by RT-qPCR (Fig. 4C).

**Figure 4.**
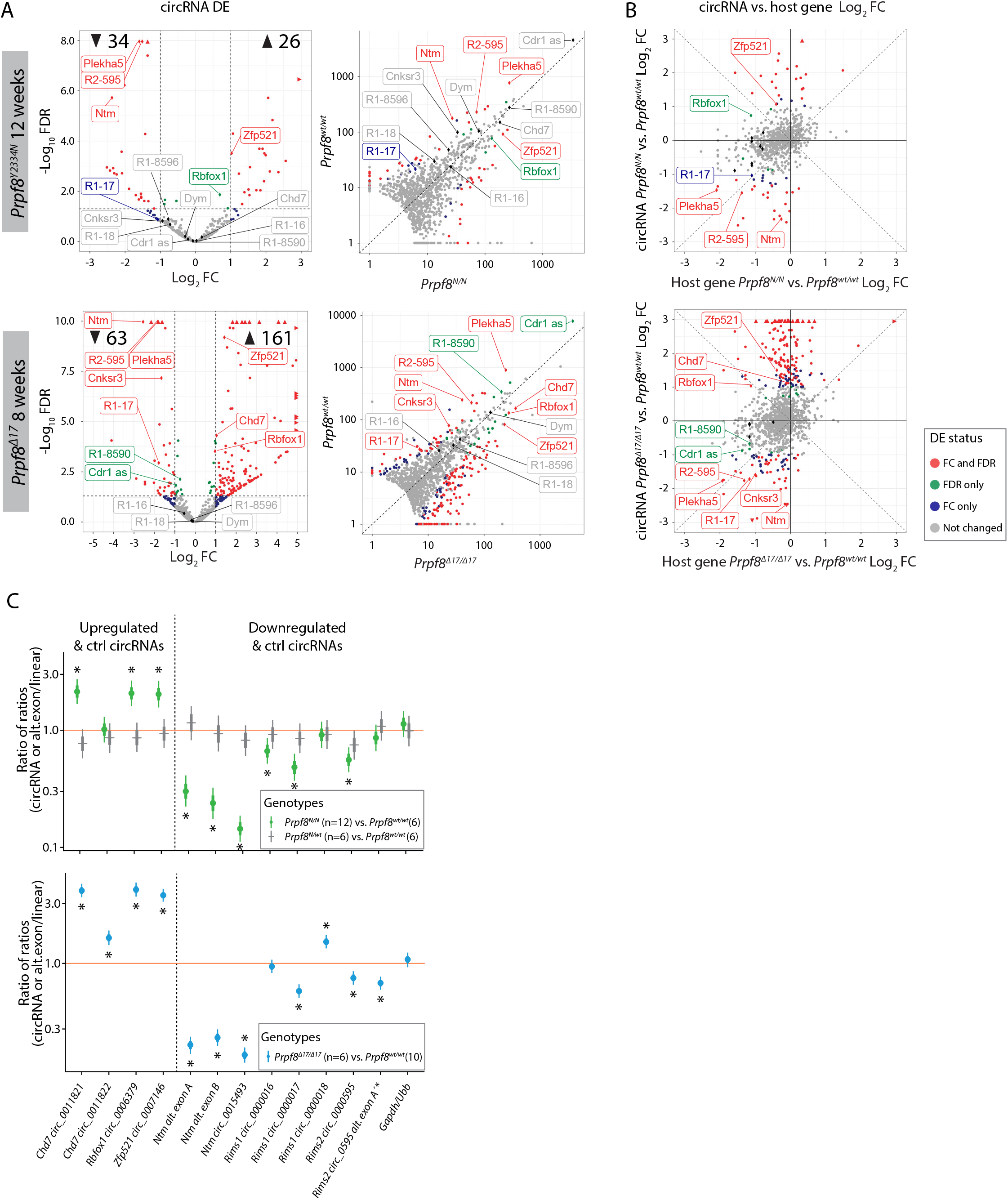
Deregulation of circular RNAs in cerebellar tissue of homozygous Prpf8 mutant mice. (*A*) Volcano plot showing -log10 (FDR) versus log_2_ fold difference in circular RNA expression between 12-weeks-old *Prpf8^Y2334N/Y2334N^* animals (top row), or 8-weeks-old *Prpf8^Δ17/Δ17^* animals and their *Prpf8^wt/wt^* littermates (second row). Scatterplot comparing library-normalized abundance of individual circRNAs (in BSJ read counts) between homozygous mutant mice (on the X-axis) and wt controls (Y-axis). Data accompanying all plots can be found in Table S4. (*B*) Comparison of Log_2_ fold change in circRNAs expression and the corresponding fold change of the linear transcripts expression from the corresponding parental gene in homozygous mutant mice and wt controls. DE-differential expression, FC-fold change, LFC-Log fold change, *R1-Rims1, R2-Rims2*. Assorted circRNAs are highlighted by diamond symbols and their specifications are provided in Table S5. Triangles represent circRNAs with values reaching beyond the displayed scale. Only circRNAs with mean normalized BSJ read counts minimum of 3 are shown in all plots. (*C*) Statistical model-based estimates of relative expression of alternative exons, and circRNAs and their corresponding linear transcripts in cerebellar samples collected from 12-weeks-old animals from the *Prpf8^Y2334N^* strain and 8-weeks-old *Prpf8^Δ17/Δ17^mice*. The estimates are based on a linear mixed model of the RT-qPCR data. Posterior distribution of ratio of each product (circRNA or *Ntm* alternative exons; horizontal axis) to a corresponding canonical product (linear mRNA transcript) was computed for each genotype. Estimates of the ratio of those ratios across genotypes are shown (vertical axis, log scale). Values below 1 indicate a lower proportion of the circRNAs or decreased inclusion of the alternative exons, respectively, in the mutant animals compared to wt counterparts. Shown are posterior credible intervals (lines: 95% - thin, 50% - thick), and means (points). Asterisks indicate comparisons for which the 95% credible interval excludes 1 (no difference). The comparison “Rims2 circ_0595 alt. exon A’*” measures inclusion of alternative exon A’ into the *Rims2* circ_0000595.

The disequilibrium in the cerebellar circRNAs in the older mice might have been influenced by the altered representation of individual cell types in the degenerating tissue. To investigate whether cerebellar circRNAs are perturbed already prior to the onset of the granule cells decay, we examined the cerebellar transcriptome by RNA-Seq approach in 4 weeks old animals. At this timepoint we did not detect any pathological changes by immunohistochemistry, TUNEL assays, and RT-qPCR approaches (Fig. S10). The RNA-seq profiling revealed no substantial differences in expression of linear RNA forms between wt and homozygous mice of either strain (Fig. S11A) and we did not record any significant deregulation of circRNAs either (Fig. S11B). However, a more detailed analysis of circRNA expression in the 4 weeks old animals by RT-qPCR revealed that expression of several circRNAs was already shifted in the same direction as in the degenerated cerebellum. We observed a decline of circ_0015493 produced from *Ntm* and *Rims1* circ_0000017 and *Rims2* circ_0000595 derived from *Rims1* and *Rims2* synaptic genes, respectively (Figs. 5A-B and S11C). *Rims1* & *Rims2*-derived circRNAs are highly abundant in the cerebellum and *Rims2* circ_0000595 represented the dominant form of total *Rims2* transcripts in 4 weeks as well as older animals (Fig. S11C and Table S4) (Rybak-Wolf et al., 2015). Importantly, we observed a significant drop in *Rims1* circ_0000017 amounts, while other cognate circRNAs formed from the same locus, such as *Rims1* circ_0000016 and *Rims1* circ_0000018 were changed by a lesser extent (Fig. 5A-B). In addition, we observed a significant decline in *Ntm* circ_0015493 in the pre-degenerative cerebella of both Prpf8 mutant strains (Figs. 5A-B and S11D) but not the cognate linear mRNA (Fig. S12). Similarly to *Rims2* circ_0000595, the *Ntm* circRNA entity represented the prevailing form of total cerebellar *Ntm* content (Table S4). We further analyze circRNA-specific exons that were only marginally included into linear mRNA (*Ntm* and *Rims1*) or in the case of the *Rims2* gene were fully circRNA specific (Fig. S11C-D and Table S6) (Rybak-Wolf et al., 2015). Both transcriptome-wide analysis and RT-qPCR in 4-week old animals revealed lower inclusion of these circRNA-specific exons, which further documented the lower expression of the cognate circRNAs (Table S6 and Fig. 5A-B).

**Figure 5.**
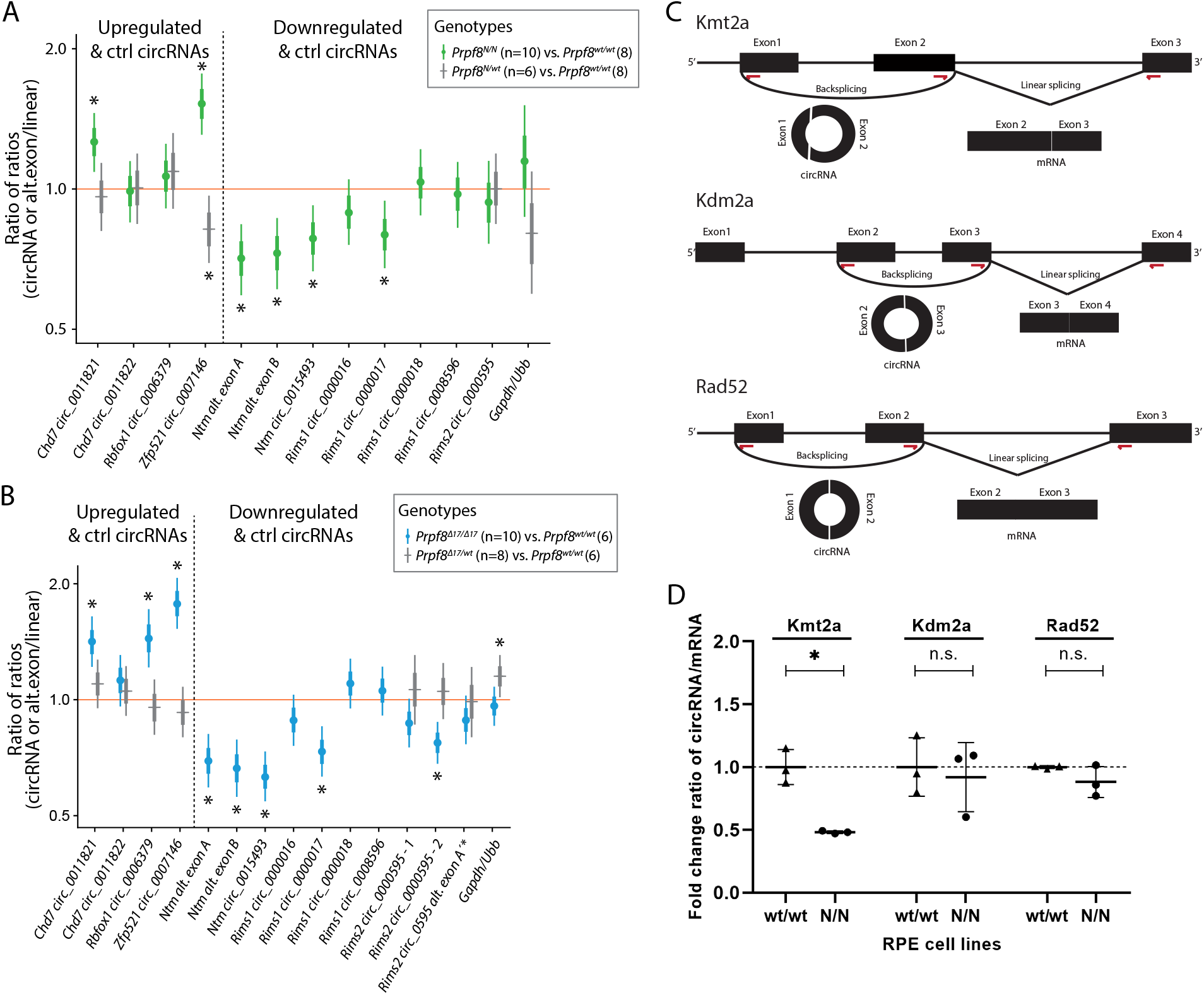
Differential circRNA expression in 4 weeks old cerebella of both Prpf8 mutant strains and cells expressing PRPF8^Y2334N^. (*A-B*) Statistical model-based estimates (see Fig. 4 for details) of relative expression of circRNAs and their corresponding linear transcripts in cerebellar samples gained from 4 weeks old animals from PRP*Prpf8^Y2334N^* strain (*A*), or 4 weeks old *Prpf8^Δ17^* mice (*B*). Shown are posterior credible intervals (lines: 95% - thin, 50% - thick), and means (points). Asterisks indicate comparisons for which the 95% credible interval excludes 1 (no difference). (*C*) Schematic representation of circRNA producing reporters. Red arrows indicate positions of primers used for detection of circular and linear forms. (*D*) CircRNA reporters were transfected into RPE expressing GFP-PRPF8^WT^ (wt/wt) or GFP-PRPF8^Y2334N^ (N/N) from both alleles and circRNA:linearRNA ratio was determined by RT-qPCR. N=3, statistical significance was determined by paired t-test. Expression of mRNAs and circRNA based on RNA-seq data is presented in Fig. S11A,B and Tables S6-S9.

Analyses of splice site (ss) strength scores collectively identified a weak 3’-ss rating for *Rims1* alternative exons A and B, for *Rims2* exon 22, and for *Ntm* alternative exon B” (Table S7). This data indicate that the suboptimal 3’-ss that surround these alternative exons are likely not properly recognized by the mutated variants of Prpf8, and this finding might explain why only some circRNA species originating from the same host gene portion are selectively affected.

In contrast to circRNA downregulation, several circRNAs (but the not linear mRNAs) were upregulated in the pre-degenerative stage (Figs. 5A-B and S12A), and this elevation persisted to later stages in both Prpf8 mutant strains (Fig. 4C). In detail, we examined the expression of *Rbfox1* mmu_circ_0006379, *Zfp521* mmu_circ_0007146, and *Chd7* mmu_circ_0011821 (Fig. 5A-B), because their parental genes are physiologically enriched in granule and Golgi cells. The linear-to-circular ration was distorted in favor of the circRNA form (Fig. 5A-B), however parallel quantification by RT-qPCR did not suggest that the circularization was carried out at the expense of linear splicing (Fig. S12). Interestingly, in advanced stages of cerebellar neurodegeneration the expression of *Rbfox1, Zfp521*, and *Chd7* host mRNAs was substantially decreased (Fig. S12), in contrast to the surplus of the circRNA forms (Fig. 4C).

To further test a role of Prpf8 mutation on circ RNA expression we established three reporters derived from mouse gene loci producing circRNAs (Fig. 5C) and expressed them in human RPE cell line containing the Y2334N mutation in both *PRPF8* alleles (N/N). Comparing the expression of circular to linear RNA form, we detected significant reduction of circRNA generated from the Kmt2a reporter in RPE^N/N^ cell line (Fig. 5D). Consistently, circRNA expressed from the *Kmt2a* (ENSMUSG00000002028) gene was downregulated in mutated animals (Table S4). These data further show that mutations in Prpf8 protein affect expression of particular circRNAs.

To investigate a potential contribution of mis-regulated splicing of protein coding genes, we analyzed splicing of linear mRNAs in RNA-Seq runs from 4 weeks old animals. Deregulated splicing involved intron retention, exon skipping, and alternative 5’-/3’-splice site recognition. Only events with ≥15% difference between the mutant mice and wt controls were considered biologically relevant (Fig. S13 and Table S8). In the *Prpf8^Δ17/Δ17^* animals we observed large numbers of intron retention events and 25 of those genes overlapped between *Prpf8^Δ17/Δ17^* and *Prpf8^Y2334N/Y2334N^* animals. A more detailed analysis revealed that the same intron was targeted in only one gene, CDH18, but the effect of Prpf8 mutation was opposite, meaning that the targeted intron was better spliced in *Prpf8^Δ17/Δ17^* animals while less efficiently spliced in *Prpf8^Y2334N/Y2334N^* mice. A comparison of exon skipping and alternative 5’/3’ splice site usage did not reveal any overlap between the two mouse strains (Table S8). We further subjected the differentially spliced genes to gene set enrichment analysis (GSEA). However, we did not identify any overlapping categories between the two mutant strains (Table S9).

Altogether, animals carrying aberrant variants of splicing factor Prpf8 displayed distorted splicing and expression of several circRNAs that are presumably granule neuron-specific, and these RNA processing defects preceded the onset of pathological changes affecting the cerebellar tissue.

### Cerebral aging is associated with downregulation of splicing proteins including Prpf8

The expression of spliceosomal components has been previously shown to vary in different developmental stages and on the course to full adulthood (Cao et al., 2011). Here, we took advantage of our mouse models and test the hypothesis that lower expression of Prpf8 correlates with onset of the neurodegeneration. To examine the abundance of Prpf8 during and after cerebellar postnatal maturation, we monitored the levels of Prpf8 and three other splicing factors Prpf6, Prpf31 and Snrnp200 in the cerebellum, retina and control liver snips (Fig. 6). We harvested samples from the organs in postnatal week 1, 2, 4 and 8, and observed a significant decline in all tested splicing components in the cerebellum from week 4 onwards. A similar reduction of splicing protein expression was observed in retina but here, the expression of Prfp8 declined to 41% while in cerebellum Prpf8 protein levels dropped more than 5 times to 14% of the amount observed in the first week. Downregulation of three of four tested spliceosomal components (Prpf8, Prpf6 and Prpf31) was highest in aging cerebellum followed by retina samples. In livers, only Prpf31 protein declined below 50% at week 8. The downregulation of splicing factors was even more pronounced in animals expressing the RP variants, which indicates that RP mutations, and namely the Y2334N substitution, negatively affect stability of splicing proteins and might contribute to the neurodegeneration phenotype (Fig. 7).

**Figure 6.**
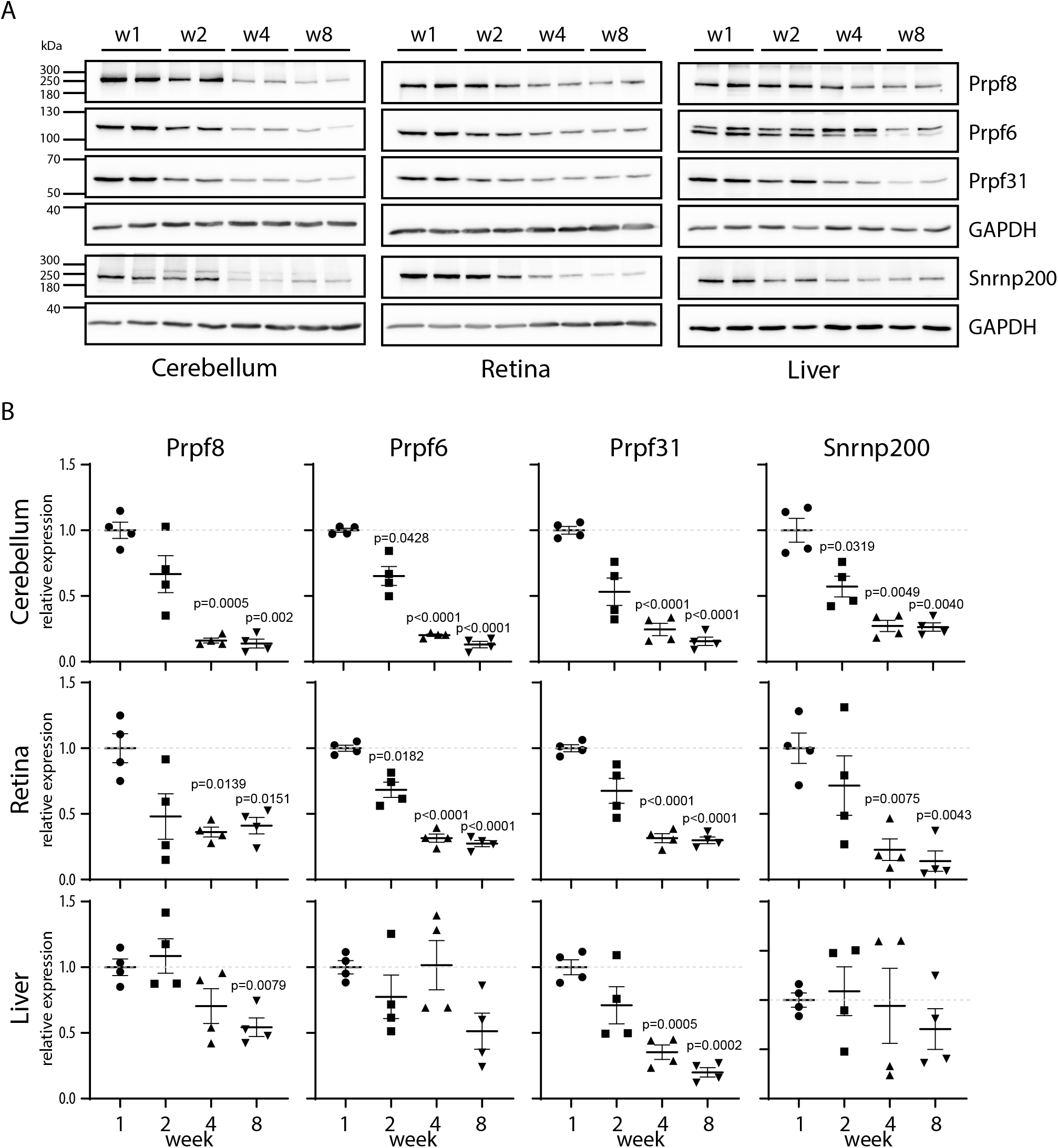
Expression of snRNP proteins is reduced in mature cerebellum. (*A*) Expression of snRNP-specific proteins was assayed in cerebella, retina and liver of wild-type animals at 1, 2, 4, and 8 weeks by Western blotting. Samples from two animals were analyzed per each timepoint. (*B*) Statistical analysis of protein expression at 1, 2, 4, and 8 weeks. Protein abundance in individual animals was first normalized to levels of GAPDH is the same tissue and then calculated as % of average value of the value at the first week animals. N=4. Statistical significance of differential protein abundance in the given genotype groups was examined by one-way ANOVA followed by Dunnett’s T13 test to find samples that significantly differ from expression in week 1 (mean with SEM are displayed together with p values).

**Figure 7.**
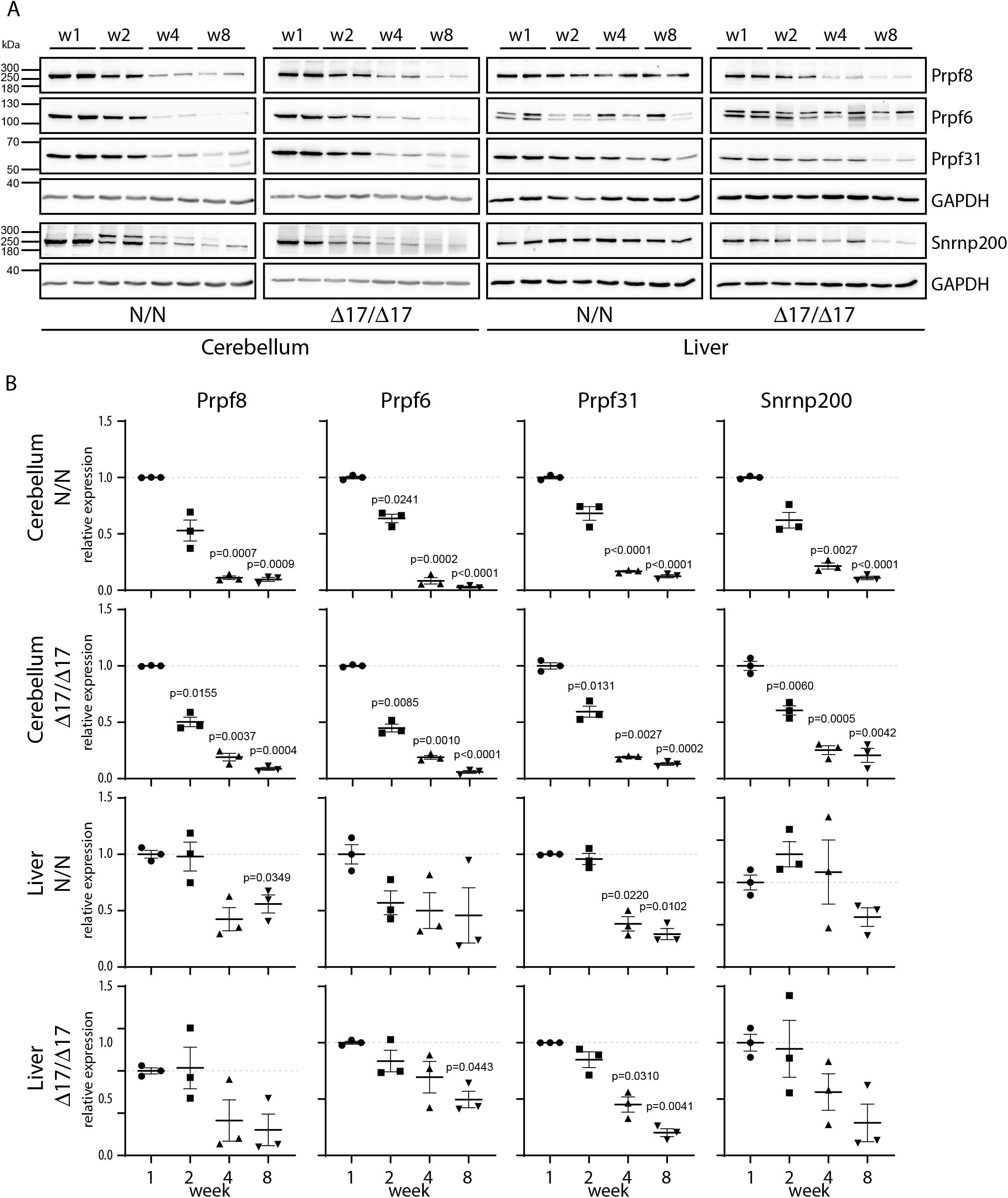
Expression of snRNP proteins is reduced in mutant animals. (*A*) Expression of snRNP-specific proteins was assayed in cerebella and liver of *Prpf8^Y2334N^* and *Prpf8^Δ17/Δ17^mice* at 1, 2, 4, and 8 weeks by Western blotting. Samples from two animals were analyzed per each timepoint. (*B*) Statistical analysis of protein expression at 1, 2, 4, and 8 weeks. Protein abundance in individual animals was first normalized to levels of GAPDH is the same tissue and then calculated as % of average value of the value at the first week animals. N=3. Statistical significance of differential protein abundance in the given genotype groups was examined by one-way ANOVA followed by Dunnett’s T13 test to find samples that significantly differ from expression in week 1 (mean with SEM are displayed together with p values).

In parallel to the measurement of Prpf8 protein levels we also investigated the expression *of Prpf8* RNA in 4 weeks old mice of both strains, and in 8- and 12-weeks old *Prpf8^Δ17^* and *Prpf8^Y2334N^* animals, respectively. The *Prpf8* transcripts were quantified in the RNA-Seq datasets and then verified in independent samples by RT-qPCR approach. The results surprisingly revealed similar or slightly enhanced abundance of *Prpf8* mRNA in *Prpf8^Y2334N/Y2334N^* and *Prpf8^Δ17/Δ17^* animals in all the examined timepoints when compared to levels of wt *Prpf8* (Fig. S14). These findings indicated that the amounts of Prpf8 in cerebellar cells are primarily regulated at the protein level.

To further analyze the relationship between RNA and protein expression of Prpf8, we took advantage of a mouse strain *Prpf8^Δ366^* that carries a large deletion of 366 bp that eliminated the 3’ part of *Prpf8* exon 37 until intron 38-39 (chr11:75,506,472-chr11:75,506,837; Fig. 1). Since no *Prpf8^Δ366/Δ366^* individuals were born to heterozygotic breeding pairs, the *Prpf8^Δ366^* allele is genetically a null variant, and this finding was in agreement with embryonic lethality previously described for full *Prpf8* deficiency (Graziotto et al., 2011). We did not detect any productive expression originating from the *Prpf8^Δ366^* allele, indicating a rapid decay of the faulty mRNA. In cerebella harvested from *Prpf8^Δ366/wt^* rodents the *Prpf8* mRNA content reached 60% of the control animals, which strongly argued against a substantial compensatory transcription from the *Prpf8* wt allele (Fig. 8A). This shortage in *Prpf8* transcription was, however, compensated at the level of Prpf8 protein synthesis and/or stability, since no differences were recorded in Prpf8 protein abundance estimated by Western blotting and immunohistochemistry (Fig. 8B-D). Consistently with the unchanged expression of the Prpf8 protein, the absence of one *Prpf8* copy did not provoke any pathological changes affecting the cerebellum (Fig. 8D), and the *Prpf8^Δ366/wt^* animals attained body weight equivalent to their wt littermates (Fig. S15A). These results demonstrated that loss of one *Prpf8* allele *per se* did not induce cerebellar degeneration and ruled out haploinsufficiency as a mode of action in the case of murine aberrant Prpf8. However, the physiological drop in Prpf8 abundance that coincides with cerebellar postnatal maturation, ongoing synaptogenesis and with the onset of circRNA formation suggests that reduced levels of Prpf8 and other splicing factors may sensitize granule neurons to Prpf8 mutations. Interestingly, analysis of the *Prpf8^Δ366^* strain breeding records revealed a potential distortion of paternal allele transmission (Fig. S15B). This observation may indicate that deficiency in Prpf8 and hence execution of splicing can impact the haploid stages of spermatogenesis, or alternatively the Prpf8 protein can execute unknown, extraspliceosomal functions as suggested for PRPF31 (Buskin et al., 2018).

**Figure 8.**
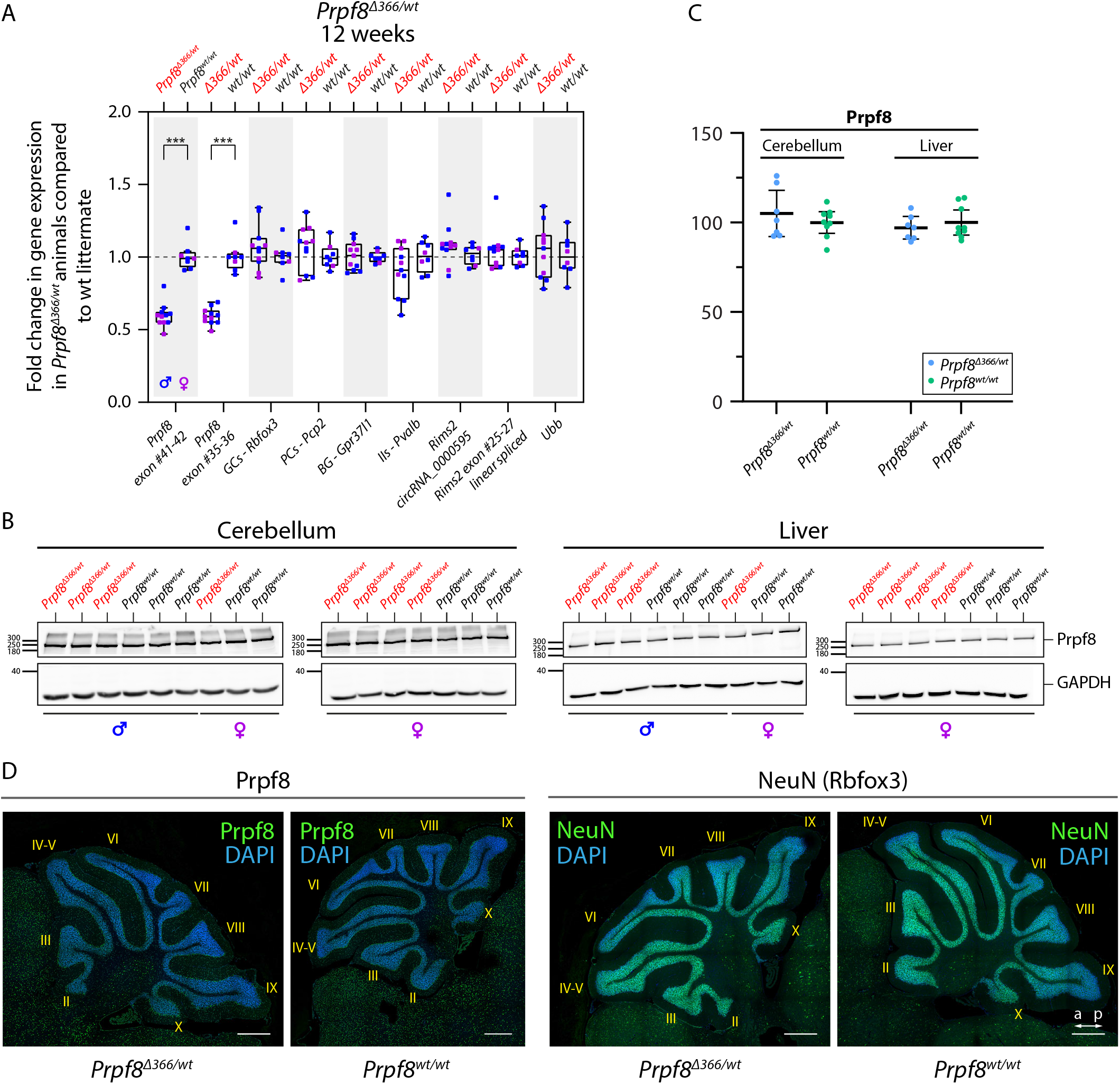
12-weeks-old *Prpf8^Δ366/wt^* mice do not exhibit any degenerative phenotype. (*A*) RT-qPCR-based quantification of selected transcripts in the cerebellum of 12-weeks-old *Prpf8^Δ366/wt^* mice and their *Prpf8^wt/wt^* littermates. GCs-granule neurons, PCs-Purkinje cells, BG-Bergmann glia, IIs-inhibitory interneurons. (*B*) Western blots from cerebellar and liver samples collected from 12 weeks old *Prpf8^Δ366/wt^* mice and their wt littermates. (*C*) Densitometric quantification of Western blots from two *Prpf8^Δ366^* cohorts (*Prpf8^Δ366/wt^* n=7, *Prpf8^wt/wt^* n=9) did not reveal any significant differences in Prpf8 protein levels in *Prpf8^Δ366/wt^* animals in neither cerebellar nor liver specimen. Protein abundance of Prpf8 in individual animals was first normalized to levels of GAPDH is the same tissue and then calculated as % of average value of wt animals. Statistical significance of differential protein abundance in the given genotype groups was examined by t-test using GraphPad software (displayed is mean with 95% confidentiality interval). (*D*) Immunohistochemical analysis of Prpf8 and NeuN abundance in cerebella of 12-weeks-old *Prpf8^Δ366/wt^* animals and their corresponding *Prpf8^wt/wt^* counterparts. Representative images are shown for each genetic condition. Nuclear counterstaining by DAPI, scale bar = 200 μm. a - anterior, p - posterior. For further characterization of *Prpf8^Δ366/wt^* animals see Fig. S15.

## Discussion

Spliceosomopathies are inborn disorders, where congenital mutations impair generic factors involved in spliceosome assembly and/or function, but the primary phenotypes manifest in distinct organs. Such selective organ impairment indicates that development and/or homeostasis of certain tissues is specifically sensitive to mutations that are elsewhere tolerated. In addition, the dissimilar outset dynamics imply that particular cell types are likely vulnerable only during a defined developmental or age window. This suggests that naturally occurring changes in the intracellular environment can create conditions for an outbreak of a pathology but mechanisms that triggers pathological changes are unknown. In this study, we established and analyzed two aberrant variants of splicing factor Prpf8 in a mouse experimental model and observed that both strains displayed rapid and specific decay of cerebellar granule neurons. To better understand the dynamic of splicing factors expression in the targeted tissue, we analyzed amount of selected key splicing proteins including Prpf8 during first eight weeks of life in three different tissues of wild type animals. We observed steep and significant drop of all tested splicing proteins in cerebellum and retina between week one and week four. Moreover, majority of tested splicing proteins were more downregulated in cerebellum then in retina. This physiological reduction of splicing components precedes the onset of neurodegeneration and we speculate that this could be one of the main reasons why mouse cerebellum is specifically sensitive to mutations in Prpf8.

This hypothesis is consistent with the fact that most RP-linked mutations in splicing components reduce their ability to form functional spliceosomal snRNPs, which further decreases the amount of splicing-competent complexes inside the nucleus (Gonzalez-Santos et al., 2008; Huranova et al., 2009; Linder et al., 2014; Malinova et al., 2017; Tanackovic et al., 2011b). The aberrant substitution p.Tyr2334Asn studied in this work does not affect the Prpf8 interaction with Prpf6 and Snrnp200 and snRNP maturations (Malinova et al., 2017) and (Fig. S1C). However, we showed that this mutation negatively impacts protein expression and further reduces already low levels of Prpf8 protein in cerebellum (Figs. 6 and 7). It should be noted that two pathological missense mutations in SNRNP200 did not either inhibit the assembly of snRNPs, but rather had a negative effect on the selection of correct splice sites (Cvackova et al., 2014). It is thus plausible that all RP mutations negatively affect more or less severely the function of the mutated protein. Most cells are able to tolerate this partial malfunction due to natural high expression of splicing proteins. However, cells and tissues with naturally low expression of these proteins are specifically vulnerable to mutation in splicing machinery. Surprisingly, our data show that the regulation of splicing protein occurs at the protein level because we observed, that the dosage compensation for the missing *Prpf8* allele occur at the protein but not mRNA level (Fig. 8). Therefore, monitoring mRNA levels is not the best indicator of actual splicing protein expression, at least in some tissues.

Next, we analyzed how RP mutations alters transcriptome in the targeted tissue in order to identify common defects between the two established RP-mimicking strains. We did not identify any significant overlap in terms of intron retention, exon skipping and alternative 5’ and 3’ splice site usage in 4 week old cerebella of *Prpf8^Δ17/Δ17^* and *Prpf8^Y2334N/Y2334N^* mice. However, it should be noted that only events with higher than 15% shift were considered and we cannot exclude that minor changes in splicing pattern of particular mRNAs are common to both Prpf8 RP variants. In contrast, we observed perturbation of specific subclasses of circRNAs in both strains that preceded the onset of neural death. Because circRNA are formed by the splicing machinery, deregulation of circRNA might be a direct effect of Prpf8 mutations. Consistently, inhibition of spliceosome activity was previously shown to disrupt the circRNA pool (Starke et al., 2015; Wang et al., 2019), and downregulation of Prpf8 in Drosophila increased expression of two model circRNAs (Liang et al., 2017).

Circular RNAs are highly abundant in neural tissues and cerebellar granule cells are substantially rich for circRNAs (Rybak-Wolf et al., 2015), yet their biological roles remain poorly characterized. Some circRNAs were proposed to regulate gene expression by interfering with miRNA and splicing pathways (Ashwal-Fluss et al., 2014; Hansen et al., 2013; Yu et al., 2017). Transcription of numerous circRNAs, including *Rims2* circ_0000595, was found to be uncoupled from the parental mRNAs ((Rybak-Wolf et al., 2015) and this work), which suggests that the circRNA production is regulated independently of the spatial and temporal dynamics of the host gene expression. Indeed, the formation of the *Rims2* circ_0000595 was *in vitro* induced by neuronal maturation, and in the developing mouse brain the circ_0000595 amounts substantially increased around P10 to peak in the adulthood (Rybak-Wolf et al., 2015). This time window coincides with establishment of connections between mossy fibers and granule dendrites, and of parallel fibers with Purkinje cells (Kano and Watanabe, 2019). In agreement, many further circRNAs were shown to change their abundance in response to the onset of synaptogenesis (You et al., 2015), suggesting the circRNA species likely contribute to establishment and/or maintenance of synapses. In accord, circRNAs and specifically the *Rims2* circ_0000595 were found enriched in the brain synaptosome (Rybak-Wolf et al., 2015). Genetic ablation of the cerebellar most prominent circRNA *Cdr1os* mmu_circ_0001878 provoked excitatory synapse malfunction (Kleaveland et al., 2018; Piwecka et al., 2017), and specific elimination of the *Rims2* circRNA by a shRNA approach evoked retinal neuron apoptosis (Sun et al., 2021). Together, circRNAs likely represent components of a complex network neurons utilize to fine-tune their transcriptome and to modulate synaptic events, and their disturbance might compromise neuronal fitness.

What can the newly established mouse models tell us about retinal degeneration in humans? In mice, the Rims proteins are localized to presynaptic active zones in conventional synapses, but also to presynaptic ribbons in photoreceptor ribbon synapses (Wang et al., 1997). Consistently, inborn errors in human *RIMS1* and *RIMS2* underlie cone-rod dystrophies (OMIM #603649 and #618970) (Johnson et al., 2003; Mechaussier et al., 2020). However, it is not known whether the *RIMS* mutations may simultaneously impact the biogenesis of the cognate circRNAs. Retina belongs to neuronal tissues expressing high amount of various circRNA species, many of them are conserved among human and mouse including *RIMS1* hsa_circ_0132250, RIMS2 hsa_circ0005114 (corresponding to the murine *Rims 1* and *Rims2* circRNAs, respectively (Fig. S15C,D)) and circHIPK3 (Izuogu et al., 2018; Mellough et al., 2019; Rahimi et al., 2021; Sun et al., 2019). Downregulation of the circHIPK3 accelerated apoptosis in cultured lens epithelial cell (Liu et al., 2018), which indicates that circRNAs indeed play an important role in eye homeostasis. Overall, the novel mouse models of aberrant Prpf8 suggest that deregulation of circRNAs could represent a pathological mechanism that co-provokes the retinal dystrophy in splicing factor-RP.

In summary, we suggest that vulnerability of the murine cerebellum towards mutant Prpf8 proteins probably originates from a physiological drop in splicing factor abundance that coincides with ongoing processes of tissue maturation and circRNA-mediated synaptogenesis. Deregulation of circular RNAs may represent a pathological mechanism that perturbs cellular homeostasis and initiates apoptosis of vulnerable cells. Finally, the infiltration and activation of microglia and astrocytes can further intensify and complete the damage of the neuron tissues.

## Material and Methods

### Generation, breeding, and genotyping of mutant mice

All animal models and experiments of this study were ethically reviewed and approved by the Animal Care Committee of the Institute of Molecular Genetics (Ref. No. 45/2020). All novel alleles of *Prpf8* were established in the Transgenic and Archiving Module of the Czech Centre for Phenogenomics (BIOCEV, Czech Republic). The pathogenic substitution Prpf8 Tyr2334Asn was introduced to the exon 42 of the *Prpf8* gene using Cas9-mediated homology-directed repair (HDR); the short guide RNA (sgRNA) specifically targeting the vicinity of Tyr2334 (*Prpf8* ex42 sgRNA-#1) was devised with assistance of the MIT Design website (https://crispr.mit.edu) (Ran et al., 2013). The *Prpf8* ex42 sgRNA-#1 was assembled and it’s efficacy first assayed *in vitro*. In detail, complementary guide oligodeoxynucleotides were first phosphorylated by T4 Polynucleotide Kinase (Thermo Fisher) in ATP-containing T4 DNA Ligase buffer (Thermo Fisher), then denatured for 5 min at 95°C and annealed by gradual cooling down at −0.1 °C/sec using a PCR thermocycler (BioRad T100). Annealed oligos were subsequently inserted into BbsI-digested vector pX330-U6-Chimeric_BB-CBh-hSpCas9 (Addgene #42230), where co-expression of sgRNA and human codon-optimized *S.pyogenes* Cas9 is driven from human U6 promoter and CAGEN promoter/enhancer, respectively (Cong et al., 2013). Efficacy of the *Prpf8* ex42-#1 sgRNA towards the target sequence in *Prpf8* (5’-AAGGAAGTAACTAGGCATAG-AGG; 11:75,509,277-75,509,299 coding strand) and two predicted, highest-scoring off-target loci (chr.11 5’-AAGGAACTAACTAGGCATGG-AGG 11:5,006,551-5,006,573; chr.14 5’-AAGAAAGAAACTTGGCATAG-CAG 14:56,271,668-56,271,690) was assessed in transfected HeLa cells using TurboRFP open reading frame reconstitution reporter plasmid (pAR-TurboRFP; Addgene #60021) as described previously (Kasparek et al., 2014).

To prepare individual components for the microinjection mixture, Cas9 was transcribed from linearized parental pX330 plasmid using mMESSAGE mMACHINE T7 kit (Ambion/Thermo Fisher), the resulting mRNA was polyadenylated using Poly(A) Tailing kit (Ambion/Thermo Fisher), and purified with RNA Clean & Concentrator kit (Zymo Research). PCR template for *in vitro* sgRNA synthesis included T7 promoter, 20nt sgRNA and tracrRNA sequences was amplified from pX330-*Prpf8* ex42-#1 sgRNA plasmid using T7 promoter-containing and tracrRNA primers. *Prpf8* ex42-#1 sgRNA was subsequently produced by MEGAshortscript T7 Transcription Kit (Ambion/Thermo Fisher), and purified using the ssDNA/RNA Clean & Concentrator kit (Zymo Research). A synthetic oligodeoxynucleotide or double-stranded DNA fragment encoding the mutated Tyr2334Asn motif were used in parallel as potential HDR donors to navigate the homology-directed repair machinery. Briefly, the donor oligodeoxynucleotide contained 35bp homology arms flanking a central region where the sequence encoding for five Prpf8 C-terminal residues spanning Glu2331 to STOP codon was silently mutagenized to prevent re-editing of once modified allele (Krchnakova et al., 2019). These substitutions were deliberately designed to introduce novel DraI and MluI restriction sites that allowed downstream genotypic screening by Restriction Fragment Length Polymorphism (RFLP). In addition, the donor oligodeoxynucleotide contained three terminal linkages at both 5’- and 3’-modified to phosphorothioate. The HDR oligo was purchased from Sigma Aldrich (vendor purification by HPLC), and was additionally purified using ssDNA/RNA Clean & Concentrator kit (Zymo Research). In contrast, the double stranded DNA HDR donor was prepared in a step-wise manner in pGEM-T Easy vector (Promega) by first cloning in approximately 800bp long homology arms that were amplified from C57BL6/N-derived genomic DNA (Left arm: 799bps upstream of Tyr2334; Right arm: 864bp downstream of Tyr2334; total insert length 1,666bps chr11:75,508,482-75,510,147). We subsequently used mutagenic PCR to alter the vicinity of Tyr2334 with silent substitutions that were identical to those present in the HDR oligo. To prepare the final dsDNA fragment destined for the microinjection, the donor portion was amplified from the pGEM-T Easy Prpf8 Y2334N vector using Phusion High-Fidelity DNA polymerase (Thermo Fisher), the sample was then treated with DpnI to remove the parental plasmid, and purified using DNA Clean & Concentrator kit (Zymo Research). Prior to microinjection, the exact sequences of both the oligo and dsDNA fragment were verified by Sanger sequencing. The microinjection sample comprised Cas9 mRNA (100ng/μL final concentration), sgRNA (50 ng/μL), and either dsDNA fragment (5 ng/μL) or 10μM oligodeoxynucleotide. The final mixture was filter-sterilized by passing through 0.22μm Millex-GV PVDF filter (Millipore), microinjected into pronuclei of C57BL6/N-derived zygotes and those were transferred into pseudo-pregnant recipient mice.

The *Prpf8 Δ366* strain was obtained secondary to genome editing effort aimed at residue Ser2118 in exon38 of the *Prpf8* gene. TALENs were designed using TAL Effector Nucleotide Targeter 2.0 (tale-nt.cac.cornell.edu (Doyle et al., 2012)), assembled using the Golden Gate Cloning protocol, and inserted into the ELD-KKR backbone plasmids as described previously (Kasparek et al., 2014). The DNA binding domains of TALEN comprised following repeats: NN-HD-NG-NG-NI-NI-NN-NI-NI-NN-NG-NG-HD-NI-NG-HD-NG (5’-TALEN-Prpf8 ex38), and NI-NN-HD-NG-NI-HD-NG-HD-NI-HD-NG-NG-NN-NN-NN-HD-NI-HD (3’-TALEN-Prpf8 ex38). Linearized plasmids were *in vitro* transcribed using mMESSAGE mMACHINE T7 kit (Ambion/Thermo Fisher), the resulting mRNA polyadenylated using Poly(A) Tailing kit (Ambion/Thermo Fisher), and purified with RNA Clean & Concentrator kit (Zymo Research). TALEN-encoding mRNA (20ng/uL each) was mixed with targeting single-stranded oligodeoxynucleotide (10μM final concentration; Sigma-Aldrich), and the final solution was filtered through 0.22μm Millex-GV PVDF filter prior to microinjection into C57BL6/N-derived zygotes.

In the IMG mouse husbandry animals were maintained under 12/12-h light cycle, and access to food and water was provided *ad libitum*. All novel *Prpf8* strains were generated in zygotes isolated from C57BL6/N substrain and successively back-crossed to C57BL6/J background to eliminate the *Crb1^rd8^* mutation that may else confound interpretation of ocular phenotypes (Mattapallil et al., 2012). Phenotypic analysis were carried out after seven generations of breeding to C57BL6/J, when contribution of the recurrent parent genome is >99% (Visscher, 1999). Routine PCR genotyping was performed with tail biopsies snipped from three-week-old animals on weaning. Tissue sample was incubated overnight at 56°C in 100μL of lysis solution (10mM TrisCl pH8.3, 50mM KCl, 0.45% (v/v) Nonidet P 40 Substitute, 0.45% (v/v) TWEEN 20, 0.1 mg/mL gelatin from porcine skin; Sigma Aldrich) supplemented with Proteinase K (0.2 mg/mL; Thermo Fisher), then heat inactivated for 15 minutes at 70°C, and 1 μL of the crude lysate was directly utilized as input for PCR reactions (DreamTaq Green PCR Master Mix (Thermo Fisher) supplemented with 1M betaine and 3% DMSO (Sigma Aldrich)). To increase the specificity of the genotyping primers, an extra mismatch was deliberately introduced at the third position from the 3’ end of one of the primers; primers were designed using PRIMER1 ARMS-PCR tool (primer1.soton.ac.uk/primer1.html). In contrast, RFLP was performed with amplicons produced by DreamTaq DNA polymerase (Thermo Fisher), which were purified by sodium acetate-ethanol precipitation, and then subjected to restriction digestion with MluI enzyme (New England Biolabs) to identify mutant animals. To reveal the spectrum of Prpf8 exon 42 genetic variants present in the founder animals, RFLP amplicons were cloned into pGEM-T Easy vector and subsequently analyzed by Sanger sequencing. List of all primers relevant to mice generation and breeding can be found in Table S1.

### Immunohistochemistry

Mice were sacrificed at indicated timepoints by CO2 asphyxiation and *post mortem* perfused with 10mL phosphate-buffered saline (PBS) followed by 10mL of 4% paraformaldehyde (PFA; Electron Microscopy Services)/PBS by intracardial puncture. Following decapitation, skin removal, and skull opening, whole head was fixed *en bloc* in 4% PFA/PBS for 24 hrs, transferred to 70% ethanol (Penta), and then cerebella were dissected out for subsequent tissue processing (Leica ASP200s) and paraffin embedding (Leica EG1150H). 5-μm sections were stained according to standard protocols. Briefly, specimens were deparaffinized in xylene (Penta) and rehydrated through graded ethanol series; antigen retrieval was performed by boiling in 10mM sodium citrate buffer pH 6.0 (Sigma Aldrich) in pressure cooker for 20 minutes. Endogenous peroxidase activity was blocked by immersion in 3% H2O2 (Sigma Aldrich) in methanol (Penta) for 10 minutes; interference from endogenous biotin and/or streptavidin-binding activity was moreover reduced by Avidin/Biotin Blocking kit (Thermo Fisher). Tissue sections were incubated overnight in a humidified chamber with following primary antibodies: anti-CD45 (ab10558, Abcam), anti-GFAP (#12389, Cell Signaling), anti-IP3R-I/II/III (sc-377518, Santa Cruz), anti-NeuN (#24307, Cell Signaling), anti-Prpf8 (ab79237, Abcam), anti-PSD95 (#3409, Cell Signaling), anti-S100 beta (ab52642, Abcam). Subsequently, biotin-conjugated secondary antibodies (Biotin-XX Goat anti-Mouse IgG B2763, or Biotin-XX Goat anti-Rabbit IgG B2770, Thermo Fisher) were utilized at a dilution of 1:750, and the signal was finally visualized with Alexa Fluor 488-labelled streptavidine (S11223, Thermo Fisher). Nuclei were counterstained with DAPI (Sigma). Immunofluorescence images were acquired using Dragonfly Spinning Disc confocal microscope (Andor), deconvolved using Huygens Professional software, and processed using FiJi. Control sections accompanying immunohistochemistry slides were counterstained with Hematoxylin solution modified according to Gill II (Penta), followed by 0.5% aqueous solution of Eosin-Y (Sigma Aldrich). NissI staining was performed with 0.1% acidic Cresyl Violet solution (Abcam) according to the manufacturer’s instructions.

Histopathological analyses were carried out on biopsies harvested from mice euthanized by cervical dislocation. For investigation of eye specimen, skinned heads were first fixed for 48 hours in Davidson’s fixative before eye removal; other examined tissues were routinely fixed for 24 hours in phosphate buffered 10% formalin and then transferred to 70% ethanol solution. Fixed samples were processed using automated tissue processor (Leica ASP6025), and embedded in paraffin blocks using a Leica EG1150H embedding station. 2-μm sections were counterstained with Hematoxylin, Harris modified, in combination with Eosin B (Sigma) processed in Leica Stainer Integrated Workstation (ST5020-CV5030), in combination with the Leica CV5030 coverslipper. Special stains, including Alcian Blue, Congo Red, Luxol Fast Blue, and Masson’s Trichrome stains, were performed using Ventana BenchMark Special Stains platform (Roche). Sections were scanned using AxioScan.Z1 microscope (Zeiss) using 20x Plan-Apochromat objective.

Apoptotic cells were visualized with NeuroTACS II In Situ Apoptosis Detection Kit (Trevigen), and the signal was subsequently enhanced using a combination of Vectastain ABC Kit (Vector Laboratories) and Alexa Fluor 488 Tyramide Super Boost Streptavidin kit (Thermo Fisher).

Composition of animal cohorts collected for staining procedures (all genetic conditions contained animals of both genders):

1. Prpf8 Y2334N strain at 4 weeks - Histopathology: *Prpf8^Y2334N/Y2334N^* n=1, *Prpf8^Y2334N/wt^* n=4, *Prpf8^wt/wt^* n=1 (Fig. S3A)
2. Prpf8 d17 strain at 4 weeks - Histopathology: *Prpf8^Δ17/Δ17^* n=3, *Prpf8^Δ17/wt^* n=4, *Prpf8^wt/wt^* n=1 (Fig. S4A)
3. Prpf8 Y2334N strain at 6 weeks - Histopathology: *Prpf8^Y2334N/Y2334N^* n=1, *Prpf8^Y2334N/wt^* n=6, *Prpf8^wt/wt^* n=1 (Fig. S3B)
4. Prpf8 Y2334N strain at 12 weeks - Histopathology: *Prpf8^Y2334N/Y2334N^* n=2, *Prpf8^Y2334N/wt^* n=1, *Prpf8^wt/wt^* n=1 (Fig. S3C)
5. Prpf8 d17 strain at 6 weeks - Histopathology: *Prpf8^Δ17/Δ17^* n=3, *Prpf8^Δ17/wt^* n=8, *Prpf8^wt/wt^* n=2 (Fig. S4B)
6. Prpf8 Y2334N strain at 17 weeks - Histopathology: *Prpf8^Y2334N/Y2334N^* n=4, *Prpf8^Y2334N/wt^* n=6, *Prpf8^wt/wt^* n=4 (Fig. S3D)
7. Prpf8 d17 strain at 15 weeks - Histopathology: *Prpf8^Δ17/Δ17^* n=4, *Prpf8^Δ17/wt^* n=5, *Prpf8^wt/wt^* n=4 (Fig. S4C)
8. Prpf8 Y2334N strain at 22 weeks - Histopathology: *Prpf8^Y2334N/Y2334N^* n=5, *Prpf8^Y2334N/wt^* n=9, *Prpf8^wt/wt^* n=5 (Figs. S3E and S5A, C, E, G)
9. Prpf8 d17 strain at 22 weeks - Histopathology: *Prpf8^Δ17/Δ17^* n=3, *Prpf8^Δ17/wt^* n=8, *Prpf8^wt/wt^* n=5 (Figs. S4D and S5B, D, F, H)
10. Prpf8 d17 strain at 22 weeks - Histopathology, OCT, ERG: *Prpf8^Δ17/Δ17^* n=3, *Prpf8^Δ17/wt^* n=5, *Prpf8^wt/wt^* n=7 (Fig. S6A-F)
11. Prpf8 Y2334N strain at 12 weeks - NissI, IHC-P: *Prpf8^Y2334N/Y2334N^* n=6, *Prpf8^Y2334N/wt^* n=7, *Prpf8^wt/wt^* n=5 (Figs. 2B,D and S8)
12. Prpf8 d17 strain at 8 weeks - NissI, IHC-P: *Prpf8^Δ17/Δ17^* n=4, *Prpf8^Δ17/wt^* n=10, *Prpf8^wt/wt^* n=2 (Figs. 2A,C and S8)
13. Prpf8 Y2334N strain at 4 weeks - IHC-P: *Prpf8^Y2334N/Y2334N^* n=3, *Prpf8^Y2334N/wt^* n=4, *Prpf8^wt/wt^* n=1 (Fig. S10A)
14. Prpf8 d17 strain at 4 weeks - IHC-P: *Prpf8^Δ17/Δ17^* n=3, *Prpf8^Δ17/wt^* n=4, *Prpf8^wt/wt^* n=1 (Fig. S10A)
15. Prpf8 d17 strain at 5 weeks - IHC-P, TUNEL: *Prpf8^Δ17/Δ17^* n=4, *Prpf8^Δ17/wt^* n=6, *Prpf8^wt/wt^* n=4 (Fig. S10B)
16. Prpf8 Y2334N strain at 5 weeks - IHC-P: *Prpf8^Y2334N/Y2334N^* n=2, *Prpf8^Y2334N/wt^* n=3, *Prpf8^wt/wt^* n=3 (Fig. S10C)

### Grip strength measurement

Testing was performed in a room with light intensity set to 110 lux. To assess neuromuscular function, each mouse was gently pulled by tail over metal bars. The grip of only forelimbs and combined forelimbs and hindlimbs was thereby recorded using an automated Grip Strength Meter (Bioseb). The average of three trials was calculated; data are presented as absolute grip strength (g) and normalized to body weight.

### Open field

The open field test evaluates animal motility triggered by exploratory drive in a new environment. Fully automated analysis of animal behavior in open space was based on video tracking system (Viewer, Biobserve GmbH). The software distinguishes an animal as an object contrasting with the background. Testing apparatus was uniformly illuminated with the light intensity of 200 lux in the center of the field. Each animal was tested in open field for 20 minutes. The distance travelled, average speed, and resting time were automatically computed for each 5 min-long interval.

### Electroretinography (ERG)

Animals were dark-adapted over-night in red, individually ventilated cages (Tecniplast). Experiments were carried out in general anaesthesia (Tiletamine + Zolazepam, 30 + 30 mg/kg, i.m., Virbac). Anaesthetized mice were kept on a heating pad and their eyes were protected against drying using a small amount of transparent eye gel (Vidisic, Bausch&Lomb). Experiments were approved by the Animal Care and Use Committee of the Academy of Sciences of the Czech Republic.

Single flash stimulation and full field ERG recording was performed inside a ganzfeld globe controlled by RETIanimal system (Roland Consult). Active golden ring electrodes were gently positioned on the cornea and reference and grounding needle electrodes were placed subdermally in the middle line of the snout and in the back of the animal, respectively. Signal was band-pass filtered between 1 - 300 Hz and recorded with 1024 sampl/s resolution. Stimulation was repeated several times and 20 - 30 individual responses were averaged.

Signals were analyzed using custom scripts in Matlab (MathWorks). Amplitudes and latencies (implicit times) of waves a and b were quantified. The oscillatory potential (OP) was extracted from the original recordings by high-pass filtering with cut-off frequency set to 70 Hz. Amplitude and latency of the four major OP peaks was quantified and then summed. Statistical analysis of data where two parameters were changing, i.e. genotype of animals and intensity of the stimulation, 2-way ANOVA was used and p-values for the factor of genotype were reported.

### Optical coherence tomography (OCT)

The optical coherent tomograph (OCT, Heidelberg Engineering) scans and quantifies the reflection of a light beam sent from the layers of the retina and composes cross-sectional images of the retina (57 cross-sectional images as minimum). Each cross-section image of the retina was evaluated and following parameters were measured: the thickness and the gross morphology of the retina, form and position of the optical disc, and the superficial blood vessels and their pattern. The animals were anaesthetised using intramuscular injection of Tiletamine + Zolazepam (30 + 30 mg/kg, Virbac), and the pupils were dilated using eye drops containing 0.5% (w/v) Atropin (Samohyl).

### Protein extracts and Immunoblotting

Dissected cerebella and control liver snips were rinsed in ice-cold PBS, briefly chopped with a razor blade, and then homogenized with a 5mm generator probe in 0.5mL of modified RIPA buffer (composition: 50mM TrisCl pH 7.4, 150mM NaCl, 1% (v/v) Triton X-100, 0.5% (w/v) Sodium Deoxycholate, 0.1% (w/v) SDS, 1mM EDTA, 10mM NaF, 1mM PMSF, and 1:200 Protease Inhibitor Cocktail Set III (all chemicals purchased from Sigma Aldrich)). Cell disruption was moreover promoted by sonicating the crude lysate with thirty ultrasonic pulses of 1 sec that were interrupted by 1 sec pauses (sonicator IKA Labortechnik). Protein concentration was determined with Pierce BCA Assay Kit (Thermo Fisher); loads comprised 20μg of total protein for liver samples and 30μg for brain tissue, respectively. Immunoblotting was performed according to established protocols (Malinova et al., 2017), using the following primary antibodies: anti-GAPDH (#5174, Cell Signaling), anti-NeuN (#24307, Cell Signaling), anti-Prpf8 (ab79237, Abcam), anti-Prpf6 (#sc-166889, Santa Cruz Biotechnology), anti-Prpf31 (#188577, Abcam), and anti-Snrnp200 (#HPA029321, Sigma-Aldrich). Peroxidase-linked anti-rabbit secondary antibody (#7074, Cell Signaling) was utilized in combination with SuperSignal West Femto Maximum Sensitivity Substrate or West Pico PLUS Chemiluminescent Substrate (Thermo Fisher) to obtain a chemiluminescent signal.

### Immunoprecipitation

Dissected cerebella were rinsed with ice-cold PBS, chopped with a razor blade, and transferred into 1 ml of NET2 buffer (150 mM NaCl, 0.05% NP-40, 50 mM Tris-HCl pH 7.4, supplemented with 1:200 Protease Inhibitor Cocktail Set III (#539134, Merck Millipore)), where they were homogenized for 5 secs on ice. The crude lysates were then sonicated with 30 consecutive pulses of 1 sec (60 % amplitude), and cleared by centrifugation at 15000xg for 5 min at 4 °C. The cleared lysates were further incubated with 4 μg of Prpf8 antibody (#79237, Abcam) or with 4 μg of control IgG antibody (#I5381, Sigma-Aldrich) for 1 hour at 4 °C with continuous rotation. 30 μl of Protein G agarose beads (#sc-2002, Santa Cruz Biotechnology) were then added to the mixture, and the reaction was incubated for additional 2 hours at 4 °C with continuous rotation. The beads were then washed five times with NET2 buffer, and resuspended in 2x sample buffer (250 mM Tris-HCl pH 6.8, 20% glycerol, 4% SDS, 0.02% bromphenol blue), supplemented with 24 mM DTT (Sigma-Aldrich)). Precipitated proteins were analyzed by Western blotting. Primary antibodies used for the immunoblotting were: anti-GAPDH (#5174T, Cell Signaling Biotechnology), anti-Prpf6 (#sc-166889, Santa Cruz Biotechnology), anti-Prpf8 (#79237, Abcam), anti-Prpf31 (#188577, Abcam), and anti-Snrnp200 (#HPA029321, Sigma-Aldrich).

### RT-qPCR

Dissected cerebella (bulk tissue) were briefly rinsed in ice-cold PBS and homogenized in 750 μL of TRIzol reagent (Thermo Fisher). Total RNA was extracted using Direct-Zol RNA Miniprep Kit (Zymo Research) including in-column DNase-I treatment. First-strand cDNA synthesis was performed with Superscript III Reverse transcriptase (Thermo Fisher) supplemented with random hexamers (Sigma Aldrich) and RiboLock RNase inhibitor (Thermo Fisher). RT-qPCR was carried out in technical triplicates using SYBR Green I Master Mix and Lightcycler 480 apparatus (Roche); negative control was represented by cDNA synthesis samples performed in the absence of the reverse transcriptase enzyme. Primers are listed in Table S1.

Statistical evaluation of the RT-qPCR data: for individual genes, average of threshold cycle (Ct) values from technical triplicates was first normalized to glyceraldehyde-3-phosphate dehydrogenase (*Gapdh*) expression to obtain the ΔCt values. Relative mRNA abundance was then calculated as fold change (FC) difference of homozygous (hom) or heterozygous (het) animals to *Prpf8^wt/wt^* controls using the 2^^ΔΔCt^ approach (i.e. 2^^ΔCt(wt/wt)-ΔCt(hom)^ or 2^^ΔCt(wt/wt)-ΔCt(het)^, respectively). The fold change values are plotted in Box-and-whiskers diagram according to Tukey with indicated position of median; statistical significance of differential expression in the given genotype groups was examined by t-test using GraphPad software. * P < 0.05, ** P < 0.01, *** P < 0.001.

RT-qPCR linear model: to model qPCR data, we used linear mixed-effects model as outlined in (Matz et al., 2013; Steibel et al., 2009). Briefly, this method can be understood as an extension of the delta-delta method. The main advantage of mixed-effects models is that they can account for the correlations in the data introduced at multiple levels (replicates from the same animal, samples in the same qPCR runs, animals from the same litter). In designs involving such correlations, using a mixed-effects model provides better quantification of uncertainty than simpler approaches such as averaging replicates prior to making comparisons. The models were fitted using the brms R package (http://dx.doi.org/10.18637/jss.v080.i01). For the experiment with most data points (*Prpf8^Δ17^* at 4 weeks), the model included fixed effects of primer, and the full interaction of primer and genotype, and primer and sex. Varying intercepts were then introduced to model variability between qPCR runs, litters, and animals. We also let the residual standard deviation to vary between primers and qPCR runs. For other experiments this full model failed to converge due to lack of data to inform all the coefficients and some varying intercepts were omitted. We checked that the reduced models did not underestimate the biological and technical variability using posterior predictive checks https://doi.org/10.1111/rssa.12378.

To compare the expression of circular/alternative exons to the canonical transcripts, we used the fitted model to compute posterior predictions for the ratio of the circular/alternative exon species to the canonical species within each genotype. We then computed estimates of the ratio of those ratios between the genotypes. In other words, we estimated to what extent does the proportion of the products converted to circular/alternative exon change between the genotypes, which we find more relevant to the current investigation than comparing the absolute expression of the circular/alternative forms. We note that the estimates of this “ratio of ratios” do not depend on the choice of reference, only on the relative PCR efficiency of the two species. Here, we report results assuming equal efficiency, but almost all of the qualitative patterns are robust to even noticeably different efficiencies (data not shown). To compare the absolute expression of the linear forms, we used *Gapdh* as reference. Complete code and data to reproduce the qPCR analysis and the detailed outputs of the models are available at https://github.com/cas-bioinf/prpf8.

### RNA-Seq

Freshly dissected cerebella (bulk tissue) were briefly rinsed in ice-cold PBS and immediately homogenized in 750 μL of TRIzol reagent (Thermo Fisher). The homogenizer probe was sanitized between individual specimen to prevent any potential carry-over. Total RNA was extracted using Direct-Zol RNA Miniprep Kit (Zymo reseach) including in-column DNase-I treatment, and it’s quality was assessed using the 2100 Bioanalyzer (Agilent); only samples with RNA Integrity Number (RIN)>8 were permitted for downstream processing. Removal of ribosomal RNAs from samples collected from 4- and 8-weeks-old *Prpf8^Δ17^* mice, and 12-weeks old *Prpf8^Y2334N^* animals was achieved using RiboCop rRNA Select and Deplete rRNA Depletion Kit v1.2 (Lexogen), and libraries were generated using Total RNA-Seq Library Prep Kit (Lexogen). Cerebellar specimen gained from 4-weeks old *Prpf8^Y2334N^* mice were processed using KAPA RNA Hyperprep Kit with RiboErase (Kapa Biosystems/Roche) due to discontinuation of the Lexogen library construction kit. Libraries were sequenced with single-end 75nt reads (8-weeks-old *Prpf8^Δ17^* mice, 12-weeks old *Prpf8^Y2334N^* animals), or with pair-end 75+75nt reads (4-weeks-old *Prpf8^Δ17^* mice, 4-weeks old *Prpf8^Y2334N^* animals) on Illumina NextSeq 500 System. A total of 10-14 animals of both genders was selected for individual RNA-Seq runs, representation of individual genotypes was thereby as follows: 1. 12-weeks old *Prpf8^Y2334N^* animals (a total of 14 animals): n=6 for *Prpf8^Y2334N/Y2334N^* biological replicates (3xmale, 3xfemale), n=4 for *Prpf8^Y2334N/wt^* mice (2xmale, 2xfemale), and n=4 for *Prpf8^wt/wt^* litter matched controls (2xmale, 2xfemale); 2. 8-weeks-old *Prpf8^Δ17^* mice (a total of 10 animals): n=6 for *Prpf8^Δ17/Δ17^* biological replicates (4xmale, 2xfemale), and n=4 for *Prpf8^wt/wt^* litter matched controls (2xmale, 2xfemale); 3. 4-weeks-old *Prpf8^Δ17^* animals (a total of 10 animals): n=6 for *Prpf8^Δ17/Δ17^* biological replicates (4xmale, 2xfemale), and n=4 for *Prpf8^wt/wt^* litter matched controls (2xmale, 2xfemale); 4. 4-weeks old *Prpf8^Y2334N^* animals (a total of 10 animals): n=6 for *Prpf8^Y2334N/Y2334N^* biological replicates (4xmale, 2xfemale), and n=4 for *Prpf8^wt/wt^* litter matched controls (2xmale, 2xfemale).

### Differential gene expression and alternative splicing analysis

For gene level expression quantification, a bioinformatic pipeline nf-core/rnaseq (doi.org/10.5281/zenodo.3503887) was used (version 1.3 for 8-weeks-old *Prpf8^Δ17^* mice and 12-weeks old *Prpf8^Y2334N^* animals; version 1.4.2 for 4-weeks-old *Prpf8^Δ17^* mice and 4-weeks old *Prpf8^Y2334N^* animals). Individual steps included removing sequencing adaptors with Trim Galore! (www.bioinformatics.babraham.ac.uk/projects/trim_galore), mapping to reference genome GRCm38 (Ensembl annotation version 94) with HISAT2 (Kim et al., 2015), and quantifying expression on gene level with featureCounts (Liao et al., 2014). Per gene uniquely mapped read counts served as input for differential expression analysis using DESeq2 R Bioconductor package (Love et al., 2014), done separately for each sequencing run data. Prior to the analysis, genes not detected in at least two samples were discarded. We supplied experimental model assuming sample genotype as main effect, while accounting for breeding litter (for 8-weeks-old *Prpf8^Δ17^* mice samples) or gender (for 4-weeks-old *Prpf8^Δ17^* mice and 4-weeks old *Prpf8^Y2334N^* animals) as batch effect. Resulting per gene expression log_2_-fold changes shrunken using the adaptive shrinkage estimator (Stephens, 2017) were used for differential expression analysis. Genes exhibiting |Log_2_ Fold change |>1 and statistical significance FDR < 0.05 between compared groups of samples were considered as differentially expressed. As next, gene set over representation analysis with differentially expressed genes was done using gene length bias aware algorithm implemented in goseq R Bioconductor package (Young et al., 2010) against KEGG pathways and GO terms gene sets. Alternative splicing analysis was assessed using ASpli R Bioconductor package (Mancini et al., 2021) version 2.0.

### CircRNAs quantification and differential expression

CircRNAs back-spliced junctions sites were determined independently in each individual sequencing run data using CIRI2 tool (Gao et al., 2018)following respective guidelines with the same genome reference data supplied as for the gene level analysis. Expression of circRNAs was consequently quantified using CIRI2 (adaptor-trimmed data for 8-weeks-old *Prpf8^Δ17^* mice and 12-weeks old *Prpf8^Y2334N^* animals), or CIRIquant tool (Zhang et al., 2020) (full length data for 4-weeks-old *Prpf8^Δ17^* mice and 4-weeks old *Prpf8^Y2334N^* animals). Analysis of alternative splicing and quantification of exon expression within detected circRNAs was done for 4-weeks-old *Prpf8^Δ17^* mice and 4-weeks old *Prpf8^Y2334N^* animals with CIRI-AS tool provided alongside CIRI2. Differential expression analysis of circRNAs was done using either DESeq R Bioconductor package (for 8-weeks-old *Prpf8^Δ17^* mice and 12-weeks old *Prpf8^Y2334N^* animals) or EdgeR (McCarthy et al., 2012) R Bioconductor package pipeline provided alongside CIRIquant (for 4-weeks-old *Prpf8^Δ17^* mice and 4-weeks old *Prpf8^Y2334N^* animals). For all sequencing data, the same parameters for defining differentially expressed circRNAs were set (|Log_2_ Fold change|>1 and FDR<0.05). For parental genes of differentially expressed circRNAs, GO terms gene sets over representation was analyzed (for 8-weeks-old *Prpf8^Δ17^* mice and 12-weeks old *Prpf8^Y2334N^* animals only) using goseq R Bioconductor package without gene length bias considered.

### Generation and analysis of circRNA reporters

The minimal sequences surrounding the circularized exons of the differentially expressed circRNAs were identified from the RNA-seq data and were sorted keeping the maximum length of the construct within 5kbp simultaneously taking into account the repeat elements important for backsplicing on both sides of the backspliced junction. The final reporter construct constituted three exons (two exons getting backspliced and a downstream exon to denote the linear splicing with the middle exon) and two introns between them. Classical restriction cloning (Kdm2a, restriction sites-NheI,BamHI) as well as Gibson assembly technique (for Kmt2a, Rad52) were used to create circRNA expression vectors of the corresponding genes Kmt2a (chr9:44,828,270-44,830,122), Kdm2a (chr19:4,319,095-4,322,547) and Rad52 (chr6:119,919,824-119,922,533). Genomic DNA was isolated from Mouse Embryonic fibroblast cells (MEF) using QIAmp DNA mini kit (Qiagen), PCR amplified and subsequently cloned in vector pEGFPC1 with a constitutive promoter. Positive clones were confirmed by sequencing.

Human cervical adenocarcinoma-derived epithelial cell line HeLa (ATCC) and immortalized hTERT-RPE cells were maintained in Dulbecco’s Modified Eagle’s Medium (DMEM, Thermo Fisher), supplemented with 10% fetal bovine serum (FBS, Thermo Fisher), and Penicillin-Streptomycin mix (Thermo Fisher). Cells were transfected using Lipofectamine LTX Reagent or 3000 (Thermo Fisher) according to manufacturer’s protocol, and analyzed 24 hours post plasmid delivery.

Total RNA were isolated by Trizol (Ambion) and treated with DNase. 200ng of RNA was used for complementary DNA (cDNA) production by reverse transcription (Superscript III, ThermoFischer Scientific) using random hexamers and 1/10^th^ dilution of the cDNA. One set of primers was designed in reverse orientation to span the back-spliced junction between 5’ end of exon 1 and 3’ end of exon 2. Circular form of circRNA was confirmed by resistance to the RNaseR (McLab) treatment (10 minutes at 37°C) prior RT-qPCR. The second set of primers span the junction between the 3’ end of exon 2 and the 5’ end of exon 3. This amplifies the linear spliced variant. The fold change expression level (2^-ΔΔCt^) was normalized to normalizing to the negative control (pEGFPC1) and to housekeeping gene (GAPDH) and presented as a ratio of circular/ linear RNA between the wild type and mutant cells.

## Acknowledgements

This work was supported by the Czech Academy of Sciences RVO 68378050 and 68378050-KAV-NPUI and the Czech Science Foundation (20-04099S). M.Kra. was supported by the Czech Academy of Sciences (L200521652), and the Czech Academy of Sciences/Deutscher Akademischer Austauschdienst Mobility Programme (DAAD-17-10). The Czech Centre for Phenogenomics (to D.Z., J.L., A.K.-Z., M.P., J.P., and R.S.) is supported by the project LM2018126 Czech Centre for Phenogenomics provided by MEYS. The additional funding for CCP was provided by projects CZ.02.1.01/0.0/0.0/16_013/0001789 and CZ.02.1.01/0.0/0.0/18_046/0015861 provided by MEYS and ESIF. This work was also supported by ELIXIR CZ research infrastructure project (MEYS Grant No: LM2018131) including access to computing and storage Facilities (to M.M., J.K., M.Ko.). The microscopy images were acquired at the Light Microscopy Core Facility, Institute of Molecular Genetics in Prague, Czech Republic supported by MEYS (LM2015062, CZ.02.1.01/0.0/0.0/16_013/0001775) and OPPK (CZ.2.16/3.1.00/21547).

The authors thank Jana Machatova-Krizova, Sarka Kocourkova and Martina Krausova for excellent technical assistance, Gabriela Vavrova for her careful approach to mouse keeping, and Peter Makovicky for his significant contribution to the histopathological analyses. We would also like to express our gratitude to Assoc.Prof. Jakub Otahal from Institute of Physiology of the Czech Academy of Sciences for his help with assignment of stereotaxic coordinates used for description of histopathological sections. We would also like to thank to dr. Trevor Epp from Institute of Molecular Genetics of the Czech Academy of Sciences for helpful discussion on the transmission ratio distortion.

## Declarations

### Ethical approval

All animal procedures were conducted following the European Union guidelines (regulation n°86/609), the Czech law regulating animal experimentation (246/1992 Sb.) and approved by the ethics committee of Czech Republic (reference number 31255/2019-MZE-18134, file ID 16OZ9707/2019-18134, valid till July 2nd, 2024).

### Data availability

The datasets generated and/or analysed during the current study are available in the:

- RNA-Seq data: Array Express accession E-MTAB-10753 (www.ebi.ac.uk/arrayexpress/experiments/E-MTAB-10753 “\t”_blank)
- RT-qPCR linear model (github.com/cas-bioinf/prpf8)

### Author contributions

M.Kra. performed most of the experiments and together with D.S. conceived and designed the project; M.Kra. and R.S. generated the mouse strains; M.Kra., D.Z., J.L., A.K.-Z., and M.P. analyzed various parameters in the mice; M.Kre analyzed splicing factor expression and performed immunoprecipitations; P.B. created circRNA reporters and analyzed their expression; J.K. performed the transcriptomic analysis; M. Kra. and M.M. analyzed the RT-qPCR data; J.P., M.Ko., R.S., and D.S. supervised experiments; M.Kra. and D.S. wrote the manuscript.

## References

Ahmad-Annuar, A., Ciani, L., Simeonidis, I., Herreros, J., Fredj, N. B., Rosso, S. B., Hall, A., Brickley, S. and Salinas, P. C. (2006). Signaling across the synapse: a role for Wnt and Dishevelled in presynaptic assembly and neurotransmitter release. J Cell Biol 174, 127–39.

Ashwal-Fluss, R., Meyer, M., Pamudurti, N. R., Ivanov, A., Bartok, O., Hanan, M., Evantal, N., Memczak, S., Rajewsky, N. and Kadener, S. (2014). CircRNA Biogenesis competes with Pre-mRNA splicing. Mol Cell 56, 55–66.

Audo, I., Bujakowska, K., Mohand-Said, S., Lancelot, M. E., Moskova-Doumanova, V., Waseem, N. H., Antonio, A., Sahel, J. A., Bhattacharya, S. S. and Zeitz, C. (2010). Prevalence and novelty of PRPF31 mutations in French autosomal dominant rod-cone dystrophy patients and a review of published reports. BMC Med Genet 11, 145.

Buskin, A., Zhu, L., Chichagova, V., Basu, B., Mozaffari-Jovin, S., Dolan, D., Droop, A., Collin, J., Bronstein, R., Mehrotra, S. et al. (2018). Disrupted alternative splicing for genes implicated in splicing and ciliogenesis causes PRPF31 retinitis pigmentosa. Nat Commun 9, 4234.

Cao, H., Wu, J., Lam, S., Duan, R., Newnham, C., Molday, R. S., Graziotto, J. J., Pierce, E. A. and Hu, J. (2011). Temporal and tissue specific regulation of RP-associated splicing factor genes PRPF3, PRPF31 and PRPC8--implications in the pathogenesis of RP. PLoS ONE 6, e15860.

Chakarova, C. F., Hims, M. M., Bolz, H., Abu-Safieh, L., Patel, R. J., Papaioannou, M. G., Inglehearn, C. F., Keen, T. J., Willis, C., Moore, A. T. et al. (2002). Mutations in HPRP3, a third member of pre-mRNA splicing factor genes, implicated in autosomal dominant retinitis pigmentosa. Hum Mol Genet 11, 87–92.

Chen, X., Liu, Y., Sheng, X., Tam, P. O., Zhao, K., Chen, X., Rong, W., Liu, Y., Liu, X., Pan, X. et al. (2014). PRPF4 mutations cause autosomal dominant retinitis pigmentosa. Hum Mol Genet 23, 2926–39.

Cong, L., Ran, F. A., Cox, D., Lin, S., Barretto, R., Habib, N., Hsu, P. D., Wu, X., Jiang, W., Marraffini, L. A. et al. (2013). Multiplex genome engineering using CRISPR/Cas systems. Science 339, 819–23.

Cvackova, Z., Mateju, D. and Stanek, D. (2014). Retinitis Pigmentosa Mutations of SNRNP200 Enhance Cryptic Splice-Site Recognition. Hum Mutat 35, 308–17.

De Erkenez, A. C., Berson, E. L. and Dryja, T. P. (2002). Novel Mutations in the PRPC8 Gene, Encoding a Pre-mRNA Splicing Factor in Patients with Autosomal Dominant Retinitis Pigmentosa. Invest Ophthalmol Vis Sci 43, 791.

Divya, T. S., Lalitha, S., Parvathy, S., Subashini, C., Sanalkumar, R., Dhanesh, S. B., Rasheed, V. A., Divya, M. S., Tole, S. and James, J. (2016). Regulation of Tlx3 by Pax6 is required for the restricted expression of Chrnalpha3 in Cerebellar Granule Neuron progenitors during development. Sci Rep 6, 30337.

Doyle, E. L., Booher, N. J., Standage, D. S., Voytas, D. F., Brendel, V. P., Vandyk, J. K. and Bogdanove, A. J. (2012). TAL Effector-Nucleotide Targeter (TALE-NT) 2.0: tools for TAL effector design and target prediction. Nucleic Acids Res 40, W117–22.

Farkas, M. H., Lew, D. S., Sousa, M. E., Bujakowska, K., Chatagnon, J., Bhattacharya, S. S., Pierce, E. A. and Nandrot, E. F. (2014). Mutations in pre-mRNA processing factors 3, 8, and 31 cause dysfunction of the retinal pigment epithelium. Am J Pathol 184, 2641–52.

Gao, Y., Zhang, J. and Zhao, F. (2018). Circular RNA identification based on multiple seed matching. Brief Bioinform 19, 803–810.

Gonzalez-Santos, J. M., Cao, H., Duan, R. C. and Hu, J. (2008). Mutation in the splicing factor Hprp3p linked to retinitis pigmentosa impairs interactions within the U4/U6 snRNP complex. Hum Mol Genet 17, 225–39.

Graziotto, J. J., Farkas, M. H., Bujakowska, K., Deramaudt, B. M., Zhang, Q., Nandrot, E. F., Inglehearn, C. F., Bhattacharya, S. S. and Pierce, E. A. (2011). Three gene-targeted mouse models of RNA splicing factor RP show late-onset RPE and retinal degeneration. Invest Ophthalmol Vis Sci 52, 190–8.

Hansen, T. B., Jensen, T. I., Clausen, B. H., Bramsen, J. B., Finsen, B., Damgaard, C. K. and Kjems, J. (2013). Natural RNA circles function as efficient microRNA sponges. Nature 495, 384–8.

Huranova, M., Hnilicova, J., Fleischer, B., Cvackova, Z. and Stanek, D. (2009). A mutation linked to retinitis pigmentosa in HPRP31 causes protein instability and impairs its interactions with spliceosomal snRNPs. Hum Mol Genet 18, 2014–2023.

Izuogu, O. G., Alhasan, A. A., Mellough, C., Collin, J., Gallon, R., Hyslop, J., Mastrorosa, F. K., Ehrmann, I., Lako, M., Elliott, D. J. et al. (2018). Analysis of human ES cell differentiation establishes that the dominant isoforms of the lncRNAs RMST and FIRRE are circular. BMC Genomics 19, 276.

Johnson, S., Halford, S., Morris, A. G., Patel, R. J., Wilkie, S. E., Hardcastle, A. J., Moore, A. T., Zhang, K. and Hunt, D. M. (2003). Genomic organisation and alternative splicing of human RIM1, a gene implicated in autosomal dominant cone-rod dystrophy (CORD7). Genomics 81, 304–14.

Kano, M. and Watanabe, T. (2019). Developmental synapse remodeling in the cerebellum and visual thalamus. F1000Res 8.

Kasparek, P., Krausova, M., Haneckova, R., Kriz, V., Zbodakova, O., Korinek, V. and Sedlacek, R. (2014). Efficient gene targeting of the Rosa26 locus in mouse zygotes using TALE nucleases. FEBS Lett 588, 3982–8.

Kim, D., Langmead, B. and Salzberg, S. L. (2015). HISAT: a fast spliced aligner with low memory requirements. Nat Methods 12, 357–60.

Kleaveland, B., Shi, C. Y., Stefano, J. and Bartel, D. P. (2018). A Network of Noncoding Regulatory RNAs Acts in the Mammalian Brain. Cell 174, 350–362 e17.

Krausova, M. and Stanek, D. (2018). snRNP proteins in health and disease. Semin Cell Dev Biol 79, 92–102.

Krchnakova, Z., Thakur, P. K., Krausova, M., Bieberstein, N., Haberman, N., Muller-McNicoll, M. and Stanek, D. (2019). Splicing of long non-coding RNAs primarily depends on polypyrimidine tract and 5’ splice-site sequences due to weak interactions with SR proteins. Nucleic Acids Res 47, 911–928.

Kukhtar, D., Rubio-Pena, K., Serrat, X. and Ceron, J. (2020). Mimicking of splicing-related retinitis pigmentosa mutations in C. elegans allow drug screens and identification of disease modifiers. Hum Mol Genet 29, 756–765.

Liang, D., Tatomer, D. C., Luo, Z., Wu, H., Yang, L., Chen, L. L., Cherry, S. and Wilusz, J. E. (2017). The Output of Protein-Coding Genes Shifts to Circular RNAs When the Pre-mRNA Processing Machinery Is Limiting. Mol Cell 68, 940–954 e3.

Liao, Y., Smyth, G. K. and Shi, W. (2014). featureCounts: an efficient general purpose program for assigning sequence reads to genomic features. Bioinformatics 30, 923–30.

Linder, B., Hirmer, A., Gal, A., Ruther, K., Bolz, H. J., Winkler, C., Laggerbauer, B. and Fischer, U. (2014). Identification of a PRPF4 loss-of-function variant that abrogates U4/U6.U5 tri-snRNP integration and is associated with retinitis pigmentosa. PLoS ONE 9, e111754.

Liu, X., Liu, B., Zhou, M., Fan, F., Yu, M., Gao, C., Lu, Y. and Luo, Y. (2018). Circular RNA HIPK3 regulates human lens epithelial cells proliferation and apoptosis by targeting the miR-193a/CRYAA axis. Biochem Biophys Res Commun 503, 2277–2285.

Love, M. I., Huber, W. and Anders, S. (2014). Moderated estimation of fold change and dispersion for RNA-seq data with DESeq2. Genome Biol 15, 550.

Malinova, A., Cvackova, Z., Mateju, D., Horejsi, Z., Abeza, C., Vandermoere, F., Bertrand, E., Stanek, D. and Verheggen, C. (2017). Assembly of the U5 snRNP component PRPF8 is controlled by the HSP90/R2TP chaperones. J Cell Biol 216, 1579–1596.

Mancini, E., Rabinovich, A., Iserte, J., Yanovsky, M. and Chernomoretz, A. (2021). Corrigendum to: ASpli: Integrative analysis of splicing landscapes through RNA-Seq assays. Bioinformatics.

Martinez-Gimeno, M., Gamundi, M. J., Hernan, I., Maseras, M., Milla, E., Ayuso, C., Garcia-Sandoval, B., Beneyto, M., Vilela, C., Baiget, M. et al. (2003). Mutations in the pre-mRNA splicing-factor genes PRPF3, PRPF8, and PRPF31 in Spanish families with autosomal dominant retinitis pigmentosa. Invest Ophthalmol Vis Sci 44, 2171–7.

Mattapallil, M. J., Wawrousek, E. F., Chan, C. C., Zhao, H., Roychoudhury, J., Ferguson, T. A. and Caspi, R. R. (2012). The Rd8 mutation of the Crb1 gene is present in vendor lines of C57BL/6N mice and embryonic stem cells, and confounds ocular induced mutant phenotypes. Invest Ophthalmol Vis Sci 53, 2921–7.

Matz, M. V., Wright, R. M. and Scott, J. G. (2013). No control genes required: Bayesian analysis of qRT-PCR data. PLoS ONE 8, e71448.

McCarthy, D. J., Chen, Y. and Smyth, G. K. (2012). Differential expression analysis of multifactor RNA-Seq experiments with respect to biological variation. Nucleic Acids Res 40, 4288–97.

McKie, A. B., McHale, J. C., Keen, T. J., Tarttelin, E. E., Goliath, R., van Lith-Verhoeven, J. J., Greenberg, J., Ramesar, R. S., Hoyng, C. B., Cremers, F. P. et al. (2001). Mutations in the pre-mRNA splicing factor gene PRPC8 in autosomal dominant retinitis pigmentosa (RP13). Human Molecular Genetics 10, 1555–1562.

Mechaussier, S., Almoallem, B., Zeitz, C., Van Schil, K., Jeddawi, L., Van Dorpe, J., Duenas Rey, A., Condroyer, C., Pelle, O., Polak, M. et al. (2020). Loss of Function of RIMS2 Causes a Syndromic Congenital Cone-Rod Synaptic Disease with Neurodevelopmental and Pancreatic Involvement. Am J Hum Genet 106, 859–871.

Mellough, C. B., Bauer, R., Collin, J., Dorgau, B., Zerti, D., Dolan, D. W. P., Jones, C. M., Izuogu, O. G., Yu, M., Hallam, D. et al. (2019). An integrated transcriptional analysis of the developing human retina. Development 146.

Mordes, D., Yuan, L., Xu, L., Kawada, M., Molday, R. S. and Wu, J. Y. (2007). Identification of photoreceptor genes affected by PRPF31 mutations associated with autosomal dominant retinitis pigmentosa. Neurobiol Dis 26, 291–300.

Mozaffari-Jovin, S., Wandersleben, T., Santos, K. F., Will, C. L., Luhrmann, R. and Wahl, M. C. (2013). Inhibition of RNA helicase Brr2 by the C-terminal tail of the spliceosomal protein Prp8. Science 341, 80–4.

Mozaffari-Jovin, S., Wandersleben, T., Santos, K. F., Will, C. L., Luhrmann, R. and Wahl, M. C. (2014). Novel regulatory principles of the spliceosomal Brr2 RNA helicase and links to retinal disease in humans. RNA Biol 11, 298–312.

Piwecka, M., Glazar, P., Hernandez-Miranda, L. R., Memczak, S., Wolf, S. A., Rybak-Wolf, A., Filipchyk, A., Klironomos, F., Cerda Jara, C. A., Fenske, P. et al. (2017). Loss of a mammalian circular RNA locus causes miRNA deregulation and affects brain function. Science 357.

Rahimi, K., Veno, M. T., Dupont, D. M. and Kjems, J. (2021). Nanopore sequencing of brain-derived full-length circRNAs reveals circRNA-specific exon usage, intron retention and microexons. Nat Commun 12, 4825.

Ran, F. A., Hsu, P. D., Wright, J., Agarwala, V., Scott, D. A. and Zhang, F. (2013). Genome engineering using the CRISPR-Cas9 system. Nat Protoc 8, 2281–2308.

Ray, P., Luo, X., Rao, E. J., Basha, A., Woodruff, E. A., 3rd and Wu, J. Y. (2010). The splicing factor Prp31 is essential for photoreceptor development in Drosophila. Protein Cell 1, 267–74.

Rio Frio, T., Wade, N. M., Ransijn, A., Berson, E. L., Beckmann, J. S. and Rivolta, C. (2008). Premature termination codons in PRPF31 cause retinitis pigmentosa via haploinsufficiency due to nonsense-mediated mRNA decay. J Clin Invest 118, 1519–31.

Ruzickova, S. and Stanek, D. (2017). Mutations in spliceosomal proteins and retina degeneration. RNA Biol 14, 544–552.

Rybak-Wolf, A., Stottmeister, C., Glazar, P., Jens, M., Pino, N., Giusti, S., Hanan, M., Behm, M., Bartok, O., Ashwal-Fluss, R. et al. (2015). Circular RNAs in the Mammalian Brain Are Highly Abundant, Conserved, and Dynamically Expressed. Mol Cell 58, 870–85.

Saunders, A., Macosko, E. Z., Wysoker, A., Goldman, M., Krienen, F. M., de Rivera, H., Bien, E., Baum, M., Bortolin, L., Wang, S. et al. (2018). Molecular Diversity and Specializations among the Cells of the Adult Mouse Brain. Cell 174, 1015–1030 e16.

Stankovic, D., Claudius, A. K., Schertel, T., Bresser, T. and Uhlirova, M. (2020). A Drosophila model to study retinitis pigmentosa pathology associated with mutations in the core splicing factor Prp8. Dis Model Mech 13.

Starke, S., Jost, I., Rossbach, O., Schneider, T., Schreiner, S., Hung, L. H. and Bindereif, A. (2015). Exon circularization requires canonical splice signals. Cell Rep 10, 103–11.

Steibel, J. P., Poletto, R., Coussens, P. M. and Rosa, G. J. (2009). A powerful and flexible linear mixed model framework for the analysis of relative quantification RT-PCR data. Genomics 94, 146–52.

Stephens, M. (2017). False discovery rates: a new deal. Biostatistics 18, 275–294.

Sullivan, L. S., Bowne, S. J., Seaman, C. R., Blanton, S. H., Lewis, R. A., Heckenlively, J. R., Birch, D. G., Hughbanks-Wheaton, D. and Daiger, S. P. (2006). Genomic rearrangements of the PRPF31 gene account for 2.5% of autosomal dominant retinitis pigmentosa. Invest Ophthalmol Vis Sci 47, 4579–88.

Sun, L. F., Ma, Y., Ji, Y. Y., Wu, Z., Wang, Y. H., Mou, H. and Jin, Z. B. (2021). Circular Rims2 Deficiency Causes Retinal Degeneration. Adv Biol (Weinh), e2100906.

Sun, L. F., Zhang, B., Chen, X. J., Wang, X. Y., Zhang, B. W., Ji, Y. Y., Wu, K. C., Wu, J. and Jin, Z. B. (2019). Circular RNAs in human and vertebrate neural retinas. RNA Biol 16, 821–829.

Tanackovic, G., Ransijn, A., Ayuso, C., Harper, S., Berson, E. L. and Rivolta, C. (2011a). A missense mutation in PRPF6 causes impairment of pre-mRNA splicing and autosomal-dominant retinitis pigmentosa. Am J Hum Genet 88, 643–9.

Tanackovic, G., Ransijn, A., Thibault, P., Abou Elela, S., Klinck, R., Berson, E. L., Chabot, B. and Rivolta, C. (2011b). PRPF mutations are associated with generalized defects in spliceosome formation and pre-mRNA splicing in patients with retinitis pigmentosa. Hum Mol Genet 20, 2116–30.

Tiwari, A., Lemke, J., Altmueller, J., Thiele, H., Glaus, E., Fleischhauer, J., Nurnberg, P., Neidhardt, J. and Berger, W. (2016). Identification of Novel and Recurrent Disease-Causing Mutations in Retinal Dystrophies Using Whole Exome Sequencing (WES): Benefits and Limitations. PLoS ONE 11, e0158692.

Towns, K. V., Kipioti, A., Long, V., McKibbin, M., Maubaret, C., Vaclavik, V., Ehsani, P., Springell, K., Kamal, M., Ramesar, R. S. et al. (2010). Prognosis for splicing factor PRPF8 retinitis pigmentosa, novel mutations and correlation between human and yeast phenotypes. Hum Mutat 31, E1361–76.

Visscher, P. M. (1999). Speed congenics: accelerated genome recovery using genetic markers. Genet Res 74, 81–5.

Vithana, E. N., Abu-Safieh, L., Allen, M. J., Carey, A., Papaioannou, M., Chakarova, C., Al-Maghtheh, M., Ebenezer, N. D., Willis, C., Moore, A. T. et al. (2001). A human homolog of yeast pre-mRNA splicing gene, PRP31, underlies autosomal dominant retinitis pigmentosa on chromosome 19q13.4 (RP11). Molecular Cell 8, 375–381.

Vithana, E. N., Abu-Safieh, L., Pelosini, L., Winchester, E., Hornan, D., Bird, A. C., Hunt, D. M., Bustin, S. A. and Bhattacharya, S. S. (2003). Expression of PRPF31 mRNA in patients with autosomal dominant retinitis pigmentosa: a molecular clue for incomplete penetrance? Invest Ophthalmol Vis Sci 44, 4204–9.

Wang, M., Hou, J., Muller-McNicoll, M., Chen, W. and Schuman, E. M. (2019). Long and Repeat-Rich Intronic Sequences Favor Circular RNA Formation under Conditions of Reduced Spliceosome Activity. iScience 20, 237–247.

Wang, Y., Okamoto, M., Schmitz, F., Hofmann, K. and Sudhof, T. C. (1997). Rim is a putative Rab3 effector in regulating synaptic-vesicle fusion. Nature 388, 593–8.

Yin, J., Brocher, J., Fischer, U. and Winkler, C. (2011). Mutant Prpf31 causes pre-mRNA splicing defects and rod photoreceptor cell degeneration in a zebrafish model for Retinitis pigmentosa. Mol Neurodegener 6, 56.

You, X., Vlatkovic, I., Babic, A., Will, T., Epstein, I., Tushev, G., Akbalik, G., Wang, M., Glock, C., Quedenau, C. et al. (2015). Neural circular RNAs are derived from synaptic genes and regulated by development and plasticity. Nat Neurosci 18, 603–610.

Young, M. D., Wakefield, M. J., Smyth, G. K. and Oshlack, A. (2010). Gene ontology analysis for RNA-seq: accounting for selection bias. Genome Biol 11, R14.

Yu, S., Li, C., Biswas, L., Hu, X., Liu, F., Reilly, J., Liu, X., Liu, Y., Huang, Y., Lu, Z. et al. (2017). CERKL gene knockout disturbs photoreceptor outer segment phagocytosis and causes rod-cone dystrophy in zebrafish. Hum Mol Genet 26, 2335–2345.

Yuan, L., Kawada, M., Havlioglu, N., Tang, H. and Wu, J. Y. (2005). Mutations in PRPF31 inhibit pre-mRNA splicing of rhodopsin gene and cause apoptosis of retinal cells. J Neurosci 25, 748–57.

Zhang, J., Chen, S., Yang, J. and Zhao, F. (2020). Accurate quantification of circular RNAs identifies extensive circular isoform switching events. Nat Commun 11, 90.

Zhao, C., Bellur, D. L., Lu, S., Zhao, F., Grassi, M. A., Bowne, S. J., Sullivan, L. S., Daiger, S. P., Chen, L. J., Pang, C. P. et al. (2009). Autosomal-dominant retinitis pigmentosa caused by a mutation in SNRNP200, a gene required for unwinding of U4/U6 snRNAs. Am J Hum Genet 85, 617–27.

