## Supplementary figures for "Retinitis pigmentosa associated mutations in mouse Prpf8 cause misexpression of circRNAs and degeneration of cerebellar granule neurons"

Figure S1

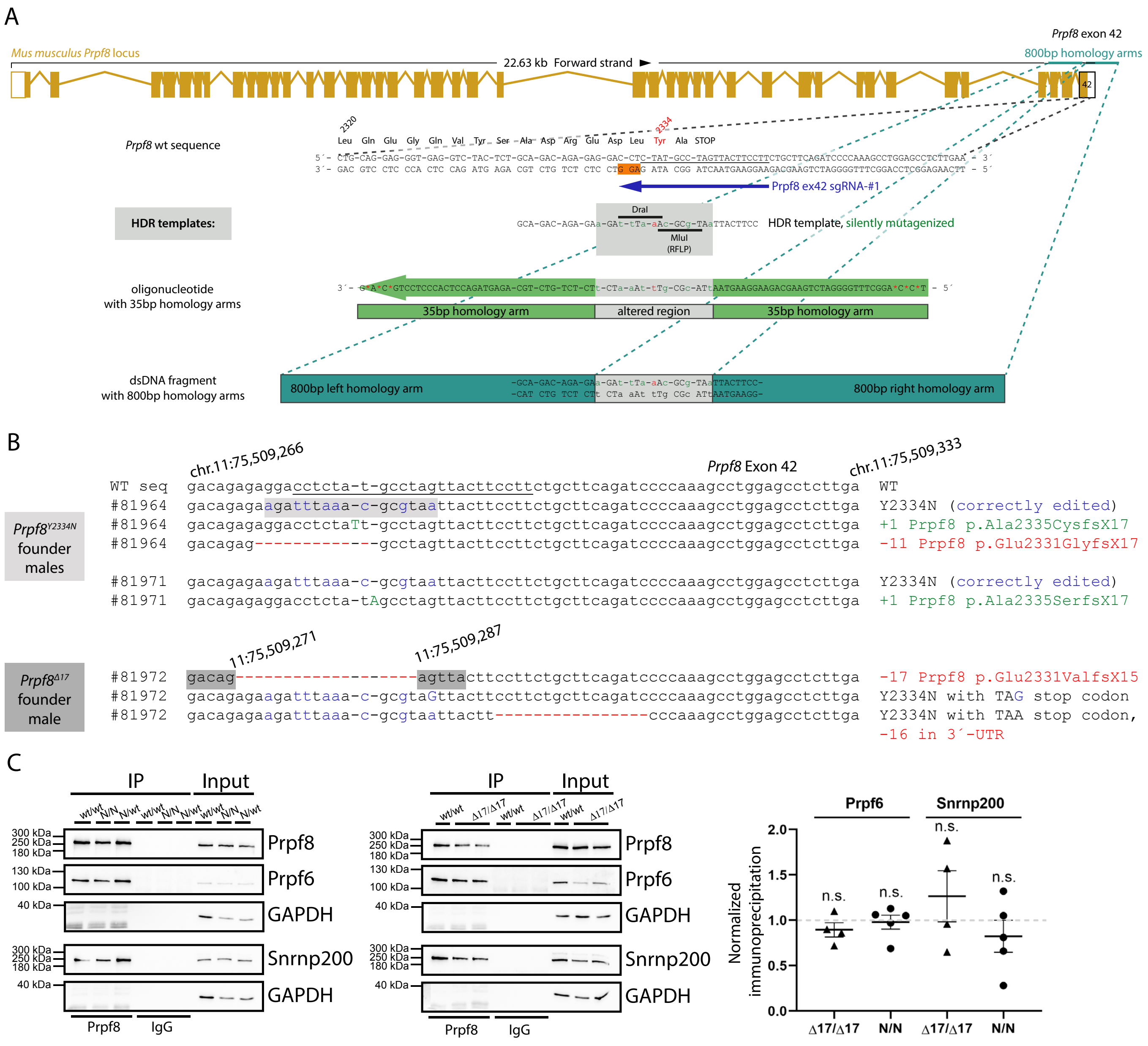


**Figure S1. Design and in vitro testing of the CRISPR/Cas9 editing tools used to introduce the aberrant Prpf8 mutations into murine gDNA.** (A) Schematic representation of the mouse Prpf8 gene with depiction of the targeted exon 42, with partial amino acid translation provided below the locus diagram. The CRISPR binding site (sgRNA: Prpf8 ex42 sgRNA-#1; dark blue arrow) is underlined in black, the protospacer adjacent motif (PAM) is highlighted in the non-target DNA strand with an orange box. A synthetic oligodeoxynucleotide (visualized as green arrow) or double-stranded DNA fragment (in teal) were employed in parallel as HDR donors. (B) Sequencing of the targeted Prpf8 exon 42 in Prpf8 founder mice. In Prpf8Y2334N founders, apart from the wt and template-edited alleles (depicted in blue) we recorded additional small InDels (highlighted in green and red). In Prpf8Δ17 founder, we detected diverse small-scale deletions (in red), that size-matched the additional MluI RFLP products. The CRISPR binding site including the sequence corresponding to the protospacer adjacent motif (PAM) is underlined in black. (C) Prpf8 was immunoprecipitated from cerebellum of 4 weeks old animals of the Prpf8Y2334N and Prpf8Δ17 strains and co-precipitation of U5 snRNP-specific proteins Snrnp200 and Prpf6 was detected by Western blotting. GAPDH was used as a loading control, and immunoprecipitation with control IgG served as a negative control. Snrnp200 and Prpf6 protein levels from four experiments were normalized to levels of Prpf8 and plotted. Prpf8Y2334N/Y2334N and Prpf8Δ17/Δ17 mice did not show statistically significant differences in protein levels, when compared to Prpf8wt/wt controls. Values presented in graph were normalized to Prpf8wt/wt controls. Statistical analysis was performed by using nonparametric t-test in GraphPad software and is displayed as mean with SEM.

Figure S2

A

| Genotype of offspring | *Δ17/Δ17* | *Δ17/wt* | *wt/wt* | Total | P |
|---|---|---|---|---|---|
| Observed number of mice | 46 | 113 | 50 | 209 | 0.464 |
| Expected number of mice (ratio 1:2:1) | 52.3 | 104.5 | 52.3 | | |

B

| Genotype of offspring | *Y2334N/Y2334N* | *Y2334N/wt* | *wt/wt* | Total | P |
|---|---|---|---|---|---|
| Observed number of mice | 49 | 81 | 44 | 174 | 0.573 |
| Expected number of mice (ratio 1:2:1) | 43.5 | 87 | 43.5 | | |

C

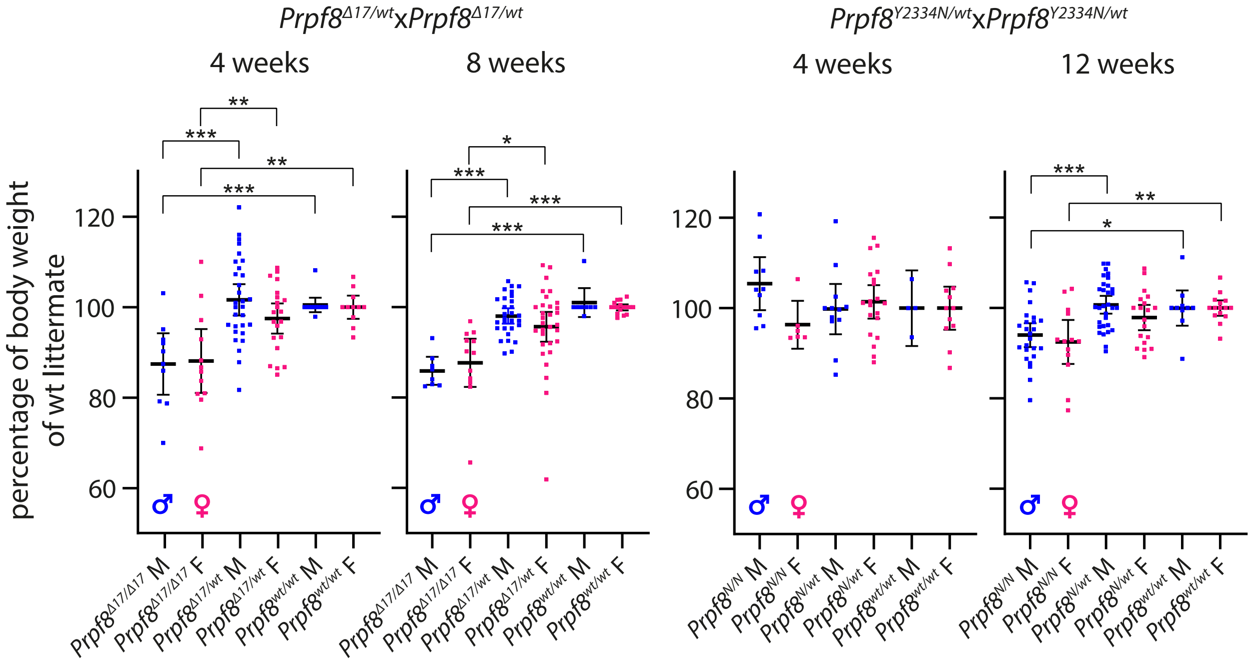


**Figure S2. Analysis of mice expressing Prpf8 variants.** (A) A total of 209 mice born from Prpf8Δ17/wt breedings is shown; the expected number of offspring was calculated according to the total number of mice born and based on the anticipated Mendelian 1:2:1 ratio. (B) A total number of 174 progeny was analyzed from Prpf8Y2334N/wt crossbreedings. (C) Growth of animals born to heterozygotic breeding pairs. Body weight of sacrificed animals was measured at 4 and 8/12 weeks. Actual body weight was normalized to median of Prpf8wt/wt peers of matching gender in individual litter. Statistical significance of differential body weight in the given genotype groups was examined by t-test using GraphPad software (solid line; * P < 0.05, ** P < 0.01, *** P < 0.001; displayed is mean with 95% confidentiality interval). Males (M) in blue, females (F) in pink.

Figure S3

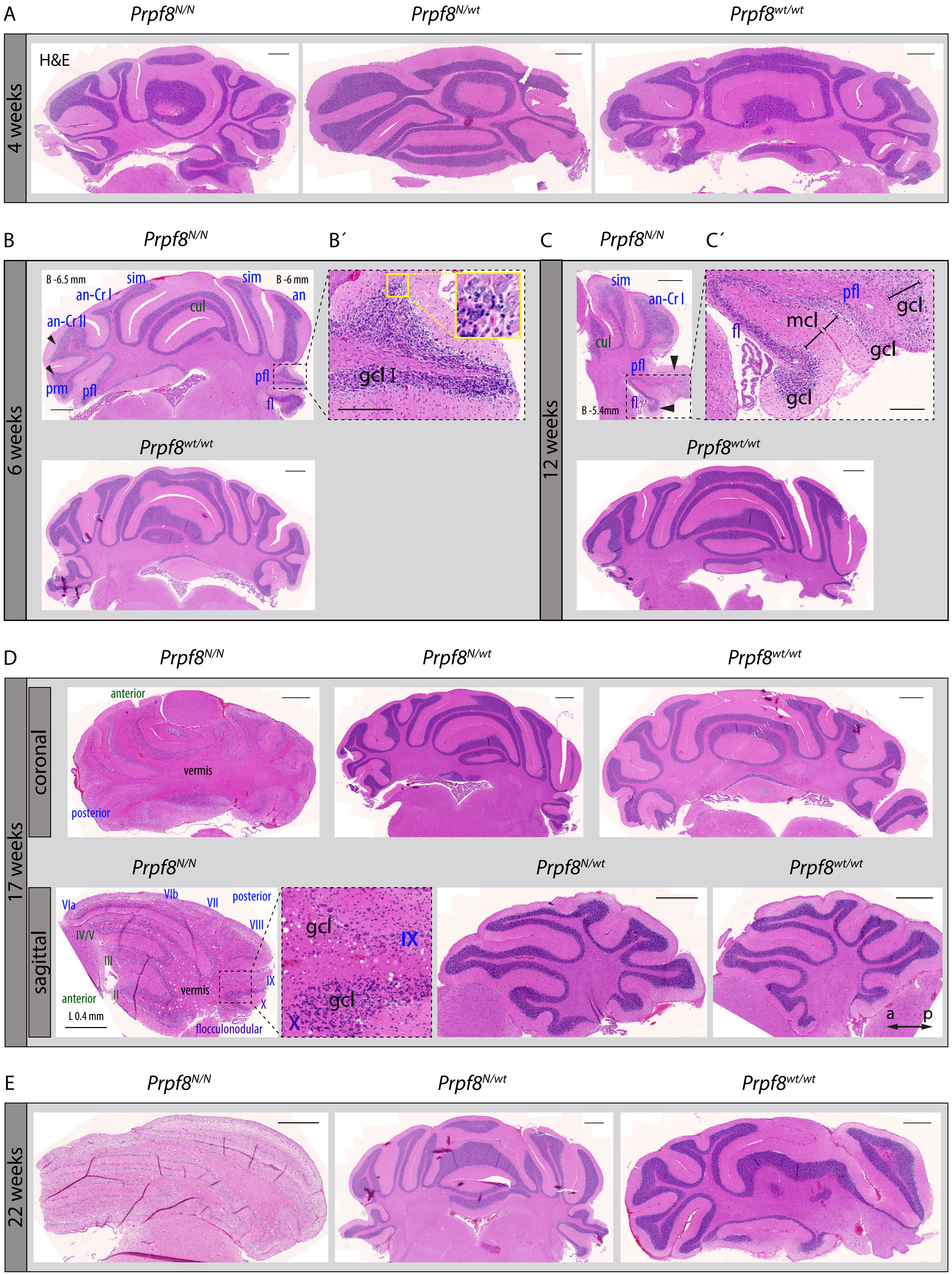

A
Prpf8N/N
Prpf8N/wt
Prpf8wt/wt
4 weeks
H&E
B
Prpf8N/N
B´
C
Prpf8N/N
C´
6 weeks
B -6.5 mm
sim
an-Cr I
an-Cr II
cul
prm
pfl
B -6 mm
an
fl
gcl
Prpf8wt/wt
12 weeks
an-Cr I
mcl
B -5.4mm
D
Prpf8N/N
Prpf8N/wt
Prpf8wt/wt
17 weeks
coronal
anterior
vermis
posterior
sagittal
VIa
VIb
VII
VIII
IV/V
III
II
IX
X
L 0.4 mm
flocculonodular
a
p
E
Prpf8N/N
Prpf8N/wt
Prpf8wt/wt
22 weeks

**Figure S3. Pilot histopathological examination revealed progressive degeneration of the cerebellar granule cell layer in aging homozygous *Prpf8*$^{Y2334N/Y2334N}$ mice.** (*A*) No pathological alterations were macroscopically detectable by 4 weeks of age in the cerebellum of homozygous *Prpf8*$^{Y2334N/Y2334N}$ (*Prpf8*$^{N/N}$) mice. (*B*) At 6 weeks of age, thinning in the granule cell layer (gcl) became noticeable in the posterior lobe (black arrowheads; lobule description ansiform Crus I (an-Cr I), ansiform Crus II (an-Cr II), culmen (cul), flocculus (fl), paramedian (prm), paraflocculus (pfl), simplex (sim)). Naming of lobules belonging to the anterior lobe is depicted in green, posterior in blue, and flocculonodular in violet. (*B´*) A detail from the apex of the paraflocculus lobule shows presence of shrunken granule cell nuclei (yellow inset). (*C*) Apparent difference (black arrowheads) in the cellular density in the stratum granulosum of the paraflocculus (pfl) and the adjoining flocculus (fl) lobules reasserted in animals aged to 12 weeks; a detail from (*C*) is provided in (*C´*). gcl - granule cell layer, mcl - molecular cell layer. (*D*) The granular layer content was severely reduced in 17 weeks old animals, with posterior lobe being more severely affected compared to the anterior one, and apical segments of lobules being more afflicted compared to base regions enclosed to vermis (top row coronal sections, bottom row sagittal sections; Roman numerals depict cerebellum lobules; a = anterior, p = posterior for sagittal sections). (*E*) Complete atrophy of the cerebellar granular layer by 22 weeks of age. H&E = hematoxylin and eosin staining. Scale bar = 500 µm, in (*B´*) and (*C´*) scale bar set to 200 µm. Approximate stereotaxic coordinates were inferred from Paxinos & Franklin´s Mouse Brain Atlas (2) and from reference Allen Brain Atlas (http://mouse.brain-map.org), and are provided in (*B*) and (*C*) (B - Bregma, coronal sections), and (*D*) (L - lateral, sagittal section) to indicate tilting of the paraffin block during sectioning.

Figure S4

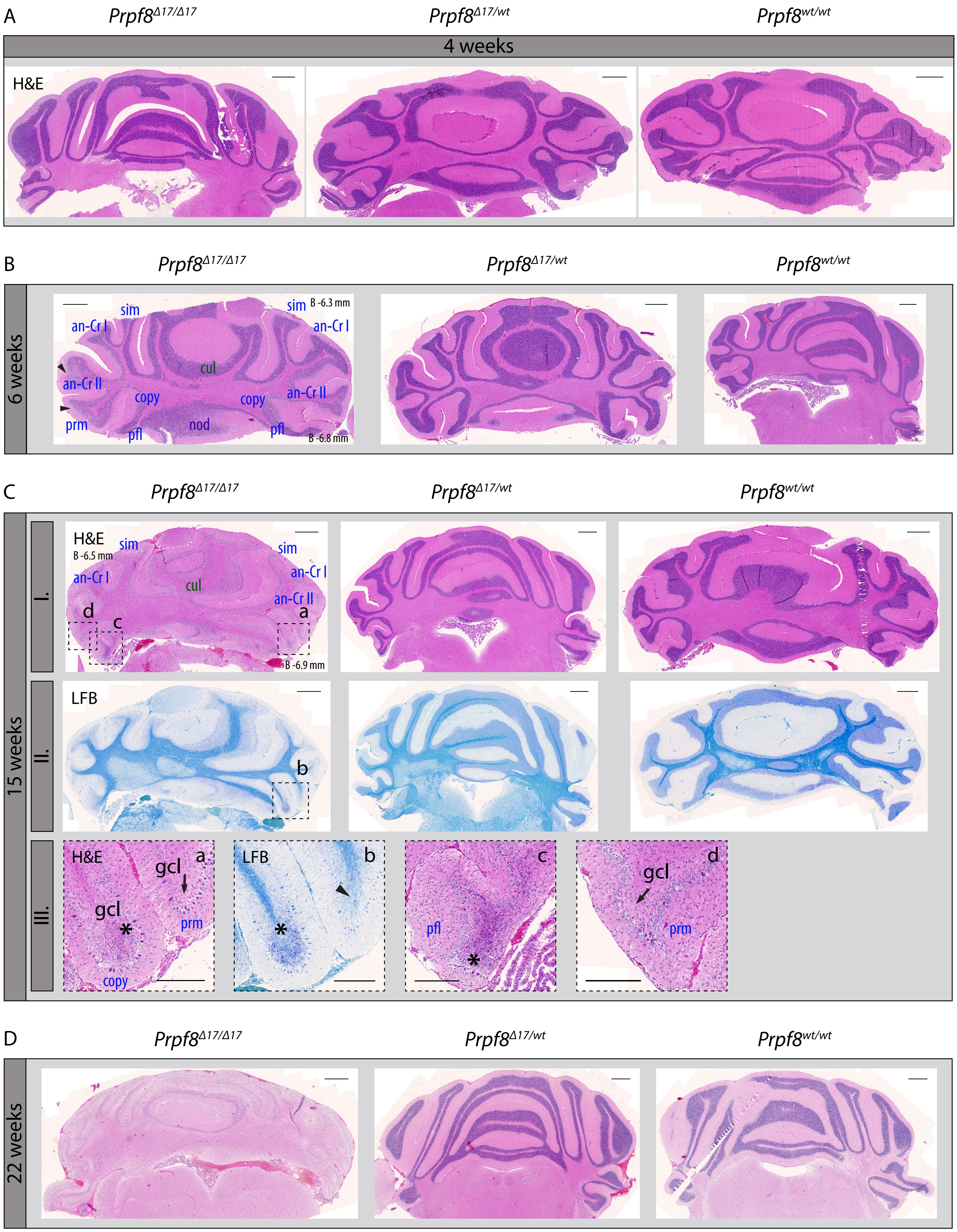

A
Prprf8Δ17/Δ17
Prpf8Δ17/wt
Prpf8wt/wt
4 weeks
H&E
B
Prpf8Δ17/Δ17
Prpf8Δ17/wt
Prpf8wt/wt
6 weeks
sim
an-Cr I
an-Cr II
cul
copy
prm
pfl
nod
B -6.3 mm
B -6.8 mm
C
Prpf8Δ17/Δ17
Prpf8Δ17/wt
Prpf8wt/wt
15 weeks
I.
II.
III.
H&E
B -6.5 mm
B -6.9 mm
LFB
gcl
a
b
c
d
D
Prpf8Δ17/Δ17
Prpf8Δ17/wt
Prpf8wt/wt
22 weeks

**Figure S4. Pilot histopathological examination revealed progressive degeneration of the cerebellar granule cell layer in aging homozygous *Prpf8*$^{\Delta17/\Delta17}$ mice.** (*A*) At 4 weeks of age, no signs of pathological changes were macroscopically detectable in the cerebellum of homozygous *Prpf8*$^{\Delta17/\Delta17}$ animals. (*B*) Two weeks later, thinning of the granule cell layer becomes noticeable in apical parts of posterior lobe lobules (black arrowheads; lobule description ansiform Crus I (an-Cr I), ansiform Crus II (an-Cr II), copula pyramidis (copy), culmen (cul), flocculus (fl), paramedian (prm), paraflocculus (pfl), simplex (sim)). (*C*) By 15 weeks of age, the granular layer content was severely reduced in homozygous individuals. Top row (I.) are sections stained with hematoxylin and eosin dyes (H&E), middle row (II.) represent near neighboring sections stained with Luxol Fast Blue (LFB). Bottom row (III.) are details derived from the 15 weeks old *Prpf8*$^{\Delta17/\Delta17}$ mouse, original locations are indicated by dashed line in row I. (H&E) and row II. (LFB) images, respectively. (III.-a; H&E) and (III.-b; LFB) depict the apical portions of copula pyramidis (copy) and paramedian (prm) lobules, while paraflocculus (pfl) and paramedian (prm) are shown in (III.-c) and (III.-d), respectively. In apical segments of copula pyramidis and paraflocculus lobules we noticed presence of shrunken and disintegrating granule cell nuclei (* in III.-a, III.-b, and III.-c) that likely indicate ongoing neural apoptosis in the granule cell layer (gcl), while in the adjoining paramedian lobule the granule cell layer has already almost been abolished (black arrows in (III.-a and III.-d)). Moreover, in the apical region of paramedian the LFB positivity (III.-b) was limited to myelin sheaths of individual axons of Purkinje cells (black arrowhead in (III.-b)). (*D*) Complete atrophy of the cerebellar granule layer by 22 weeks of age. Scale bar = 500 µm, in (*C*)-III. a-d insets scale bar is set to 200 µm. Approximate stereotaxic coordinates were inferred from Paxinos & Franklin´s Mouse Brain Atlas and Allen Brain Atlas (http://mouse.brain-map.org) and are provided in (*B*) and (*C-I.*) (B - Bregma, coronal sections) to indicate tilting of the paraffin block on sectioning.

Figure S5

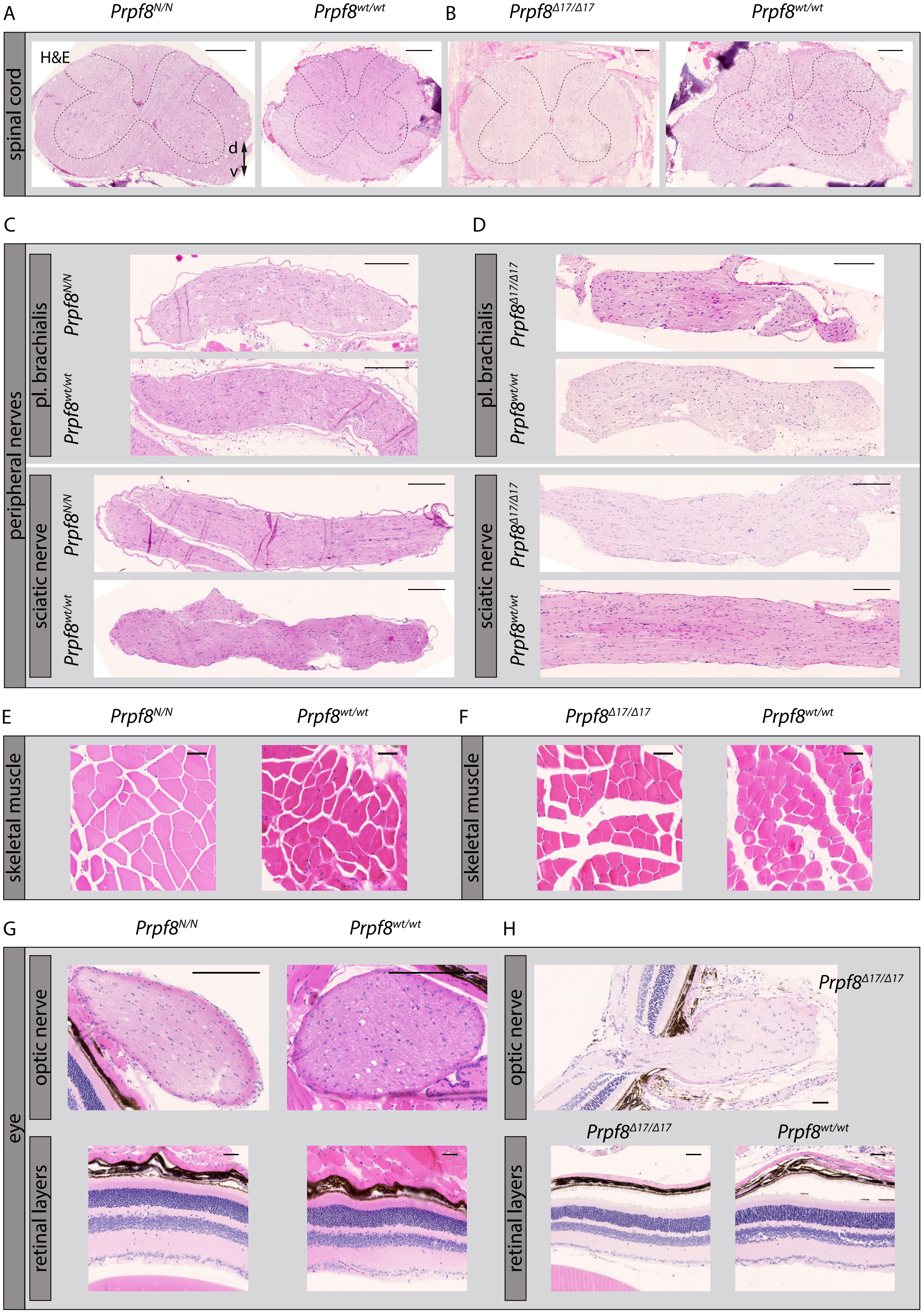
A
$Prpf8^{N/N}$
$Prpf8^{wt/wt}$
B
$Prpf8^{\Delta17/\Delta17}$
$Prpf8^{wt/wt}$
spinal cord
H&E
d
v
C
D
peripheral nerves
pl. brachialis
sciatic nerve
$Prpf8^{N/N}$
$Prpf8^{wt/wt}$
$Prpf8^{\Delta17/\Delta17}$
E
F
skeletal muscle
G
H
eye
optic nerve
retinal layers

**Figure S5. Pilot histopathological examination of assorted tissues in 22 weeks old homozygous *Prpf8*$^{Y2334N/Y2334N}$ and *Prpf8*$^{\Delta 17/\Delta 17}$ mice and corresponding wt littermates.** Investigated were specimen of spinal cord from *Prpf8*$^{Y2334N/Y2334N}$ (*A*) and *Prpf8*$^{\Delta 17/\Delta 17}$ (*B*) mice, moreover peripheral nerves (*C*, *D*), skeletal muscle (*E*, *F*), and eye (*G*, *H*). In contrast to the Prpf8 founder males, no gross pathologies were observed in any of the tissues analyzed. H&E = hematoxylin and eosin staining, d - dorsal, v - ventral. Scale bar = 200 µm in (*A*-*D*), and 50 µm in (*E*-*H*).

Figure S6

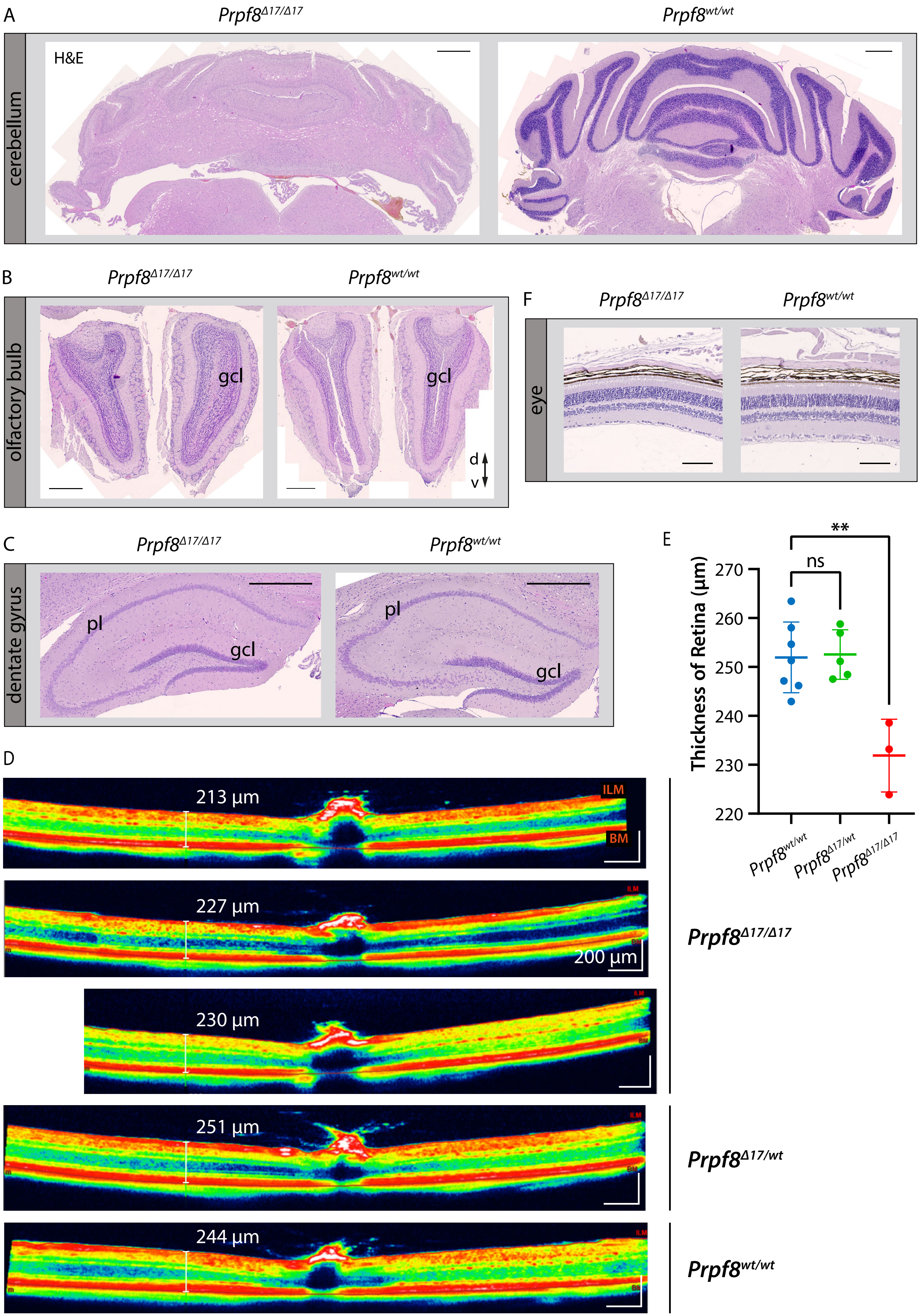

A
$Prpf8^{\Delta17/\Delta17}$
$Prpf8^{wt/wt}$
cerebellum
H&E
B
$Prpf8^{\Delta17/\Delta17}$
$Prpf8^{wt/wt}$
olfactory bulb
gcl
gcl
d
v
F
$Prpf8^{\Delta17/\Delta17}$
$Prpf8^{wt/wt}$
eye
C
$Prpf8^{\Delta17/\Delta17}$
$Prpf8^{wt/wt}$
dentate gyrus
pl
gcl
pl
gcl
E
**
ns
Thickness of Retina (µm)
270
260
250
240
230
220
$Prpf8^{wt/wt}$
$Prpf8^{\Delta17/wt}$
$Prpf8^{\Delta17/\Delta17}$
D
213 µm
ILM
BM
227 µm
200 µm
230 µm
251 µm
244 µm
$Prpf8^{\Delta17/\Delta17}$
$Prpf8^{\Delta17/wt}$
$Prpf8^{wt/wt}$

**Figure S6. Histopathological inspection of condition of selected granule cell populations in the brain and eyes in the cohort of 22 weeks old mice of the *Prpf8*$^{\Delta17}$ strain.** (*A*-*C*) While the cerebellar granule cell layer (gcl) was severely decreased (*A*), granular layers in the olfactory bulb (*B*), and in the dentate gyrus in hippocampus (*C*) did not show any signs of reduction when compared to the corresponding structure in a wt littermate. H&E = hematoxylin and eosin staining, gcl - granule cell layer, pl - pyramidal layer; d - dorsal, v – ventral. Scale bar = 500 µm. (*D*) Representative view of a cross-sectional image of retina gained by OCT examination. Retinal boundaries are indicated by red lines: ILM - internal limiting membrane (the boundary between the retina and vitreous body), BM - Bruch´s membrane (the innermost layer of the choroid). (*E*) Evaluation of the retinal thickness in 22 weeks old cohorts of *Prpf8*$^{\Delta17}$ animals revealed a decrease in retinal breadth in homozygous mice (1-way ANOVA followed by Dunnett's multiple comparison, p = 0.0017, **), while no difference was observed between heterozygous and control mice (ns). n=7 for *Prpf8*$^{wt/wt}$ animals, n=5 for *Prpf8*$^{\Delta17/wt}$, and n=3 for *Prpf8*$^{\Delta17/\Delta17}$ mice; all genotype conditions included animals of both sexes. (*F*) A representative image displaying preserved retinal layers in a homozygous *Prpf8*$^{\Delta17/\Delta17}$ mouse and a control wt littermate. Scale bar = 100 µm.

Figure S7

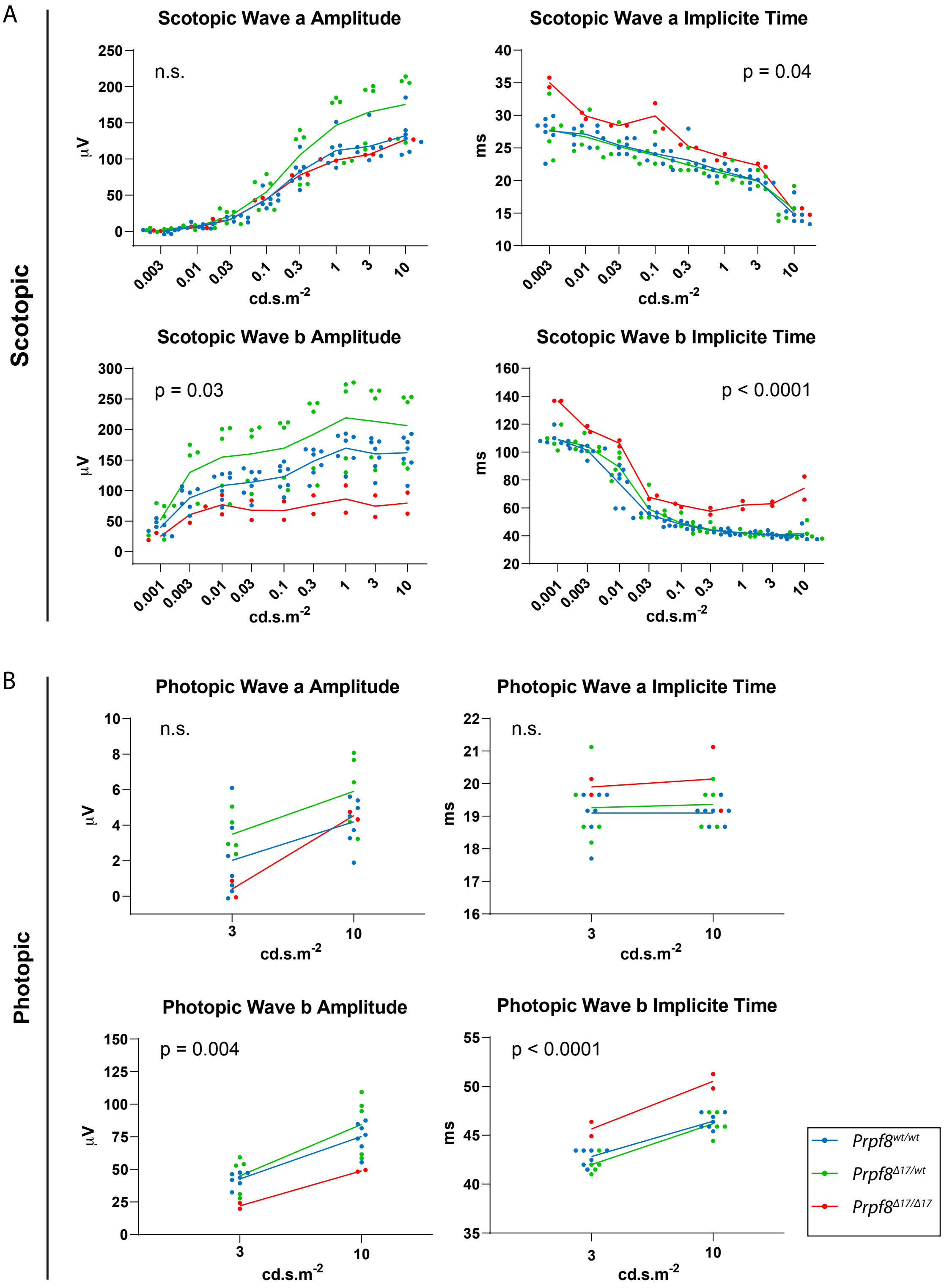


**Figure S7. Scotopic and photopic electroretinography (ERG) examination of 22 weeks old mice of the Prpf8Δ17 strain. Scotopic** (A) and photopic (B) ERG measurements. Luminance-response plots of full-field single-flash ERG for wave a are displayed in the top row, and for wave b in the second row. Values of amplitude and implicite time, respectively, were averaged between left and right eye of each animal. Dots represent individual animals, group averages of wild type, heterozygous and homozygous mutants are shown as connecting lines. P-values show a result of 2-way ANOVA for the difference among groups. n=5 for Prpf8wt/wt animals, n=7 for Prpf8Δ17/wt, and n=3 for Prpf8Δ17/Δ17 mice; results of one homozygous animal were removed from the analysis by outlier exclusion.

Figure S8

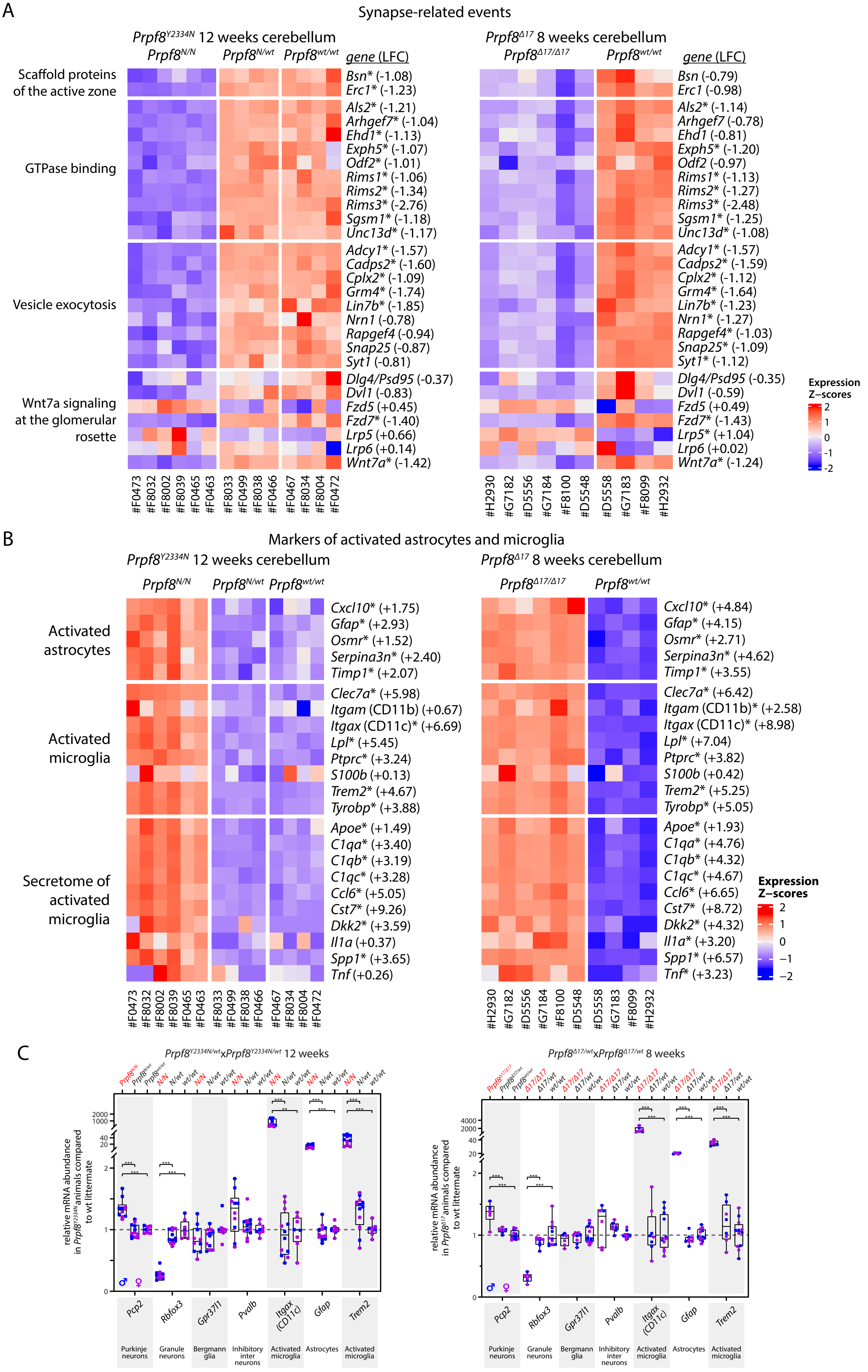
A
Synapse-related events
Prpf8Y2334N 12 weeks cerebellum
Prpf8N/N
Prpf8N/wt
Prpf8wt/wt
gene (LFC)
Prpf8Δ17 8 weeks cerebellum
Prpf8Δ17/Δ17
Prpf8wt/wt
Scaffold proteins of the active zone
GTPase binding
Vesicle exocytosis
Wnt7a signaling at the glomerular rosette
Bsn* (-1.08)
Erc1* (-1.23)
Als2* (-1.21)
Arhgef7* (-1.04)
Ehd1* (-1.13)
Exph5* (-1.07)
Odf2* (-1.01)
Rims1* (-1.06)
Rims2* (-1.34)
Rims3* (-2.76)
Sgsm1* (-1.18)
Unc13d* (-1.17)
Adcy1* (-1.57)
Cadps2* (-1.60)
Cplx2* (-1.09)
Grm4* (-1.74)
Lin7b* (-1.85)
Nrn1 (-0.78)
Rapgef4 (-0.94)
Snap25 (-0.87)
Syt1 (-0.81)
Dlg4/Psd95 (-0.37)
Dvl1 (-0.83)
Fzd5 (+0.45)
Fzd7* (-1.40)
Lrp5 (+0.66)
Lrp6 (+0.14)
Wnt7a* (-1.42)
Bsn (-0.79)
Erc1 (-0.98)
Als2* (-1.14)
Arhgef7 (-0.78)
Ehd1 (-0.81)
Exph5* (-1.20)
Odf2 (-0.97)
Rims1* (-1.13)
Rims2* (-1.27)
Rims3* (-2.48)
Sgsm1* (-1.25)
Unc13d* (-1.08)
Adcy1* (-1.57)
Cadps2* (-1.59)
Cplx2* (-1.12)
Grm4* (-1.64)
Lin7b* (-1.23)
Nrn1* (-1.27)
Rapgef4* (-1.03)
Snap25* (-1.09)
Syt1* (-1.12)
Dlg4/Psd95 (-0.35)
Dvl1 (-0.59)
Fzd5 (+0.49)
Fzd7* (-1.43)
Lrp5* (+1.04)
Lrp6 (+0.02)
Wnt7a* (-1.24)
Expression Z-scores
2
1
0
-1
-2
#F0473 #F8032 #F8002 #F8039 #F0465 #F0463 #F8033 #F0499 #F8038 #F0466 #F0467 #F8034 #F8004 #F0472
#H2930 #G7182 #D5556 #G7184 #F8100 #D5548 #D5558 #G7183 #F8099 #H2932
B
Markers of activated astrocytes and microglia
Prpf8Y2334N 12 weeks cerebellum
Prpf8N/N
Prpf8N/wt
Prpf8wt/wt
Prpf8Δ17 8 weeks cerebellum
Prpf8Δ17/Δ17
Prpf8wt/wt
Activated astrocytes
Activated microglia
Secretome of activated microglia
Cxcl10* (+1.75)
Gfap* (+2.93)
Osmr* (+1.52)
Serpina3n* (+2.40)
Timp1* (+2.07)
Clec7a* (+5.98)
Itgam (CD11b) (+0.67)
Itgax (CD11c)* (+6.69)
Lpl* (+5.45)
Ptprc* (+3.24)
S100b (+0.13)
Trem2* (+4.67)
Tyrobp* (+3.88)
Apoe* (+1.49)
C1qa* (+3.40)
C1qb* (+3.19)
C1qc* (+3.28)
Ccl6* (+5.05)
Cst7* (+9.26)
Dkk2* (+3.59)
Il1a (+0.37)
Spp1* (+3.65)
Tnf (+0.26)
Cxcl10* (+4.84)
Gfap* (+4.15)
Osmr* (+2.71)
Serpina3n* (+4.62)
Timp1* (+3.55)
Clec7a* (+6.42)
Itgam (CD11b)* (+2.58)
Itgax (CD11c)* (+8.98)
Lpl* (+7.04)
Ptprc* (+3.82)
S100b (+0.42)
Trem2* (+5.25)
Tyrobp* (+5.05)
Apoe* (+1.93)
C1qa* (+4.76)
C1qb* (+4.32)
C1qc* (+4.67)
Ccl6* (+6.65)
Cst7* (+8.72)
Dkk2* (+4.32)
Il1a* (+3.20)
Spp1* (+6.57)
Tnf* (+3.23)
Expression Z-scores
2
1
0
−1
−2
#F0473 #F8032 #F8002 #F8039 #F0465 #F0463 #F8033 #F0499 #F8038 #F0466 #F0467 #F8034 #F8004 #F0472
#H2930 #G7182 #D5556 #G7184 #F8100 #D5548 #D5558 #G7183 #F8099 #H2932
C
Prpf8Y2334N/wtxPrpf8Y2334N/wt 12 weeks
Prpf8Δ17/wtxPrpf8Δ17/wt 8 weeks
relative mRNA abundance in Prpf8Y2334N animals compared to wt littermate
relative mRNA abundance in Prpf8Δ17 animals compared to wt littermate
Pcp2
Rbfox3
Gpr37l1
Pvalb
Itgax (CD11c)
Gfap
Trem2
Purkinje neurons
Granule neurons
Bergmann glia
Inhibitory inter neurons
Activated microglia
Astrocytes
Activated microglia

**Figure S8. Differential gene expression in cerebellum of 12-weeks-old *Prpf8*$^{Y2334N}$ animals and 8-weeks-old *Prpf8*$^{\Delta 17}$ mice.** (*A*) Heatmap representation of expression levels of selected genes in the RNA-Seq data from categories related to synaptic structure, events, and Wnt signaling. (*B*) Heatmap representation of gene expression levels of groups of selected genes that represent transcriptional signature of activated astrocytes and microglia. Values were standardized to display the same range of expression values for each gene. * - differentially expressed genes defined by $|\text{Log}_2$ Fold change$|>1$ and FDR<0.05; the estimated $\text{Log}_2$ Fold change value (LFC) is provided for individual genes and represents the difference between homozygous and wt animals. (*C*) Relative mRNA abundance of assorted marker genes representing homeostatic cerebellar subtypes and degeneration-activated populations in cerebellar samples from 12-weeks-old *Prpf8*$^{Y2334N}$ animals *(left panel)* and in 8-weeks-old *Prpf8*$^{\Delta 17}$ mice (*right panel*). The fold change values are plotted in Box-and-whiskers diagram with indicated position of median; statistical significance of differential expression in the given genotype groups was examined by t-test using GraphPad software. ** $P < 0.01$, *** $P < 0.001$.

Figure S9

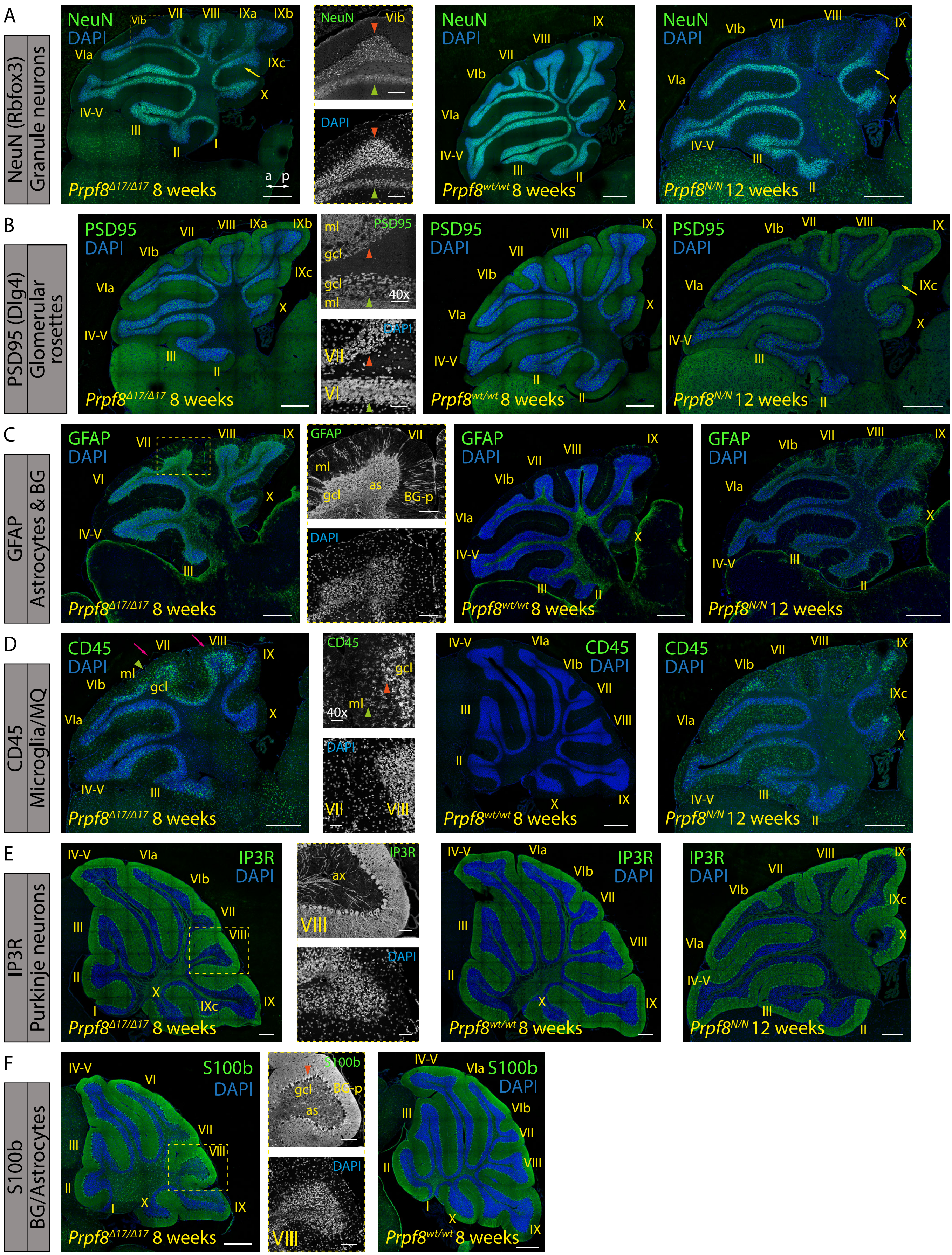
A
NeuN (Rbfox3)
Granule neurons
NeuN
DAPI
Prpf8Δ17/Δ17 8 weeks
Prpf8wt/wt 8 weeks
Prpf8N/N 12 weeks
B
PSD95 (Dlg4)
Glomerular rosettes
PSD95
DAPI
40x
C
GFAP
Astrocytes & BG
GFAP
DAPI
ml
gcl
as
BG-p
D
CD45
Microglia/MΘ
CD45
DAPI
40x
E
IP3R
Purkinje neurons
IP3R
DAPI
ax
F
S100b
BG/Astrocytes
S100b
DAPI

**Figure S9. Immunohistochemical staining for markers of selected cerebellar cell types in 8 weeks old *Prpf8*$^{\Delta17/\Delta17}$ and 12 weeks old *Prpf8*$^{Y2334N/Y2334N}$ mice.** (*A*) Rbfox3 (NeuN) immunolabeling was used to visualize nuclei of granule neurons. The detail shows an apoptotic zone in apical portion of lobule VIb (orange arrowhead) in contrast to its less affected vermian portion (green arrowhead). Boundary between posterior lobule IXb and lobule IVc indicated by the yellow arrow. a - anterior, p - posterior. (*B*) The punctate pattern of PSD95 marks glomerular rosette in the granular layer and synaptic junctions of Purkinje dendrites. Detail: observed decay of the glomeruli (orange arrowhead), the PSD95 signal within the molecular cell layer (ml) was preserved. A retained, vermian portion of lobule VI is shown for comparison (green arrowhead). Boundary between posterior lobule IXb and lobule IVc indicated by the yellow arrow. ml - molecular layer, gcl - granule cell layer. (*C*) The signal of GFAP was enhanced in the granule cell layer (gcl) of homozygous animals, confirming activation of astrocytes (as). The signal in the molecular layer (ml) represents GFAP-positive processes of Bergmann glia (BG-p). (*D*) CD45 immunostaining visualized phagocytic microglia and/or macrophages in the granule cell layer (gcl) of the degenerating posterior lobe lobules (pink arrows). The labelled microglia showed amoeboid morphology with thick cellular processes characteristic of DAMs (orange arrowhead). The signal was also present in the molecular layer (ml), which might coincide with clearance of residual parallel fibers from degenerated granule neurons (green arrowheads). (*E*) Purkinje cells were visualized based on expression of IP3R; that is present both in the somatodendritic compartment as well as in the axonal projections (ax). No major perturbance of Purkinje neurons was spotted even in the apoptotic areas (detail of lobule VIII). (*F*) The cell bodies of Bergmann glial cells were normally localized around Purkinje cell somata (orange arrowhead) and their radial processes (BG-p) properly terminated with endfeet at the pial surface, as revealed by staining with anti S100 protein. The anti S100b antibody in parallel visualized activated astrocytes (as) in the granule cell layer (gcl). (*A*-*F*) Confocal immunofluorescence images of 5 µm FFPE sagittal cerebellar slices, scale bar = 200 µm, in insets 50 µm. Only the *Prpf8*$^{wt/wt}$ littermates of *Prpf8*$^{\Delta17/\Delta17}$ animals are shown for clarity.

Figure S10

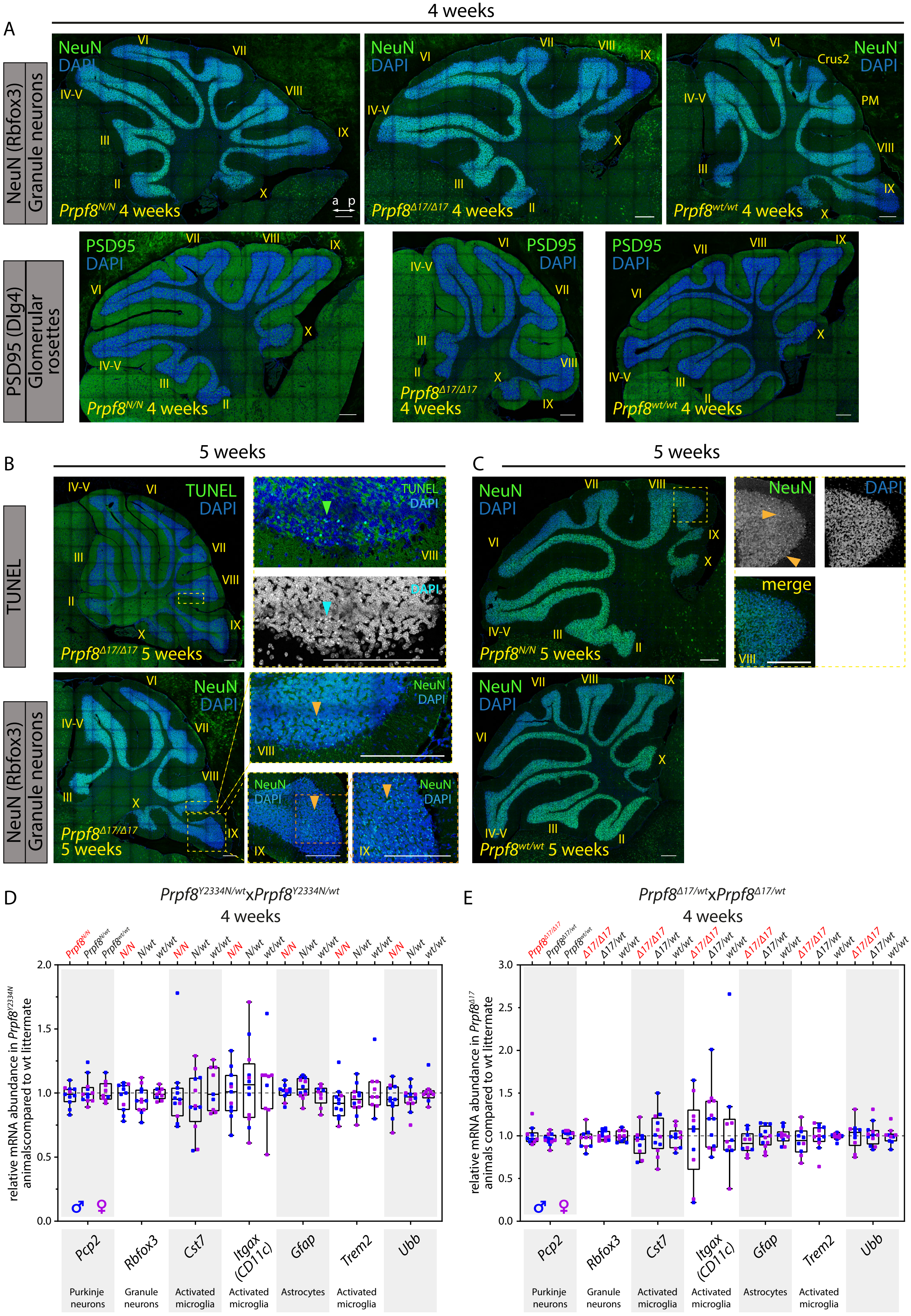
A
4 weeks
NeuN (Rbfox3) Granule neurons
PSD95 (Dlg4) Glomerular rosettes
NeuN
DAPI
PSD95
Prpf8N/N 4 weeks
Prpf8Δ17/Δ17 4 weeks
Prpf8wt/wt 4 weeks
Crus2
PM
B
5 weeks
TUNEL
Prpf8Δ17/Δ17 5 weeks
C
5 weeks
merge
Prpf8N/N 5 weeks
Prpf8wt/wt 5 weeks
D
Prpf8Y2334N/wtxPrpf8Y2334N/wt
4 weeks
relative mRNA abundance in Prpf8Y2334N animalscompared to wt littermate
E
Prpf8Δ17/wtxPrpf8Δ17/wt
4 weeks
relative mRNA abundance in Prpf8Δ17 animals compared to wt littermate
Pcp2
Rbfox3
Cst7
Itgax (CD11c)
Gfap
Trem2
Ubb
Purkinje neurons
Granule neurons
Activated microglia
Activated microglia
Astrocytes
Activated microglia

**Figure S10. Immunohistochemical staining and qPCR analyses in cerebellar samples harvested from 4-5 weeks old *Prpf8*$^{Y2334N/Y2334N}$ mice and *Prpf8*$^{\Delta17/\Delta17}$.** (*A*) No differences between the genotypes at 4 weeks of age were recorded in the immunolabeling pattern of Rbfox3/NeuN (top row) and PSD95 (bottom row). a - anterior, p - posterior, Crus2 - crus 2 of the ansiform lobule, PM - paramedian lobule. (*B-C*) Immunohistochemical staining and TUNEL assay performed with samples collected from 5 weeks old *Prpf8*$^{\Delta17}$ animals (*B*) and *Prpf8*$^{Y2334N}$ mice (*C*). (*B*) After reaching week 5, the apoptotic signal (TUNEL assay, top row, green arrowhead) started to appear in apical portions of posterior lobe lobules (provided detail is from lobule VIII). This pattern coincided with emergence of shrunken granule cell visualized by DAPI (blue arrowhead) and by the anti-Rbfox3 (NeuN) antibody (orange arrowheads, bottom row). (*C*) Staining of shrunken, likely apoptotic cells by the anti-Rbfox3 (NeuN) antibody in apical portions of posterior lobe in *Prpf8*$^{Y2334N/Y2334N}$ mice, detail is from lobule VIII. Confocal immunofluorescence images of 5 µm FFPE sagittal cerebellar slices, scale bar = 200 µm. Only the *Prpf8*$^{wt/wt}$ littermates of *Prpf8*$^{N/N}$ animals are shown for clarity in (*A*). (*D-E*) Quantitative RT-qPCR analysis performed with cerebellar samples harvested form 4 weeks old *Prpf8*$^{Y2334N}$ (*D*), and *Prpf8*$^{\Delta17}$ mice (*E*) to assay the abundance of homeostatic and activated cerebellar cell populations. The amounts of total RNA in individual samples were normalized to *Gapdh* expression, and relative mRNA abundance was then calculated as fold change (FC) difference of homozygous or heterozygous animals to *Prpf8*$^{wt/wt}$ controls using the $2^{\wedge\Delta\Delta Ct}$ approach. The results for an additional housekeeping gene ubiquitin B (*Ubb*) are shown. The higher variance in the RNA content of activated microglia markers *Cst-7* and *Itgax11* is likely caused by their physiological scarcity in healthy tissue (Ct>30). Total number of analyzed animals was as follows: 1. 4-weeks old *Prpf8*$^{Y2334N}$ mice: *Prpf8*$^{Y2334N/Y2334N}$ n=11, *Prpf8*$^{Y2334N/wt}$ n=12, *Prpf8*$^{wt/wt}$ n=9. 2. 4-weeks old *Prpf8*$^{\Delta17}$ animals: *Prpf8*$^{\Delta17/\Delta17}$ n=10, *Prpf8*$^{\Delta17/wt}$ n=12, *Prpf8*$^{wt/wt}$ n=10. All genetic conditions contained animals of both genders (samples harvested from males are depicted in blue, females in pink).

Figure S11

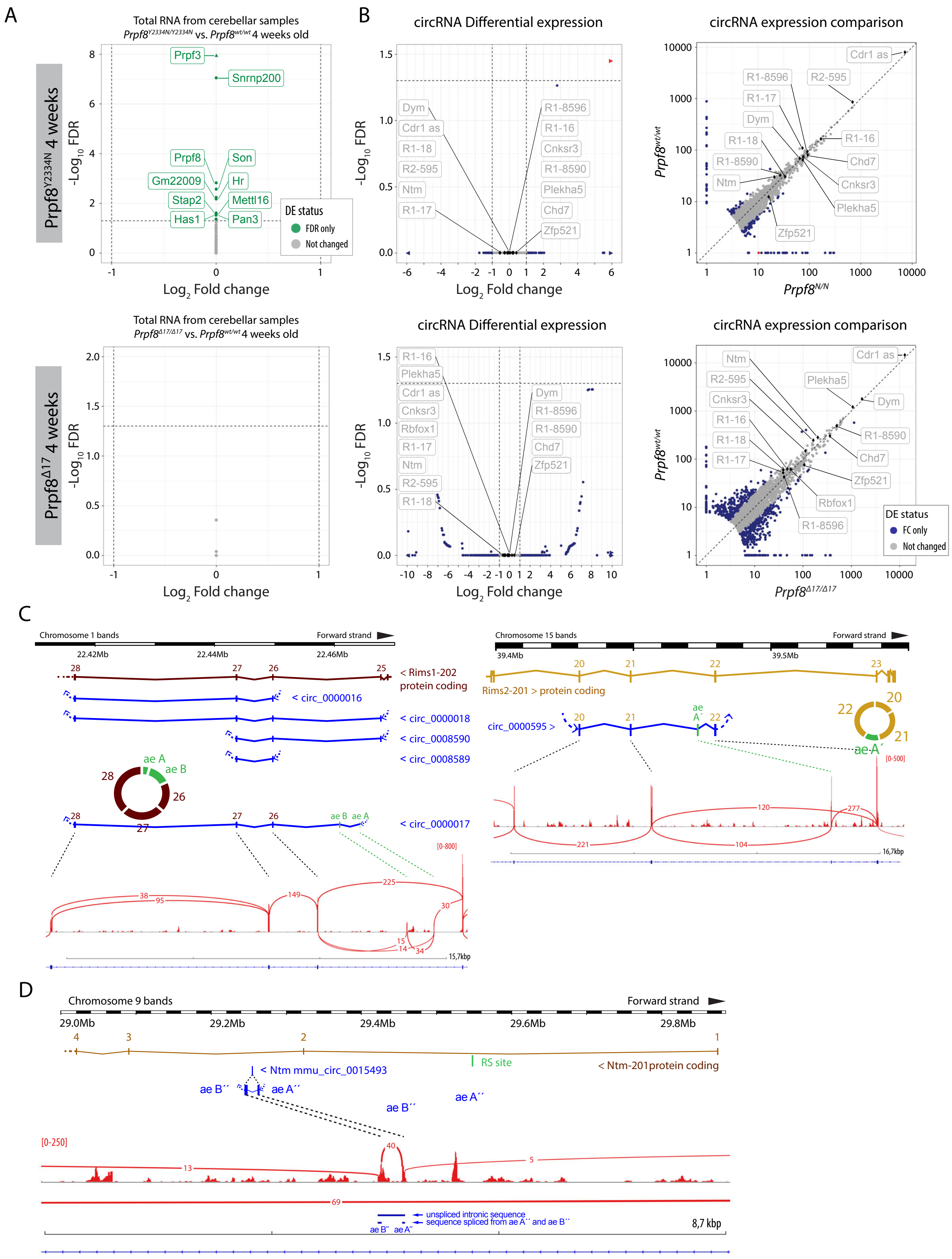
A
Total RNA from cerebellar samples
Prpf8Y2334N/Y2334N vs. Prpf8wt/wt 4 weeks old
Prpf8Y2334N 4 weeks
-Log10 FDR
Log2 Fold change
Prpf3
Snrnp200
Prpf8
Son
Gm22009
Hr
Stap2
Mettl16
Has1
Pan3
DE status
FDR only
Not changed
B
circRNA Differential expression
circRNA expression comparison
Dym
Cdr1 as
R1-18
R2-595
Ntm
R1-17
R1-8596
R1-16
Cnksr3
R1-8590
Plekha5
Chd7
Zfp521
Prpf8wt/wt
Prpf8N/N
Total RNA from cerebellar samples
Prpf8Δ17/Δ17 vs. Prpf8wt/wt 4 weeks old
Prpf8Δ17 4 weeks
Rbfox1
Prpf8Δ17/Δ17
DE status
FC only
Not changed
C
Chromosome 1 bands
Forward strand
22.42Mb
22.44Mb
22.46Mb
< Rims1-202 protein coding
< circ_0000016
< circ_0000018
< circ_0008590
< circ_0008589
< circ_0000017
ae A
ae B
[0-800]
15,7kbp
Chromosome 15 bands
Forward strand
39.4Mb
39.5Mb
Rims2-201 > protein coding
circ_0000595 >
ae A´
[0-500]
16,7kbp
D
Chromosome 9 bands
Forward strand
29.0Mb
29.2Mb
29.4Mb
29.6Mb
29.8Mb
RS site
< Ntm-201protein coding
< Ntm mmu_circ_0015493
ae B´´
ae A´´
[0-250]
unspliced intronic sequence
sequence spliced from ae A´´ and ae B´´
8,7 kbp

**Figure S11. Transcriptomic analyses of 4 weeks old cerebella of both Prpf8 mutant strains.** (*A*) Differential gene expression analyses (DESeq2) of total cerebellar RNA revealed no significant differences in linear transcription between 4 weeks old *Prpf8*$^{Y2334N/Y2334N}$ (top panel) and *Prpf8*$^{\Delta17/\Delta17}$ (bottom panel) animals and their wt littermates. Genes that were differentially expressed in *Prpf8*$^{Y2334N/Y2334N}$ animals (FDR<0.05, but did not pass the $|\text{Log}_2$ Fold change$|>1$ criterion) are listed in Supplementary Table 7. (*B*) Volcano and scatter plots showing circRNA expression between homozygous Prpf8 mutant cerebella and wt controls. Scatterplot comparing library-normalized abundance of individual circRNAs (in BSJ read counts) between homozygous mutant mice (on the X-axis) and wt controls (Y-axis). R1 = Rims1; R2 = Rims2. Accompanying data can be found in Supplementary Table 2. Assorted circRNAs are highlighted by diamond symbols and their specifications are provided in Table S8. Triangles represent circRNAs with values reaching beyond the displayed scale. Only circRNAs with mean normalized BSJ read counts minimum of 3 are shown in all plots. (*C-D*) Scheme of segments from *Rims1*, *Rims2 (C)* and *Ntm* genes *(D)* that give rise to the studied circRNA species.

Figure S12

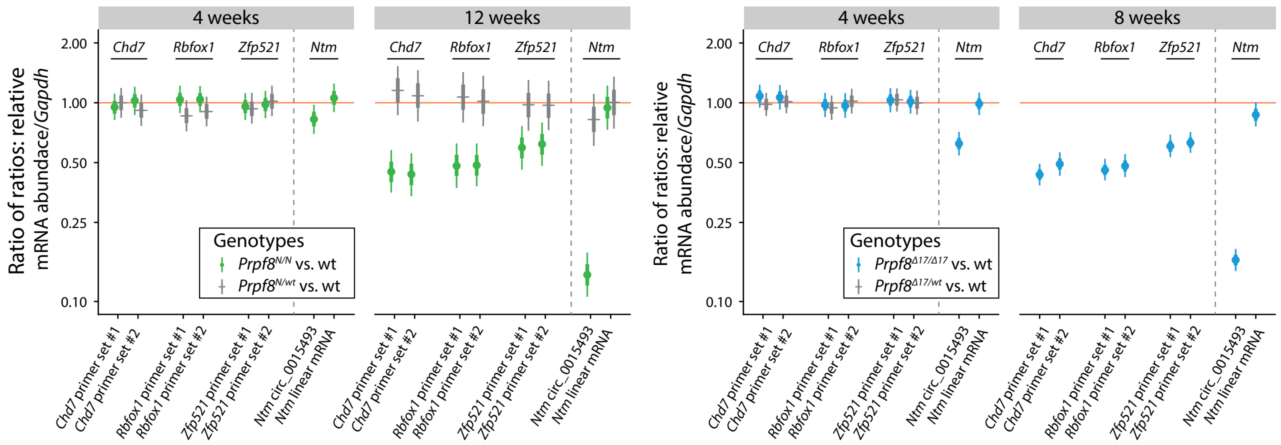


**Figure S12. Expression of Rbfox1, Zfp521, Chd7, and Ntm host genes in the cerebellar tissue of homozygous Prpf8 mutant mice.** Statistical model-based estimates of abundance of linear transcripts produced from Rbfox1, Zfp521, Chd7, and Ntm genes (for Ntm also circular RNA abundance is displayed on the left side of panels) relative to Gapdh expression in cerebella gained from 4- and 12 weeks old Prpf8Y2334N animals (left panel), and from 4- and 8 weeks old Prpf8Δ17 mice (right panel). The estimates are based on a linear mixed model of RT-qPCR data accounting for differences between sexes, litters, qPCR runs and individual animals. Posterior distribution of ratio of each transcript (horizontal axis) to Gapdh was computed for each genotype. Estimates of the ratio of those ratios are shown (vertical axis, log scale). Shown are posterior credible intervals (lines: 95% - thin, 50% - thick), and means (points). Asterisks indicate comparisons for which the 95% credible interval excludes 1 (no difference). Abundance of selected mRNAs (Rbfox1, Zfp521, and Chd7 genes) was probed using two independent sets of primers (#1 and #2).

Figure S13

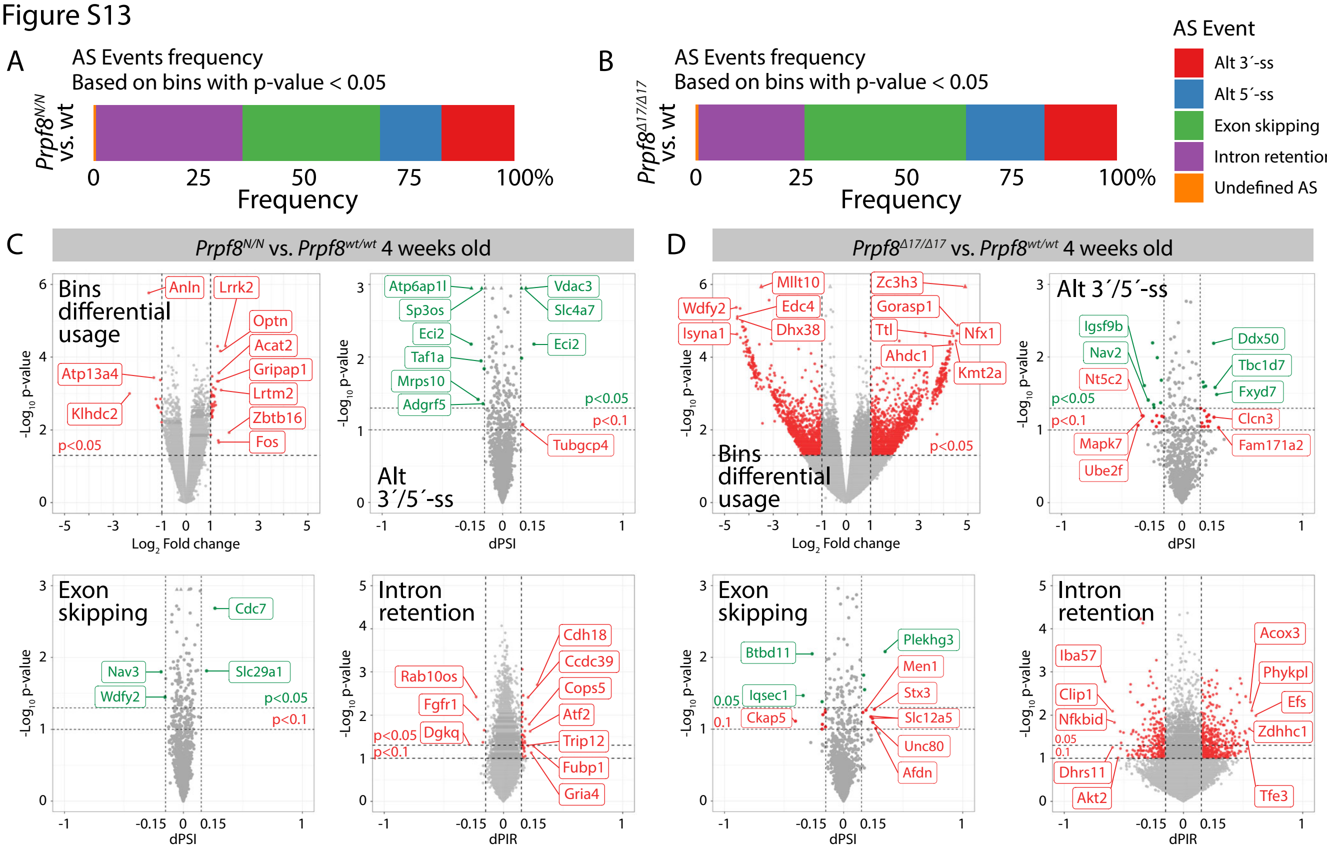


**Figure S13. Analysis of alternative splicing events and circular RNAs in 4 weeks old cerebella collected from Prpf8Y2334N and Prpf8Δ17 animals.** (A-D) ASpli analysis of alternative splicing events in 4 weeks old cerebella collected from Prpf8Y2334N (A, C) and Prpf8Δ17 (B, D) animals vs. wt controls. Transcripts were analyzed for differential bins (exons, introns, and alternative 5´- and 3´- splice sites (ss)) usage based on their expression (overlapping fragment counts). Furthermore, intron retention dPIR index, and exon skipping and alternative 5´- and 3´-splice sites dPSI indexes were calculated based on event inclusion/exclusion supporting fragment counts. dPIR/dPSI p-values were calculated using Welch´s t-test. For AS event frequency summarization, only differentially used bins (p-value < 0.05) were considered. Reported splicing changes were considered significant in case they had a p-value < 0.05 and change in inclusion level difference (dPIR or dPSI) of more than 15%. Triangles represent spicing events changes with values reaching beyond the displayed scale.

Figure S14

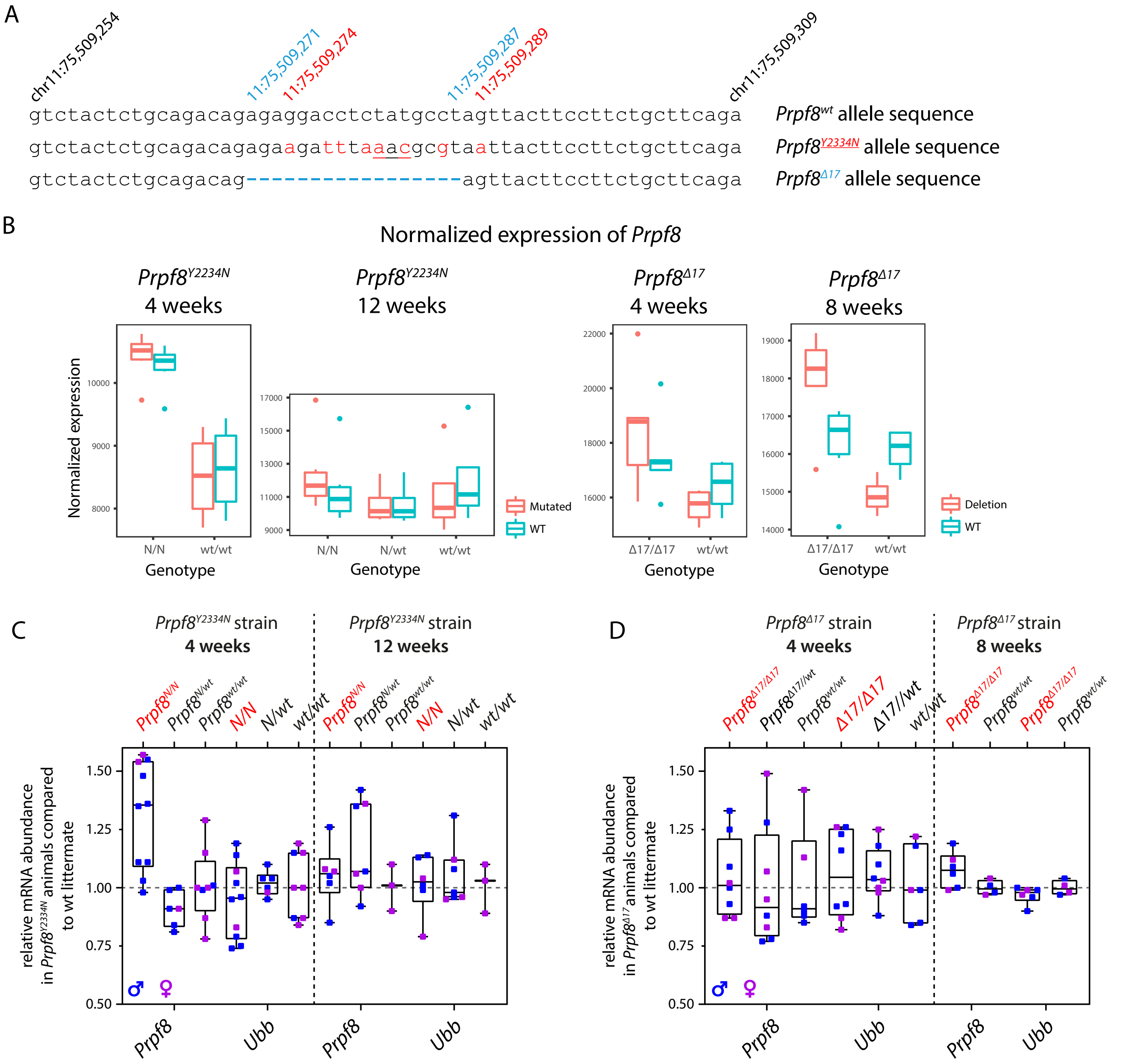


**Figure S14. Abundance of Prpf8 transcripts in cerebellar samples from Prpf8Y2334N and Prpf8Δ17 animals.** (A) To accurately quantify the expression originating from the Prpf8wt, Prpf8Y2334N, and Prpf8Δ17 alleles in the RNA-Seq data, Prpf8 reads were re-mapped to modified versions of chromosome 11 that contained the introduced genomic alterations. (B) Normalized expression of Prpf8 in the cerebellar tissue inferred from four RNA-Seq experiments. The Prpf8 RNA content seemed increased in homozygous animals of both Prpf8 mutant strains at 4 weeks, and in 8-weeks old Prpf8Δ17/Δ17 individuals when compared to Prpf8wt/wt littermates. (C-D) Quantitative RT-PCR analysis performed with cerebellar samples harvested form 4- and 12-weeks old Prpf8Y2334N mice (C), and from 4- and 8-weeks old Prpf8Δ17 animals (D). Total number of animals sacrificed at indicated timepoints was as follows: 1. 4-weeks old Prpf8Y2334N mice: Prpf8Y2334N/Y2334N n=10, Prpf8Y2334N/wt n=6, Prpf8wt/wt n=8. 2. 12-weeks old Prpf8Y2334N mice: Prpf8Y2334N/Y2334N n=6, Prpf8Y2334N/wt n=7, Prpf8wt/wt n=3. 3. 4-weeks old Prpf8Δ17 animals: Prpf8Δ17/Δ17 n=8, Prpf8Δ17/wt n=8, Prpf8wt/wt n=6. 4. 8-weeks old Prpf8Δ17 animals: Prpf8Δ17/Δ17 n=6, Prpf8wt/wt n=4. All genetic conditions contained animals of both genders (samples harvested from males are depicted in blue, females in pink). The amounts of total RNA in individual samples were normalized to Gapdh expression, and relative mRNA abundance was then calculated as fold change (FC) difference of homozygous or heterozygous animals to Prpf8wt/wt controls using the 2^ΔΔCt approach. The results for additional housekeeping gene ubiquitin B (Ubb) are shown. Similar to the RNA-Seq cohorts the Prpf8 levels appeared increased in 4-weeks old Prpf8Y2334N/Y2334N, this finding was however not significant as examined by t-test.

Figure S15

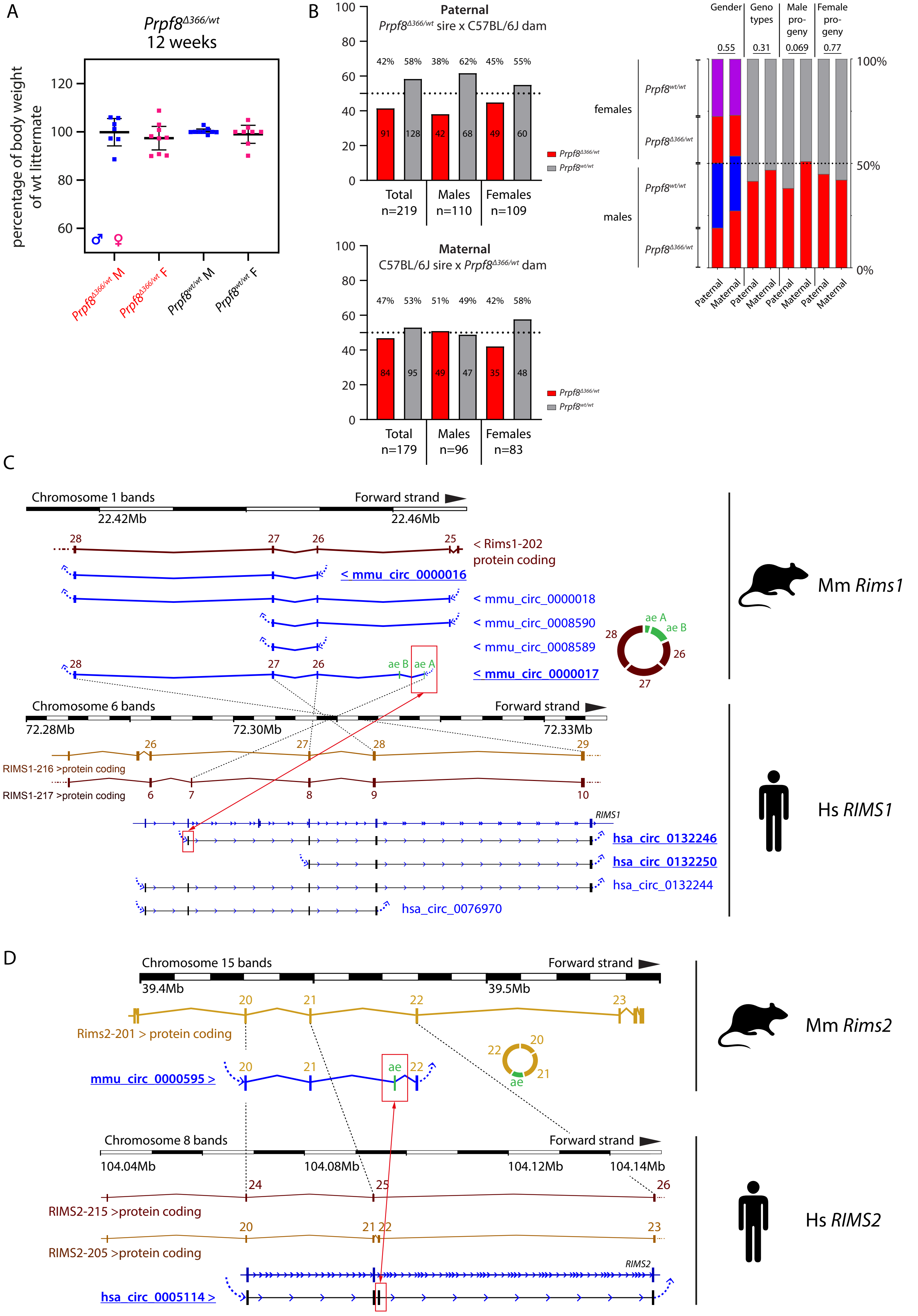
A
Prprf8Δ366/wt
12 weeks
percentage of body weight of wt littermate
120
100
80
60
♂ ♀
Prpf8Δ366/wt M
Prpf8Δ366/wt F
Prpf8wt/wt M
Prpf8wt/wt F
B
Paternal
Prpf8Δ366/wt sire x C57BL/6J dam
42% 58% 38% 62% 45% 55%
91 128 42 68 49 60
Total n=219
Males n=110
Females n=109
Prpf8Δ366/wt
Prpf8wt/wt
Maternal
C57BL/6J sire x Prpf8Δ366/wt dam
47% 53% 51% 49% 42% 58%
84 95 49 47 35 48
Total n=179
Males n=96
Females n=83
Gender
Geno types
Male pro- geny
Female pro- geny
0.55
0.31
0.069
0.77
100%
50%
0%
females
males
Prpf8wt/wt
Prpf8Δ366/wt
Paternal
Maternal
C
Chromosome 1 bands
Forward strand
22.42Mb
22.46Mb
28 27 26 25
< Rims1-202 protein coding
< mmu_circ_0000016
< mmu_circ_0000018
< mmu_circ_0008590
< mmu_circ_0008589
< mmu_circ_0000017
ae B ae A
Mm Rims1
Chromosome 6 bands
72.28Mb
72.30Mb
72.33Mb
26 27 28 29
RIMS1-216 >protein coding
RIMS1-217 >protein coding
6 7 8 9 10
RIMS1
hsa_circ_0132246
hsa_circ_0132250
hsa_circ_0132244
hsa_circ_0076970
Hs RIMS1
D
Chromosome 15 bands
Forward strand
39.4Mb
39.5Mb
20 21 22 23
Rims2-201 > protein coding
mmu_circ_0000595 >
ae
Mm Rims2
Chromosome 8 bands
Forward strand
104.04Mb
104.08Mb
104.12Mb
104.14Mb
24 25 26
RIMS2-215 >protein coding
20 21 22 23
RIMS2-205 >protein coding
RIMS2
hsa_circ_0005114 >
Hs RIMS2

**Figure S15. Analysis of 12-weeks old *Prpf8*$^{\Delta 366}$ mice and comparison of mouse and human circRNAs formed from *Rims2*/*RIMS2* and *Rims1*/*RIMS1* genes**. (*A-B*) Analysis of 12-weeks old *Prpf8*$^{\Delta 366}$ mice. (*A*) Body weight measurements of 12-weeks old *Prpf8*$^{\Delta 366/wt}$ animals and their wt counterparts did not reveal any significant differences between the two genotypes. n=7 for *Prpf8*$^{\Delta 366/wt}$ males, n=9 for *Prpf8*$^{\Delta 366/wt}$ females, and n=9 for the corresponding *Prpf8*$^{wt/wt}$ males, n=8 for *Prpf8*$^{wt/wt}$ females. Actual body weight of animals was normalized to median of *Prpf8*$^{wt/wt}$ peers of matching gender in individual litter. Displayed is mean with 95% confidentiality interval; males (M) in blue, females (F) in pink. (*B*) Analysis of *Prpf8*$^{\Delta 366/wt}$ breeding records. Offspring numbers are provided separately for the paternal and maternal transmission route, and then in a merged chart. Genotype and sex ratios were not skewed (Fischer´s exact test; p=0.31 and p=0.55, respectively), and there was as well no evidence for maternal transmission ratio distortion (p=0.77). We however noted a possible tendency for paternal transmission ratio distortion (p=0.069). (*C-D*) Comparison of mouse and human circRNAs formed from *Rims2*/*RIMS2* and *Rims1*/*RIMS1* genes. CircRNAs produced from *Rims1*/*RIMS1* and *Rims2*/*RIMS2* genes are evolutionary conserved between mice and human. (*C*) A scheme of the murine *Rims1* locus with portion of the *Rims1* transcript Rims1-202 (ENSMUST00000097808.9), and a spectrum of distinct circRNA species that arise from a region comprising *Rims1* exons 25-28. Below is provided a corresponding scheme of the human *RIMS1* gene, with a portion of transcript RIMS1-216 (ENST00000521978.5). Mouse *Rims1* circRNAs circ_0000017 and circ_0000016, and the corresponding human counterparts *RIMS1* circ_0132246 and circ_0132250, respectively, account for the most abundant circRNAs formed from *Rims1*/*RIMS1* gene in the cerebellum (underlined) (5). In human, the alternate exon located in intron 26-27, corresponding to the alternate exon A in *Rims1* circ_0000017, can be included in *RIMS1* protein-coding transcripts (RIMS1-217), and is part of all circRNA species formed from the locus. In contrast, in mouse *Rims1* the alternate exon A is not included in mRNAs, and is moreover excluded from circRNAs with excerption of circ_0000017 (red rectangles). (*D*) Scheme of the murine *Rims2* locus, with depicted 3´ part of the *Rims2* transcript Rims2-201 (ENSMUST00000082054.12). *Rims2* exons 20-22 including an alternate exon (ae) circularize to form circRNA mmu_circ_0000595. In human, the corresponding hsa_circ_0005114 is formed from *RIMS2* exons 20-23 (exon numbering according to transcript RIMS2-205, ENST00000436393.6), it however contains a different alternate exon when compared to mouse (red rectangles). Mouse *Rims2* circ_0000595 and human *RIMS2* circ_0005114 represent the most abundant circRNA formed from the *Rims2*/*RIMS2* genes in the cerebellum (5). The alternate exon located in intron 21-22 is in the mouse likely circRNA-specific, while in human the alternate exon can get included in mRNAs as well.
