## Supplementary tables for "Retinitis pigmentosa associated mutations in mouse Prpf8 cause misexpression of circRNAs and degeneration of cerebellar granule neurons": Supplementary_tables-legend.docx

**Table S1.** List of all primers used in the study.

**Table S2.** Analysis of differentially expressed genes by DESeq2 in all RNA-Seq runs.

**Table S3.** Analysis of alternative splicing in RNA-Seq runs from 12 weeks old *Prpf8^Y2334N^* mice and 8 weeks old *Prpf8^Δ17^* animals by ASpli.

**Table S4.** Ciri- and CiriQuant quantification underlying Volcano, scatter, and ratio plots in all RNA-Seq runs.

**Table S5.** Specifications of assorted circRNAs displayed in Volcano, scatter and ratio plots.

**Table S6**. Inclusion of alternate exons in *Ntm*, *Rims1* and *Rims2* genes into linear and circular RNA forms; quantification from RNA-Seq runs from 4 weeks old *Prpf8^Y2334N/Y2334N^* and *Prpf8^Δ17/Δ17^* animals.

**Table S7.** Scoring of splice site strengths of assorted *Rims1*, *Rims2*, and *Ntm* exons.

**Table S8.** Analysis of alternative splicing in RNA-Seq runs from 4 weeks old *Prpf8^Y2334N/Y2334N^* and *Prpf8^Δ17/Δ17^* animals by ASpli.

**Table S9**. GSEA analysis - overlap of GO terms enriched in genes exhibiting differences in alternative splicing of linear RNA forms (i.e. intron retention, exon skipping, and alternative 5´-/3´-ss events identified by |dPIR/dPSI|>0.15), and differentially expressed circRNA host genes defined by |Log_2_ FC|>1. The analysis was carried out from the RNA-Seq results of 4 weeks old *Prpf8^Δ17/Δ17^* and *Prpf8^Y2334N/Y2334N^* animals.
